# Community-based Reconstruction and Simulation of a Full-scale Model of Region CA1 of Rat Hippocampus

**DOI:** 10.1101/2023.05.17.541167

**Authors:** Armando Romani, Alberto Antonietti, Davide Bella, Julian Budd, Elisabetta Giacalone, Kerem Kurban, Sára Sáray, Marwan Abdellah, Alexis Arnaudon, Elvis Boci, Cristina Colangelo, Jean-Denis Courcol, Thomas Delemontex, András Ecker, Joanne Falck, Cyrille Favreau, Michael Gevaert, Juan B. Hernando, Joni Herttuainen, Genrich Ivaska, Lida Kanari, Anna-Kristin Kaufmann, James Gonzalo King, Pramod Kumbhar, Sigrun Lange, Huanxiang Lu, Carmen Alina Lupascu, Rosanna Migliore, Fabien Petitjean, Judit Planas, Pranav Rai, Srikanth Ramaswamy, Michael W. Reimann, Juan Luis Riquelme, Nadir Román Guerrero, Ying Shi, Vishal Sood, Mohameth François Sy, Werner Van Geit, Liesbeth Vanherpe, Tamás F. Freund, Audrey Mercer, Eilif Muller, Felix Schürmann, Alex M. Thomson, Michele Migliore, Szabolcs Káli, Henry Markram

## Abstract

The CA1 region of the hippocampus is one of the most studied regions of the rodent brain, thought to play an important role in cognitive functions such as memory and spatial navigation. Despite a wealth of experimental data on its structure and function, it has been challenging to reconcile information obtained from diverse experimental approaches. To address this challenge, we present a community-driven, full-scale *in silico* model of the rat CA1 that integrates a broad range of experimental data, from synapse to network, including the reconstruction of its principal afferents, the Schaffer collaterals, and a model of the effects that acetylcholine has on the system. We tested and validated each model component and the final network model, and made input data, assumptions, and strategies explicit and transparent. The unique flexibility of the model allows scientists to address a range of scientific questions. In this article, we describe the methods used to set up simulations that reproduce and extend *in vitro* and *in vivo* experiments. Among several applications in the article, we focus on theta rhythm, a prominent hippocampal oscillation associated with various behavioral correlates and use our computer model to reproduce and reconcile experimental findings. Finally, we make data, code and model available through the hippocampushub.eu portal, which also provides an extensive set of analyses of the model and a user-friendly interface to facilitate adoption and usage. This neuroscience community-driven model represents a valuable tool for integrating diverse experimental data and provides a foundation for further research into the complex workings of the hippocampal CA1 region.

## 1 Introduction

The hippocampus is thought to play a fundamental role in cognitive functions such as learning, memory, and spatial navigation (O’Keefe & Dostrovsky, 1971; Morris et al., 1982). It consists of three subfields of *cornu ammonis* (CA), CA1, CA2, and CA3 (see Amaral and Witter, 1989). CA1, for instance, one of the most studied, provides the major hippocampal output to the neocortex and many other brain regions (e.g. Soltesz and Losonczy, 2018). Therefore, understanding the function of CA1 represents a significant step towards explaining the role of hippocampus in cognition.

Each year the large neuroscientific community studying hippocampus contributes thousands of papers to an existing mass of empirical data collected over many decades of research (see Figure S1). Recent reviews have, however, highlighted gaps and inconsistencies in the existing literature (Bezaire & Soltesz, 2013; Wheeler et al., 2015; Pelkey et al., 2017; Sanchez-Aguilera et al., 2021). Currently, the community lacks a unifying, multiscale model of hippocampal structure and function with which to integrate new and existing data.

Computational models and simulations have emerged as crucial tools in neuroscience for consolidating diverse multiscale data into unified, consistent, and quantitative frameworks that can be used to validate and predict dynamic behavior (Fan & Markram, 2019). However, constructing such models requires assigning values to model parameters, which often involves resolving conflicts in the data, filling gaps in knowledge, and making explicit assumptions to compensate for any incomplete data. In order to validate the model, it must be tested under specific experimental conditions using independent sources of empirical evidence before the model can be used to generate experimentally testable predictions. Therefore, the curation of a vast range of experimental data is a fundamental step in constructing and parametrizing any data-driven model of hippocampus.

The challenge of incorporating these data into a comprehensive reference model of hippocampus, however, is considerable and calls for a community effort. While community-wide projects are common in other disciplines (e.g. Human Genome Project in bioinformatics, CERN in particle physics, NASA’s Great Observatories program in astronomy - Rockström et al., 2009; Hood and Rowen, 2013; Aad and Abbott, 2015), they are a relatively recent development in neuroscience. OpenWorm, for example, is a successful, decade-long community project to create and simulate a realistic, data-driven reference model of the roundworm *Caenorhabditis elegans* (*C. elegans*) including its neural circuitry of *∼*302 neurons to study the behavior of this relatively simple organism *in silico* (Szigeti et al., 2014; Gerkin et al., 2018). By contrast, for the hippocampus, with a circuit many orders of magnitude larger than *C. elegans*, models have typically been constructed with a minimal circuit structure on a relatively small scale and often their model parameters have been tuned with the goal of reproducing a single empirical phenomenon (see Sutton and Ascoli, 2021). Comparing the results from a variety of circuit models is problematic because they vary in their degree of realism and frequently rely on one or a few single neuron models making generalization of their findings difficult (see Sutton and Ascoli, 2021). While these focused models have led to valuable insights (see M. E. Hasselmo et al., 2020), this piecemeal approach fails to demonstrate whether these separate phenomena can be reproduced in a full circuit model without the need to adjust parameters.

Large-scale circuit models of hippocampus using realistic multi-compartment spiking neuronal models pioneered by Traub and colleagues (Traub et al., 1988; Traub & Miles, 1991; Traub et al., 1992, 2000) have been used to explain key characteristics of oscillatory activity observed in hippocampal slices and to examine the origins of epilepsy in region CA3. More recently, with significant increases in high-performance computing resources, Cutsuridis et al. (2010) in a microcircuit model of CA1 and notably Bezaire et al. (2016) in a full-scale CA1 model, have examined the contribution of diverse types of interneurons to the generation of prominent theta (4-12 Hz) oscillations. While these large-scale circuit models provide a more holistic approach, they still need to incorporate other features to improve their realism. For instance, to better reflect the highly curved shape of the hippocampus, an atlas-based structure that more closely mimics anatomy is required. Additionally, models need to employ pathway-specific short-term synaptic plasticity known to regulate circuit dynamics and neural coding (Tsodyks & Markram, 1997). While Yu et al. (2020) have constructed a down-scaled, atlas-based model of the rat dentate gyrus (DG) to CA3 pathway, there has to date been no atlas-based, full-scale model of rat CA1 (For a more detailed comparison of these models, see Table S2).

To initiate a community effort of this magnitude requires an approach that standardizes data curation and integration of diverse datasets from different labs and uses these curated data to construct and simulate a scalable and reproducible circuit automatically. A reconstruction and simulation methodology was introduced and applied at the microcircuit scale, for the neocortex (Markram et al., 2015) and the thalamus (Iavarone et al., 2023) and at full-scale for a whole neocortical area (Reimann et al., 2022; Isbister et al., 2023). However, these models relied primarily on datasets collected specifically for the purpose rather than data sought from and curated with the help of the scientific community.

In this paper, we describe a community-driven reconstruction and simulation of a full-scale, atlas-based multiscale structural and functional model of the area CA1 of the hippocampus that extends and improves upon the approach described in Markram et al. (2015). To model stimuli originating from beyond the intrinsic circuitry, we included the synaptic input from the Schaffer collaterals (SC) from CA3, which is the largest afferent pathway to CA1 and the most commonly stimulated in experiments. Furthermore, we also added the neuromodulatory influence of cholinergic inputs, perhaps the most studied neuromodulator in the hippocampus (Teles-Grilo Ruivo & Mellor, 2013). We constrained all model parameters and data using available experimental data from different labs or explicit assumptions made when data were lacking. We extensively tested and validated each model component and the final network to assess its quality. To maximize realism of the simulations, we set up simulation experiments to represent as closely as possible the experimental conditions of each empirical validation. We demonstrated the broad applicability of the model by studying the generation of neuronal oscillations, with a specific focus on theta rhythm, in response to a variety of different stimulus conditions. Over time and with the help of the community, limitations of the model revealed by these processes can be addressed to improve upon it. To facilitate a widespread adoption by the community, we have developed a web-based resource to share the model and its components, open sourcing extensive analyses, validations, and predictions that can be accessed as a complement to direct interaction with the model (hippocampushub.eu). Finally, we have developed a massive online open course (MOOC) to introduce users to the building, analysis, and simulation of a rat CA1 microcircuit (https://www.edx.org/course/simulating-a-hippocampus-microcircuit) providing a smaller version of the full-scale model for education purposes.

## 2 Results

We divide the results section into model reconstruction and simulation of the oscillatory activity in CA1. For a list of abbreviations and acronyms used in the paper, see Table S1.

### 2.1 Model reconstruction

In this section, we describe how we reconstructed the main components of the model: the cornu Ammonis 1 (CA1), the Schaffer collaterals (SC), and the effect of acetylcholine (ACh) on CA1. Each main component is a compound model of several “building blocks” (Figures 1 and S2) for which we show how, from the sparse data available in the literature (see Tables S3-S24) and a list of assumptions (Section 4.26), we arrived at the dense data necessary to ascribe a value for each model parameter. Finally, we show validations of the building blocks to assess their validity and robustness.

**Figure 1:**
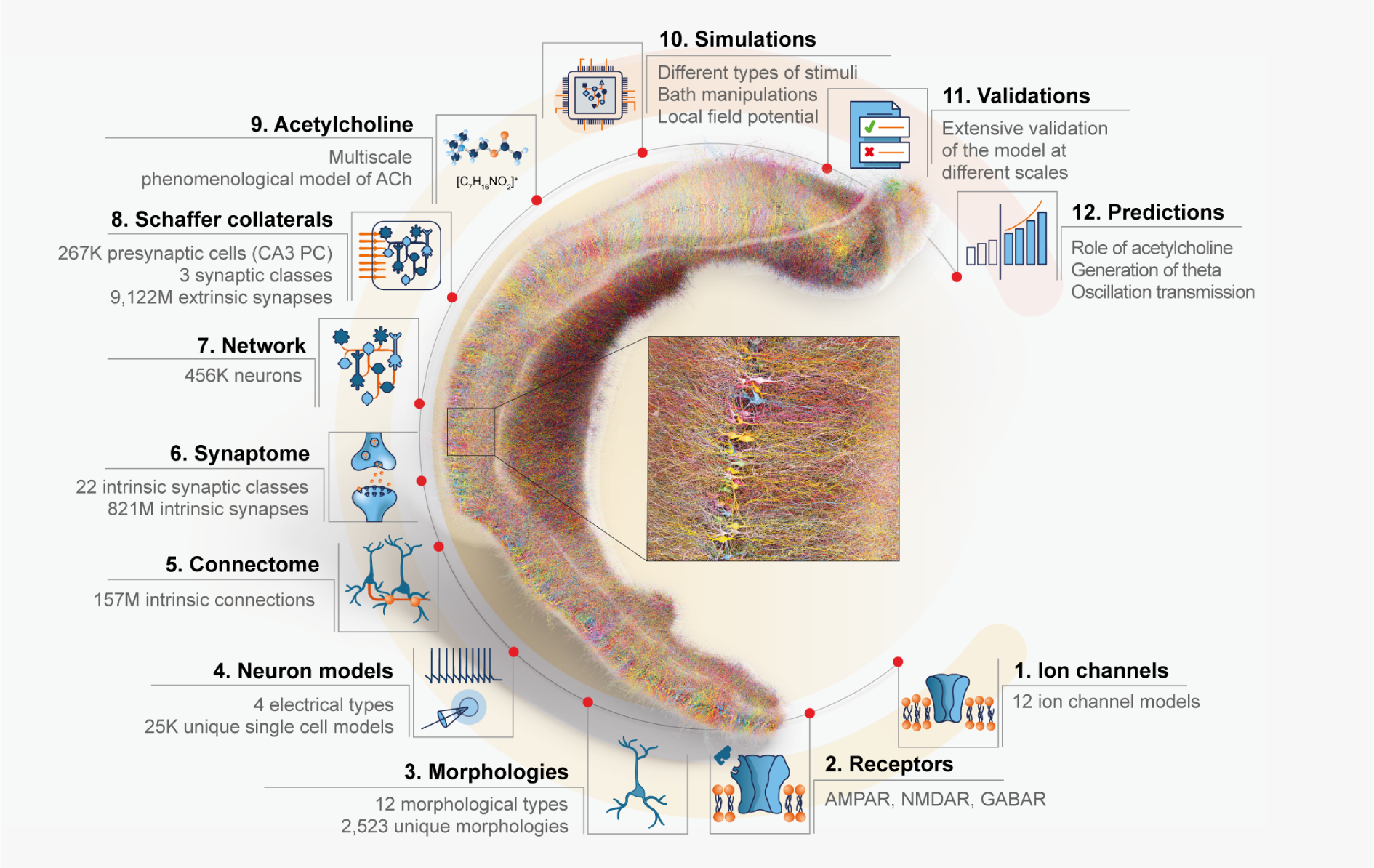
Overview of the model. A visualization of a full-scale, right-hemisphere reconstruction of rat CA1 region and its components. The number of cells is reduced to 1% for clarity, and neurons are randomly colored. The CA1 network model integrates entities of different spatial and temporal scales. The different scales also reflect our bottom-up approach to reconstruct the model. Ion channels (1) were inserted into the different morphological types (3) to reproduce electrophysiological characteristics and obtain neuron models (4). Neurons were then connected by synapses to generate an intrinsic CA1 connectome (5). For each intrinsic pathway, synaptic receptors (2) and transmission dynamics were assigned based on single neuron paired recording data (6) to create a functional intrinsic CA1 network model (7). The intrinsic CA1 circuit received synaptic input from CA3 via Schaffer collateral (SC) axons (8). The neuromodulatory influence of cholinergic release on the response of CA1 neurons and synapses was modeled phenomenologically (9). The dynamic response of the CA1 network was simulated with a variety of manipulations to model *in vitro* and *in vivo*, intrinsic and extrinsic stimulus protocols while recording intracellularly and extracellularly (10) to validate the circuit at different spatial scales against specific experimental studies (11) and to make experimentally testable predictions (12).

#### 2.1.1 Building CA1

To reconstruct a full-scale model of rat CA1, we created biophysical models of its neurons, defined an atlas volume of the region for one hemisphere, placed these neurons in the volume, and connected them together by following and adapting the method described in Markram et al. (2015) (Figure S2).

##### CA1 neurons

To achieve a full-scale version of CA1, we needed to populate the model with *∼*456,000 cells. We started by curating 43 morphological reconstructions of neurons belonging to 12 morphological types: pyramidal cell (PC), axo-axonic cell (AA), two subtypes of bistratified cell (BS), back-projecting cell (BP), cholecystokinin (CCK) positive basket cell (CCKBC), ivy cell (Ivy), oriens lacunosum-moleculare cell (OLM), perforant pathway associated cell (PPA), parvalbumin positive basket cell (PVBC), Schaffer collateral associated cell (SCA), and trilaminar cell (Tri). To increase the morphological variability, we scaled and cloned them producing an initial morphology library of 2,592 reconstructions.

To validate the resulting morphology library, we compared them morphometrically and topologically to the original morphologies. The similarity scores for the distribution of morphological features were statistically similar (Figure S3, all values *R >* 0.98, *p <* 10*^−^*^25^). Using the Topological Morphology Descriptor (TMD) (Kanari et al., 2018), the persistence diagrams (Figure S4) show an increase in morphological variability introduced by the cloning process (details per m-type in Figures S5 and S6).

To produce electrical models (e-models), we began by taking 154 single-cell recordings and classifying traces into four electrical types (e-types) using Petilla nomenclature (Petilla Interneuron Nomenclature Group et al., 2008): classical accommodating for pyramidal cells and interneurons (cACpyr, cAC), bursting accommodating (bAC), and classical non-accommodating (cNAC). From each trace, we extracted electrical features (e-features) which we used in combination with the curated morphologies to produce and validate 36 single-cell e-models (R. Migliore et al., 2018; Ecker et al., 2020). In the case of pyramidal cells e-models, they showed a close correlation to experimental findings in terms of back-propagating action potential (BPAP) (Golding et al., 2001, *R* = 0.878, *p* = 0.121) and post-synaptic potential (PSP) attenuation (Magee and Cook, 2000, *R* = 0.846, *p* = 0.001, Figure S7).

To match the proportions of the morpho-electrical type (me-type) composition of the CA1 (Table S4), we combined the 36 e-models with 2,592 curated morphologies to obtain an initial library of 26,112 unique me-type models. This is the pool of biophysical cell models available to populate the full-scale version of CA1.

##### Defining the spatial framework

To represent the CA1 spatial volume, we started with a publicly available atlas reconstruction of the hippocampus (Ropireddy et al., 2012). Our aim was to create a continuous coordinate system to represent the three axes of the hippocampus (longitudinal, transverse, and radial) and a precise vector field for cell placement and orientation (Figure 2). For our building and analysis algorithms to work effectively, we applied a series of post-processing steps (Figure S8, see Section 4.13). Within this process, we redefined the layers parametrically to be consistent with the layer thicknesses in our datasets (Section 4.13.3).

**Figure 2:**
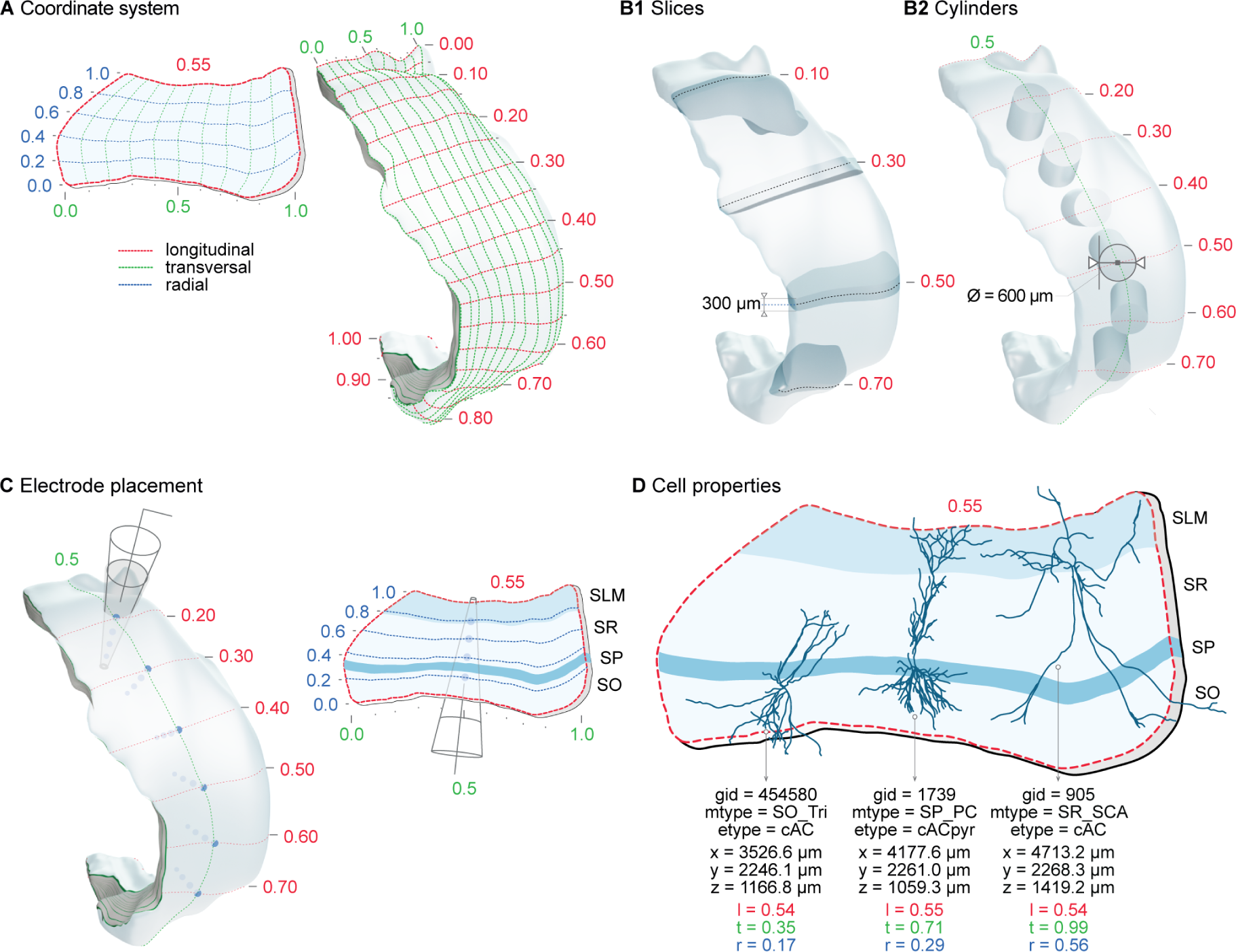
Coordinate system. Custom parametric coordinates system used as spatial reference for circuit building, circuit segmentation, and for simulation experiments. A. Longitudinal (l, red), transverse (t, green) and radial (r, blue) axes of the CA1 volume are defined parametrically in range [0,1]. Left: Slice from volume shows radial depth from SO/alveus (r=0) to SLM/pial (r=1) and transverse extent from CA3/proximal CA1 (t=1) to distal CA1/subiculum (t=0) boundaries. Right: Full volume shows surface grid of transverse vs longitudinal axes. Longitudinal axis extends from dorsal (l=0) to ventral (l=1) CA1. B. Circuit segmentation for analysis and simulation. Coordinates system used to select CA1 slices of a given thickness (B1) or a cylinders of a given diameter (B2) at specific locations along longitudinal axis. C. Extracellular electrode placed at a given surface position (left) and channels at selected laminar depth (right) in CA1 volume. D. Each neuron in the circuit is defined by a unique general identifier (gid), its morphological type (m-type), electrical type (e-type), spatial xyz-coordinates and parameterized ltr-coordinates.

##### Placing neurons in the volume

After defining the volume, we wanted the model to match the neuronal density and proportion of cell types in rat CA1. We compiled available data to derive the cellular composition (Section 4.14, see Table S3) and used it to populate the atlas with soma locations (Figure S9A,B). To check the consistency, we validated the resulting cell composition (*R* = 0.999, *p* = 1.31 *×* 10*^−^*^28^) and cell density (*R* = 0.999, *p <* 0.0001) (Figure S10B,C, Table S5). At each soma location, we needed to select from the e-model library the neuron that best fits in the space of the layers. To this purpose, we oriented its morphology according to the vector fields (Figure S9C,D) and evaluated it against a set of rules that describe the target distribution of neurites per layer (Table S6). Visually, cells in our model follow the curvature of the hippocampus and the different parts of the cells target the expected layers (Figure S10A). Subject to the multiple constraints of the cell placement algorithm, we placed 456,380 neurons in the volume, utilizing 2,523 unique morphologies and 25,355 unique neuron models.

##### Connecting CA1 neurons

To connect the placed neurons, we used the connectome algorithm previously described in Reimann et al. (2015). In brief, the algorithm searches for co-localization of axon and postsynaptic neurons within a certain distance to identify a potential synapse (or apposition). After identifying all potential synapses, a subsequent pathway-specific pruning step discards some to match the known bouton densities (Table S7) and number of synapses per single axon connection (Table S9). This algorithm has been demonstrated to accurately recreate local connectivity (Markram et al., 2015; Reimann et al., 2022; Iavarone et al., 2023) as well as higher-order topological features (Gal et al., 2017). The resulting intrinsic connectome consisted of about 821 million synapses.

Given the importance of the connectome, we wanted to validate it as widely as possible to mitigate the uncertainty in our assumptions and literature data. In the case of pathways for which we had no specific data, we were instead able to offer testable, model-based predictions (Figures S12, S13, S14, and S15). First, we verified the bouton density and number of synapses per connection used in the pruning step was preserved in the generated connectome (bouton density: *R* = 0.909, *p* = 0.0120; number of synapses per connection: *R* = 0.992, *p* = 2.41 *×* 10*^−^*^9^, Figure S12 and Tables S7 and S9). Next, we observed that the shape of the distributions for connection probability (Figure S13A), convergence (Figure S14A) and divergence (Figure S15A) were positively skewed as reported experimentally (Giacopelli et al., 2021). In the case of mean connection probability, experimental data did not allow a direct comparison because the distance between the neuron pairs tested was typically missing (Figure S13C, Table S8). For convergence, we found that the sub-cellular distribution of synapses on different compartments of pyramidal cells in our model was consistent with Megias et al. (2001) (*R* = 0.988, *p* = 0.012, Figure S14D). For divergence, the model did not always closely match the experimental data for the total number of synapses per axon formed by certain m-types (*R* = 0.524, *p* = 0.286, Figure S15C, Table S10). We compared divergence also in terms of the percentage of synapses formed with PCs or INTs (Figure S15D, Table S11) and validated the distribution of efferent synapses in the different layers (SO: *R* = 0.798, *p* = 0.057; SP: *R* = 0.905, *p* = 0.013; SR: *R* = 0.813, *p* = 0.049; SLM: *R* = 0.999, *p* = 4.11 *×* 10*^−^*^8^, Figure S15E, Table S12). Overall, this suggests the model connectome provides a reasonable approximation based on available data, while the discrepancies can be due, for example, to the small sample size and high variability in axon length recovered from *in vitro* slices.

To provide functional dynamics for synaptic connections with stochastic neurotransmitter release and short-term plasticity (see Figure S16), we used the optimized parameters we previously derived in Ecker et al. (2020) for the 22 intrinsic synaptic classes of pathways we have identified (Tables S13 and S14). The model was able to reproduce the post-synaptic potential (PSP) amplitudes (Figure S16D, R = 0.999, *p* = 1.65 *×* 10*^−^*^19^) and post-synaptic current (PSC) coefficient of variation (CV) of the first peak (Figure S16E, *R* = 0.840, *p* = 0.018) for the pathways with available electrophysiological recordings. After having constrained and validated the synapses, the reconstruction process of the intrinsic CA1 circuit is complete.

#### 2.1.1 Reconstruction of Schaffer collaterals (SC)

An isolated CA1 does not have substantial background activity (Harris & Stewart, 2001), while normally the network is driven by external inputs. The Schaffer collaterals from CA3 pyramidal cells are the most prominent afferent input to the CA1 and the most studied pathway in the hippocampus (Szirmai et al., 2012; Dumas et al., 2018). Their inclusion allows us to deliver synaptic activity input patterns to the CA1.

To reconstruct the anatomy of Schaffer collaterals, we constrained the number of CA3 fibers and their average convergence on CA1 neurons with literature data (Tables S15 and S16). Due to scarce topographical information, we distributed synapses uniformly along the transverse and longitudinal axes, while along the radial axis we followed a layer-wise distribution as reported by Bezaire and Soltesz (2013). The resulting Schaffer collaterals added more than 9 billion synapses to CA1, representing 92% of total modelled synapses. As expected, the synapse laminar distribution (Figure 3A-C) and mean convergence on PCs and interneurons (Figure 3D,E) match experimental values (one-sample t-test, *p* = 0.957 for PCs and *p* = 0.990 for INTs). Interestingly, other unconstrained properties also match experimental data. The convergence variability is comparable with the upper and lower limits identified by Bezaire and Soltesz (2013) (model PC: 20,878 *±* 5,867 synapses and experimental PC: 13,059 - 28,697, model INT: 12,714 *±* 5,541 and experimental INT 7,952 - 17,476, Figure 3D,E). In addition, the axonal divergence from a single CA3 PC is 34,135 *±* 185 synapses (Figure S17A), close to the higher end of the ranges measured by Sik et al. (1993), Li et al. (1994), and Wittner et al. (2007) (15,295 - 27,440, Figure S17B). Finally, most of the connections formed a single synapse per neuron (1.0 *±* 0.2 synapses/connection, Figure S17A), consistent with what has been previously reported (Bezaire & Soltesz, 2013).

**Figure 3:**
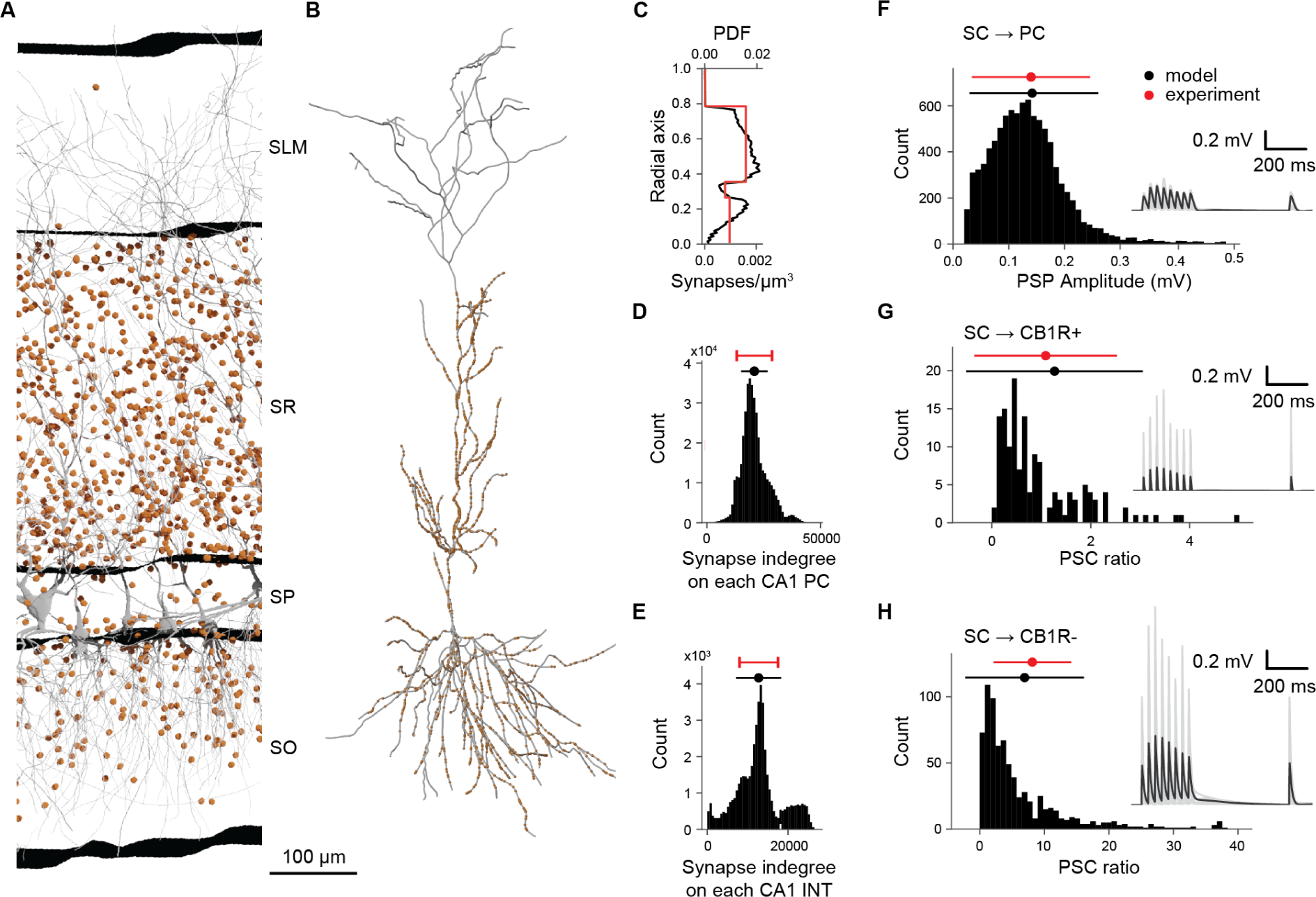
Schaffer collaterals anatomy and physiology. A. Section of a slice of the dorsal CA1 showing neurons in gray and SC synapses in orange (10% of the existing ones). B. Example of SC synapse placement (orange dots) on one reconstructed PC (in grey). C-H. Validation of the anatomy (C-E) and physiology (F-H) of the SC. Density of SC synapses (lower x axis) and probability density function (PDF) (upper x axis) at different depths (radial axis percentage) (C). Distributions of afferent synapses from SC to PC (D) and INT (E). Distribution of PSP amplitudes for SC *→* PC synapses (F). Distribution of PSC ratio (see text) for SC *→* CB1R+ (G) and SC *→* CB1R-(H). Insets in panels F-H report voltage membrane traces of 10 randomly selected pairs of SC*→* PC, SC *→* CB1R+, and SC *→* CB1R-interneurons, respectively. The presynaptic SC is stimulated to fire 8 times at 30 Hz, plus a recovery pulse after 500 ms from the last spike of the train. Solid black lines represent mean values, and shaded gray areas the standard deviation.

The CA3 afferent pathway is sparsely connected to the CA1, so the chance of obtaining paired CA3-CA1 neuronal recordings is small between PCs and much smaller from PC to interneurons (Sayer et al., 1990; Debanne et al., 1995; Wierenga & Wadman, 2003; Milstein et al., 2015). So, to constrain SC physiology we did not have enough data to follow the parametrization used for intrinsic synapses (Ecker et al., 2020). Instead, we used the available data (Tables S17 and S18) and optimized the missing parameters as shown in Figure S17C. The resulting SC*→*PC synapses match the distribution of EPSP amplitudes as measured by Sayer et al. (1990) (Figure 3F, experiment: 0.14*±*0.11 mV, CV = 0.76, model: 0.15*±*0.12 mV, CV = 0.80, z-test *p* = 0.709), giving a peak synaptic conductance of 0.85 *±* 0.05 nS and N*_RRP_* of 12. The rise and decay time constants of AMPA receptors (respectively 0.4 ms and 12.0 *±* 0.5 ms) were obtained by matching Sayer et al. (1990) EPSP dynamics (Figure S17C, 10-90% rise time model: 5.4 *±* 0.9 ms and experiment: 3.9 *±* 1.8 ms, half-width model: 20.3 *±* 2.9 ms and experiment: 19.5 *±* 8.0 ms, decay time constant model: 19.5 *±* 2.5 ms and experiment: 22.6 *±* 11.0 ms). In the case of SC*→*INT synapses, we distinguished between cannabinoid receptor type 1 negative (CB1R-) and positive (CB1R+) interneurons (Glickfeld & Scanziani, 2006). SC*→*CB1R-synapses match *EPSC_CBR_*_1_*_−_/EPSC_P C_* experimental ratio (Glickfeld & Scanziani, 2006) (model 6.950 *±* 9.200 and experiment 8.15 *±* 6.00, z-test *p* = 0.18, Figure 3G), resulting in a peak conductance of 15.0 *±* 1.0 nS and N*_RRP_* of 2. SC*→*CB1R+ synapses match the *EPSC_CBR_*_+_*/EPSC_P C_* experimental ratio (model 1.27 *±* 1.78 and experiment 1.09 *±* 1.44, z-test *p* = 0.06, Figure 3H), giving peak conductance of 1.5 *±* 0.1 nS and N*_RRP_* of 8. SC*→*INT synapses (all) match the timing in the EPSP-IPSP sequence of Pouille and Scanziani, 2001 (model: 2.69 *±* 1.18 ms, experiment: 1.9 *±* 0.6, Figure S17E), yielding a rise and decay constants for AMPA receptors of 0.1 ms and 1.0 *±* 0.1 ms, respectively. This short latency gives effective feedforward inhibition, which is a key aspect for the transmission of oscillations from CA3 to CA1 (see below).

We functionally validated SC projections reproducing Sasaki et al. (2006), where the authors examined the basic input-output (I-O) characteristics of SC projections *in vitro*. The SC pathway is thought to be dominated by feedforward inhibition, which increases the dynamic range of the CA1 network and linearizes the I-O curve (Sasaki et al., 2006; Pouille et al., 2009). Blocking gamma-aminobutyric acid receptor (GABA_A_R) drastically reduces the dynamic range of the network resulting in an I-O curve that saturates very quickly. To match the methodology of Sasaki et al. (2006), we set up the simulations to be as close as possible to the experimental conditions (slice of 300 µm, *Ca*^2+^ 2.4 m M, *Mg*^2+^ 1.4 m M, 32 °C) (Figure 4A) and we used the same sampling strategy: randomly sampling 101 neurons in the slice to find how many SC axons were required to make all of them fire (representing respectively 100% of the input and 100% of output, Figure 4B). To assess the role of feedforward inhibition, we mimic the effect of gabazine by disabling the connections from interneurons. The model captured the quasi-linearization of the I-O response in control conditions (Pearson test on linearity R = 0.992, p = 2.56 *×* 10*^−^*^9^) and the rapid saturation of the CA1 network with the simulated “no GABA” condition (Figure 4B). In control conditions, at 50% of input intensity (Figure 4C) the spiking activity of CA1 SP neurons is rather weak and rapidly suppressed by the feedforward inhibition, while without inhibition CA1 neurons fire for more than 50 ms at high frequency (up to 200 Hz). Taken together, these results suggest the SC projection represents a valid model given the available empirical data.

**Figure 4:**
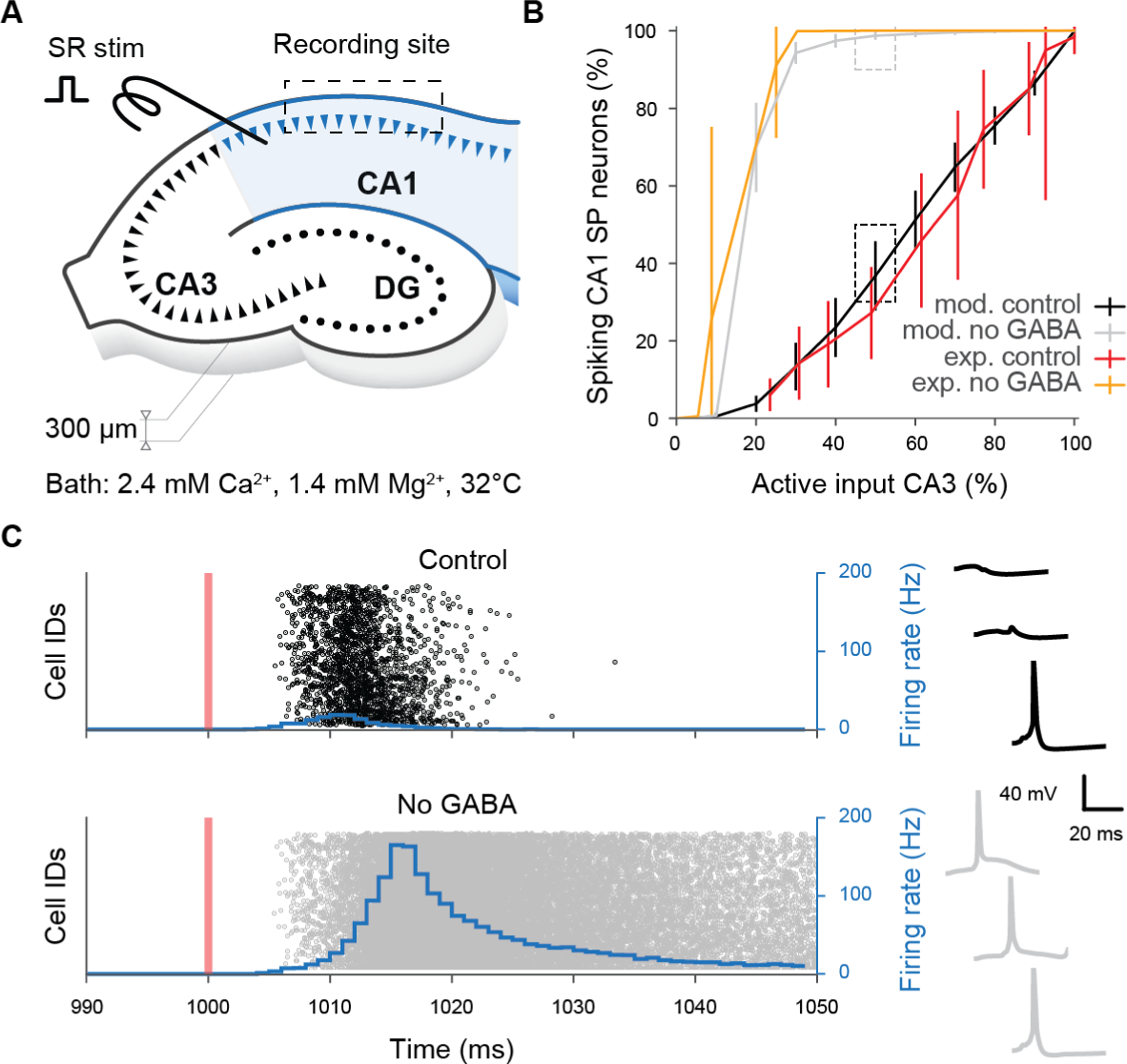
Schaffer collaterals validation. Effect of the feedforward inhibition on the input-output relationship of the network illustrated in a slice experiment. A. The illustration (redrawn from Figure 1A of Sasaki et al. (2006)) shows the *in silico* experimental setup. B. Response of 101 selected neurons to an increasing number of stimulated SC fibers, with and without GABA. The dashed boxes identify the condition (50% of active SC) that is used to show the model’s results in panel C. C. Raster plots of SP neurons in response to the SC stimulation (orange vertical line) with the overlaying firing rate (blue). On the right, membrane voltage traces of three randomly selected SP neurons in control (black) and no GABA (gray) conditions.

#### 2.1.1 Cholinergic modulation

The behavior of the hippocampus is shaped by several neuromodulators, with acetylcholine (ACh) among the most studied. Cholinergic fibers originate mainly from the medial septum (MS) and have been correlated with phenomena such as theta rhythm, plasticity, memory retrieval and encoding, as well as pathological conditions such as Alzheimer’s disease (Dannenberg et al., 2017). This section describes the reconstruction of a phenomenological model of ACh, quantifying the effects of ACh on neurons and synapses, and developing a novel method to integrate available experimental data (Tables S19 and S20, Figure 5A,B). The data used to build the model was obtained from *in vitro* application of various cholinergic agonists such as ACh and carbachol (CCh); here we assume that their effects are comparable (Colangelo et al., 2019). We modelled the effect of ACh on neurons and synapses. The effect on neurons results in an increase in resting membrane potential or firing frequency. We were able to integrate both types of experimental data by estimating the net current that is required to evoke the corresponding changes for a given concentration of ACh (see equation 9 in Methods). ACh affects synaptic transmission acting principally at the level of release probability (Tsodyks & Markram, 1997; M. E. Hasselmo, 2006; D. Yang et al., 2021) (see equation 10 in Methods).

**Figure 5:**
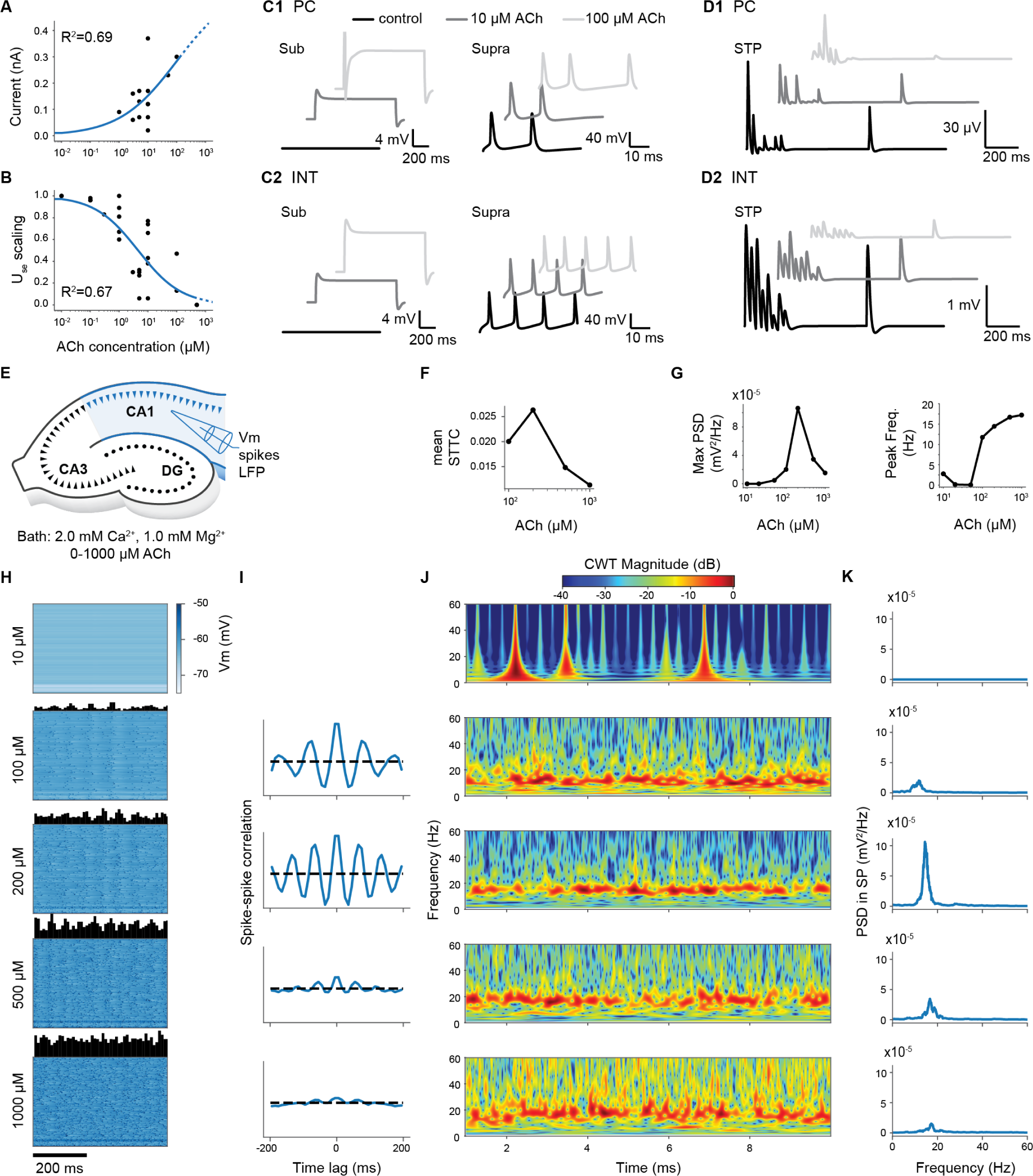
Effects of Acetylcholine on neurons, synapses, and network. A. Dose-response modulation of neuronal excitability caused by ACh. Back dots are experimental data points; blue curve represents the fitted equation. The dashed part of the curve indicates regions outside available experimental data. B. Dose-response modulation of synaptic release. Same color code as in A. C. Example traces for PC (C1) and interneurons (C2) in sub-threshold and supra-threshold conditions, with different concentrations of ACh. D. Example traces showing the STP dynamics for PC (D1) and interneurons (D2) at different concentrations of ACh. E. The illustration shows the *in silico* experimental setup to analyze network effects of ACh. Different concentrations of CCh are applied to the circuit, and multiple types of recordings made in the CA1 (membrane voltage, spike times, LFPs). F. Mean spike time tiling coefficient (STTC) as a function of ACh concentration. G. Maximum of the LFP power spectrum density (PSD) and peak frequency as a function of ACh concentration. H. The voltage of 100 randomly selected neurons at different levels of ACh. The upper histograms show the instantaneous firing rate. I. Spike-spike correlation histograms. J. Spectrogram of the LFP measured in SP at different ACh levels. K. PSD of Once we had modelled the effect of ACh on neuron excitability and synaptic transmission, we validated the effect of ACh at the network level against available data (see Table S21). To accomplish this, we simulated bath application of ACh for a wide range of concentrations (from 0 µM (i.e., control condition) to 1000 µM) (Figure 5E,F). We observed a subthreshold increase in the membrane potential of all neurons for values of ACh lower than 50 µM, without any significant change in spiking activity. At intermediate doses (i.e., 100 µM and 200 µM), the network shifted to a more sustained activity regime. Here, we observed a generalized increase in firing rate as ACh concentration increased and a progressive build-up of coherent oscillations whose frequencies ranged from 8 to 16 Hz (from high theta to low beta frequency bands). The correlation peak between CA1 neurons occurred at 200 µM ACh (Figure 5G,H). At very high concentrations (i.e., 500 µM and 1000 µM) we observed a decrease in the power of network oscillations, which was further confirmed by analysis of local field potential (LFP) (Figure 5I). Power spectral density (PSD) analysis showed a maximum absolute amplitude for 200 µM ACh with a peak frequency of *∼*15 Hz (Figure 5J,K). Higher concentrations decreased the maximum amplitude of the PSD while the peak frequency converged toward 17 Hz (Figure 5K). Thus, we observe the emergence of three different activity regimes at low, intermediate, and high levels of cholinergic stimulation. It is unclear whether the network behavior we observe is consistent with all experimental findings because of their different methodologies. Nevertheless, this phenomenological model of ACh allows the reproduction of other experiments in which ACh or its receptor agonists are necessary (see, for example, Sections 2.2.1 and 2.2.2).

### 2.2 Model simulation and investigations

By following a data-driven approach and independently validating each model component, we have arrived at a candidate reference model. It can serve as a basis for investigating several scientific questions where parameters are only adjusted to reflect the different experimental setups, rather than tuned to achieve a specific network behavior. A simulation experiment is essentially a model of the experimental setup that is reproduced with as much accuracy as possible within the limits of the model. As presented in the Model reconstruction section we can, for example, simulate slices of a certain thickness, change the extracellular concentration of ions, change temperature, and enable spontaneous synaptic events. Here, we show several simulations with particular emphasis on theta oscillations, a prominent network phenomenon observed in the hippocampus *in vivo* and related to many behavioral correlates (Buzsáki, 2005). Then, we go beyond the theta oscillations and test whether a wider range of frequencies can pass through SC more reliably than others.

#### 2.2.1 Theta oscillations

During locomotion and REM sleep, CA1 generates a characteristic rhythmic theta-band (4-12 Hz) extracellular field potential (Jung & Kornmüller, 1938; Green & Arduini, 1954; Grastyan et al., 1959; Vanderwolf, 1969; Goyal et al., 2020). Neurons in many other brain regions such as neocortex are phase-locked to these theta oscillations (Siapas et al., 2005; Sirota et al., 2008) suggesting hippocampal theta plays a crucial coordinating role in the encoding and retrieval of episodic memory during spatial navigation (Buzsáki, 2002, 2005). Yet despite more than eighty years of research, the trigger that generates theta oscillations in CA1 remains unclear because of conflicting evidence. This represents an opportunity to use our reference model to investigate these inconsistencies and gain an improved understanding of theta generation. Specifically, we investigate three main candidate mechanisms proposed in literature (Colgin, 2013): intrinsic CA1 generation and extrinsic pacemaker oscillations from CA3 or from medial septum (MS).

##### CA1 generation

To investigate possible intrinsic mechanisms of theta rhythm generation in CA1 (Goutagny et al., 2009; Ferguson et al., 2017), we examined three candidate sources of excitation that might induce oscillations: (i) spontaneous synaptic release or miniature postsynaptic potentials (minis or mPSPs), (ii) homogeneous random spiking of SC afferent inputs, and (iii) varying bath concentrations of extracellular calcium and potassium to induce tonic circuit depolarisation. While in their CA1 circuit model Bezaire et al. (2016) reported random synaptic activity was sufficient to induce robust theta rhythms, in our model we found none of these candidates generated robust theta rhythms. For minis, we found setting release probabilities to match empirically reported mPSP rates (Table S22) led to irregular, wide-band activity in CA1 (see Figure S18). For random synaptic barrage, varying presynaptic rate to match the mean postynaptic firing rate of pyramidal cells in Bezaire et al. (2016) resulted in irregular beta-band, not regular theta-band oscillations (see Figures S19 and S20). For tonic depolarisation, within a restricted parameter range it was possible to generate theta oscillations around 10-12 Hz, but their intensity was variable and episodic (see Figures S21 and S22). Therefore, we could not find any intrinsic mechanism capable of generating regular theta activity in our circuit.

##### CA3 input

To mimic the transmission of CA3 theta oscillations to CA1, we generated SC spike trains across a range of theta-modulated sinusoidal rate functions (signal frequency) at different mean individual rates (cell frequency) (Figure 6A). We delivered these stimuli at two different calcium levels (*in vitro*-like 2 mM and *in vivo*-like 1 mM) for three different circuit scales (whole circuit, thick slice, and cylinder circuit; see Figure 2) and measured extracellular LFP and intracellular membrane potential. For LFP, we found CA1 faithfully followed the theta-modulation input frequency at both *in vivo*-like and *in vitro*-like calcium levels and at the different circuit scales tested (e.g. 8Hz, see Figure 6C-F). However, for the same stimulus, the LFP at *in vivo*-like calcium levels was around three orders of magnitude less powerful than the one at *in vitro*-like calcium levels (Figure6B,C,F) due to CA1 spiking rates being far lower (e.g. in full circuit, pyramidal cell mean firing rate of 0.00018 *±* 0.0067 (1 mM) vs 0.25 *±* 0.50 Hz (2 mM)). Therefore, due to this very low firing rate, we decided not to analyse results from *in vivo*-like conditions further and focused on those from the *in vitro*-like condition only (Figure 6C-E).

**Figure 6:**
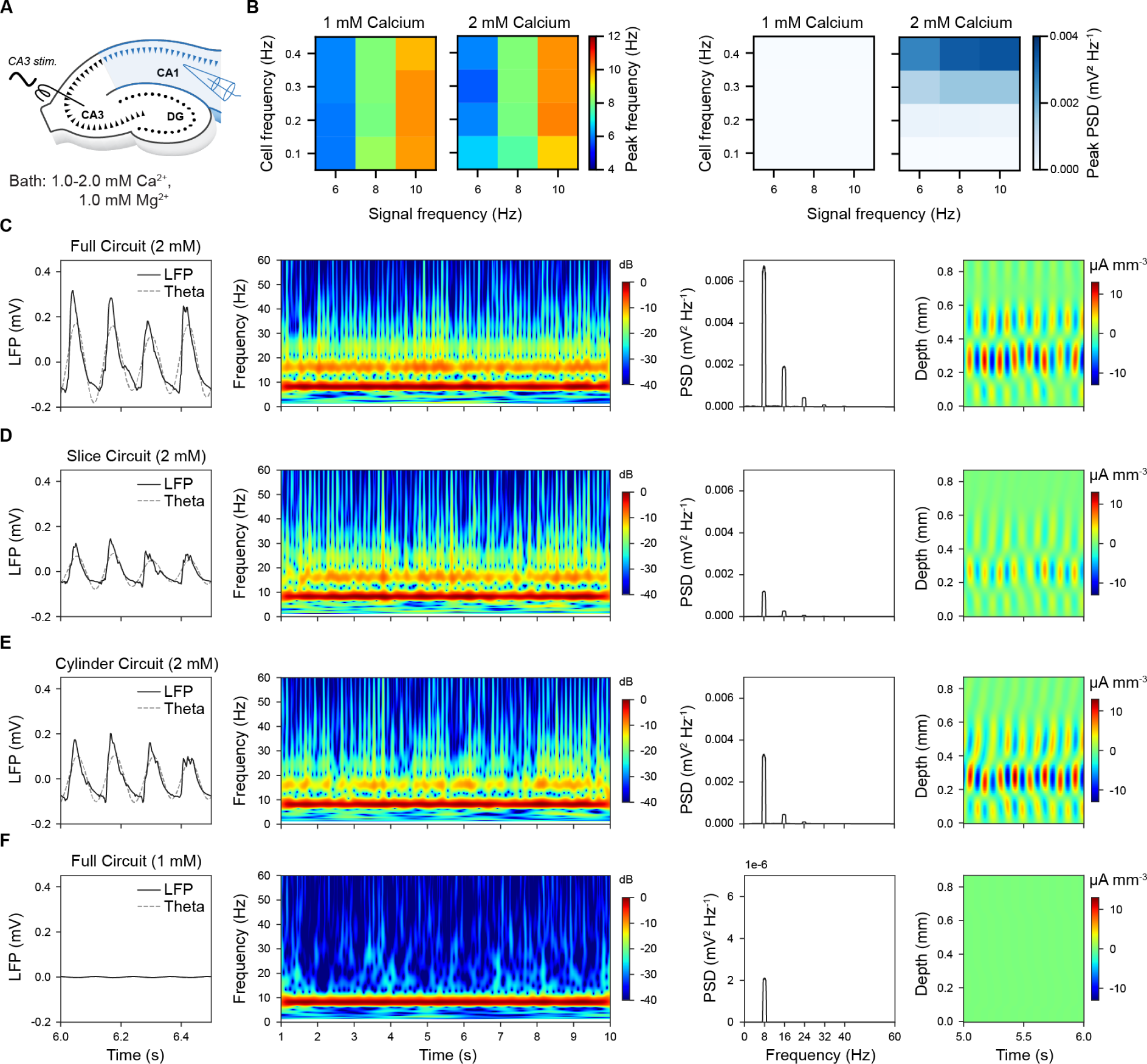
CA3 theta (8 Hz) oscillatory input entrains CA1 to matched theta oscillation across different scales of circuit. A. Schema showing the *in silico* experimental setup. B. Dependency of peak frequency (left) and PSD (right) from calcium level, cell and signal frequencies during simulations of a cylinder circuit. C. Full circuit model (2 mM calcium). LFP recordings from SP (far left), spectrogram (left middle), power spectral density (right middle), current source density (far right). D. Slice circuit model (thickness of 300 µm, 2 mM calcium). E. Cylinder circuit model (radius of 300 µm, 2 mM calcium). F. Full circuit model (1 mM calcium). Note that PSD has 1000 times smaller y-axis scaling than the ones in panels A-C).

We analyzed the properties of the model LFP and compared them to experimental data. First, we examined whether the circuit size is critical for theta generation. Contrary to Goutagny et al. (2009), we found that all the three circuit scales generated theta oscillations, but slice and cylinder circuits had reduced magnitude of the modulation and LFP power (Figure 6C-E). Second, across all the scales, the LFP waveform (Figure 6A-C, left columns) was more asymmetrical with a fast rise and slower decay (mean asymmetry index = *−*1.34 *±* 0.23; see Figure S23) with respect to what was reported during rat locomotion (asymmetry index = -0.27, Buzsáki et al. (1985)) or REM sleep periods (asymmetry index = -0.13, Belluscio et al. (2012)). Third, we observed a strong narrow-band peak of power at the same frequency as the signal that was maintained throughout the entire period of stimulation (Figure 6A-C, middle left columns). Fourth, consistent with experimental evidence (e.g. Figure 1b in Goutagny et al., 2009), first-(16 Hz) and second-order (24 Hz) harmonics of the theta modulation frequency were also present (Figure 6A-C, middle right columns). Fifth, current source density (CSD) showed a highly regular alternating current dipole between layers with a phase reversal between SP and SR (Figure 6A-C, far right column). This is similar to *in vivo* LFP recordings in the absence of perforant pathway input (Buzsáki, 2002), which we did not model. Therefore, since the results did not depend qualitatively on circuit size, for the sake of simplicity, we further analyzed only the cylinder circuit.

Next, we looked at how the spiking of different neuronal classes in the model are modulated by theta oscillations (Klausberger et al. (2003, 2004), Klausberger (2005), and Fuentealba et al. (2008); see Table S23). When the spike times of neurons close to the extracellular recording electrode in SP were compared with the phases of theta-band LFP rhythm (theta trough = 0°), all neuron types were found to respond at roughly the same phase of the theta cycle (Figure 7). As the mean rate of SC afferent spiking increased, more neurons became phase-locked yielding a denser mono-phase distribution for higher signal modulation frequencies (Figure 7A). For example, under stimuli with a 0.4 Hz SC mean spiking frequency and 8 Hz signal modulation, CA1 PCs fired first during the mid-rising phase of theta and were followed by all types of interneurons, whose spiking mostly ended before peak theta, with BS neurons emitting few or no spikes (Figure 7B left). Phase-locked neurons had a tighter tuning than *in vivo*, with pyramidal neurons typically firing before interneurons (Figure 7B middle). When compared with *in vivo* recordings of phase-locked neurons (see Table S23), the mean phase angle of model spiking was closely matched for SP_CCKBC but substantially out of phase for SP_AA (Figure 7B right). Although the angular deviation of phase-locking was generally tighter than observed *in vivo* (e.g. model vs *in vivo* for SP_AA 8.9°(n=4) vs 55.0°(n=2), SP_PVBC 12.0°(n=2) vs 68.0°(n=5), and SP_Ivy 10.9°(n=19) vs 63.1°(n=4)).

**Figure 7:**
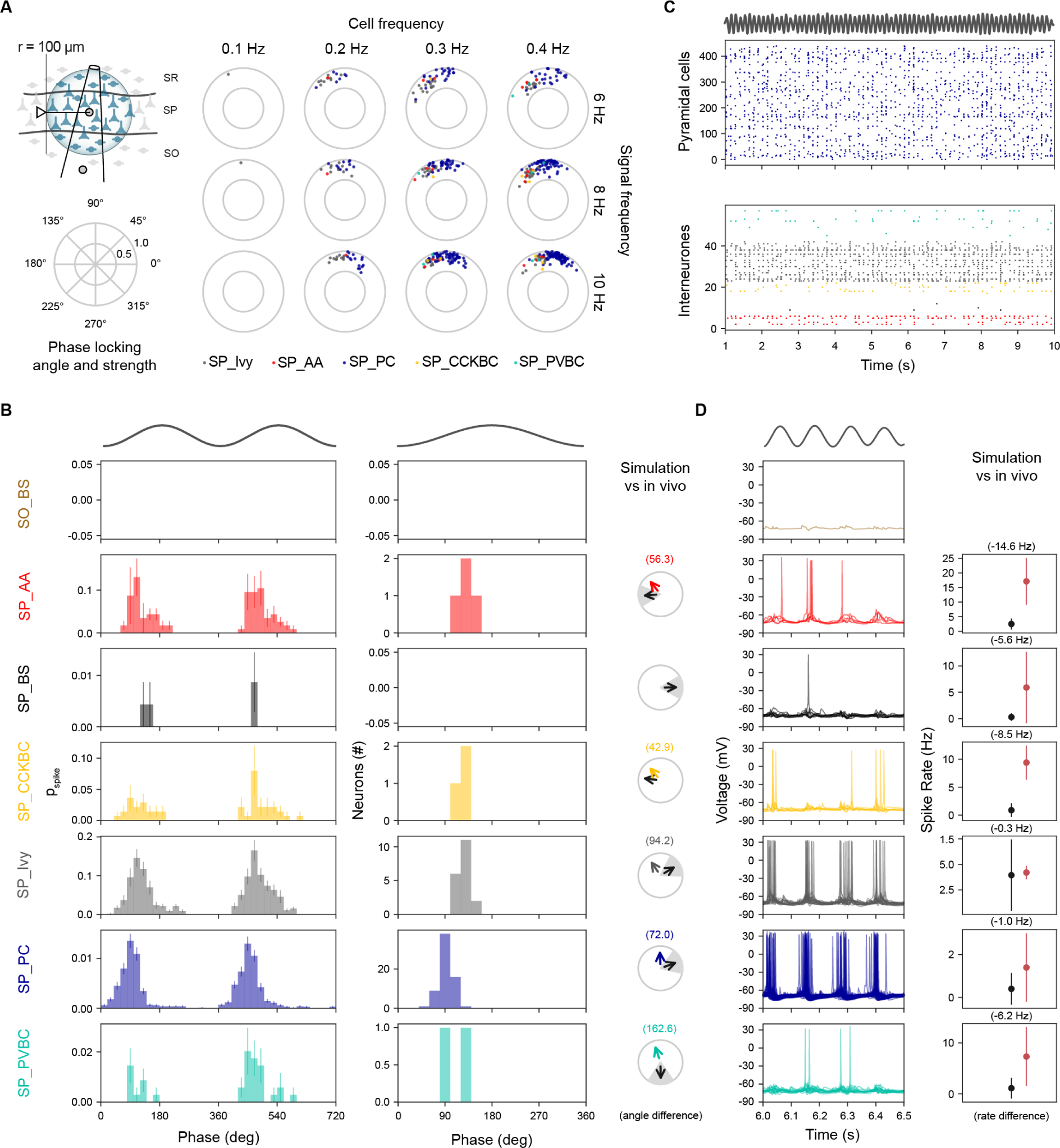
CA1 morphological types are homogeneously tuned to CA3 theta oscillatory input. A. Neurons for analysis were selected within 100 µm radius of the stratum pyramidale electrode location (top left). Phase locking angle and strength for a range of individual (CA3) cell frequencies (columns) and modulation frequencies (rows). B. Phase Modulation. Spike discharge probability of all neurons grouped by morphological type (left). Phase locked neurons tuning over theta cycle for each morphological class over a single theta cycle (middle). Experimental validation of phase-locking against *in vivo* recordings (right). C. Spiking raster plots. SP_PC cell spiking (top panel); LFP theta rhythm (trace above plot); intereuron spiking (bottom panel). D. Intraceullar traces from morphological cell types (left) and validation against in *in vivo* recordings. Stimulus panels B-D: 0.4 Hz SC mean spiking frequency and 8 Hz signal modulation.

For the same neuronal classes in the model, we compared their somatic membrane potentials and mean firing rates to experimental data. Pyramidal cell spiking was closely aligned to theta LFP rhythm although individual neurons did not spike at every cycle (Figure 7C top). SP_Ivy showed a similar pattern to SP_PC while other types of interneuron participated more sporadically (Figure 7C bottom). Intracellular voltage traces for pyramidal and ivy cells were also similar albeit with ivy cell firing slightly later and overlapping with other types of interneurons (Figure 7D left). Mean firing rates during theta were generally lower than observed *in vivo* except for ivy cells, which were a close match; SP_AA, BS, and BC (CCK+ and PV+) were well below empirical expectations (Figure 7D right). Theta modulated the amplitude of pyramidal cell membrane potential by 1.57-7.34 mV (for 0.1-0.4 Hz SC axon frequency), consistent with the *in vivo* range (2-6 mV, Ylinen et al., 1995). When we compared model population synchrony during theta oscillations with *in vivo* data (Csicsvari et al., 1998), we found that the percentage of SP_PC spiking was a poor match around the theta trough (0°) but was a better match around theta peak (180°), while fast-spiking SP_PVBC and to a lesser degree SP_AA were under-recruited (Figure S24). Overall, for this stimulus the pyramidal-interneuron theta phase order suggests that intrinsic inhibition was activated more powerfully by recurrent than by afferent excitation. Altogether, for the CA3 input, the model does not generate theta at *in vivo*-like calcium levels but does reliably at *in vitro*-like ones. However, the phase analysis suggests that the mechanism is different from what is observed *in vivo*.

##### Medial septum input

*In vivo* evidence points to a fundamental role of the medial septum (MS) in theta generation (Colgin, 2013). To model the MS input, we (i) set an *in vivo* extracellular calcium concentration (1 mM), (ii) applied a tonic depolarizing current (% of rheobase current) to all neurons to represent *in vivo* background activity, (iii) introduced an additional current to mimic the depolarizing effect of an arhythmic release of ACh from the cholinergic projection (see ACh section), and (iv) applied a theta frequency sinusoidal hyperpolarizing current stimulus only to PV+ CA1 neurons to represent the rhythmic disinhibitory action of the GABAergic projection (Hangya et al., 2009; Sun et al., 2014; Müller and Remy, 2018; see Figure 8A). Due to uncertainty regarding some of these factors, we examined the response over a wide range of physiological conditions (see Methods). Since this required a high number of simulations, we decided to use the cylinder circuit.

**Figure 8:**
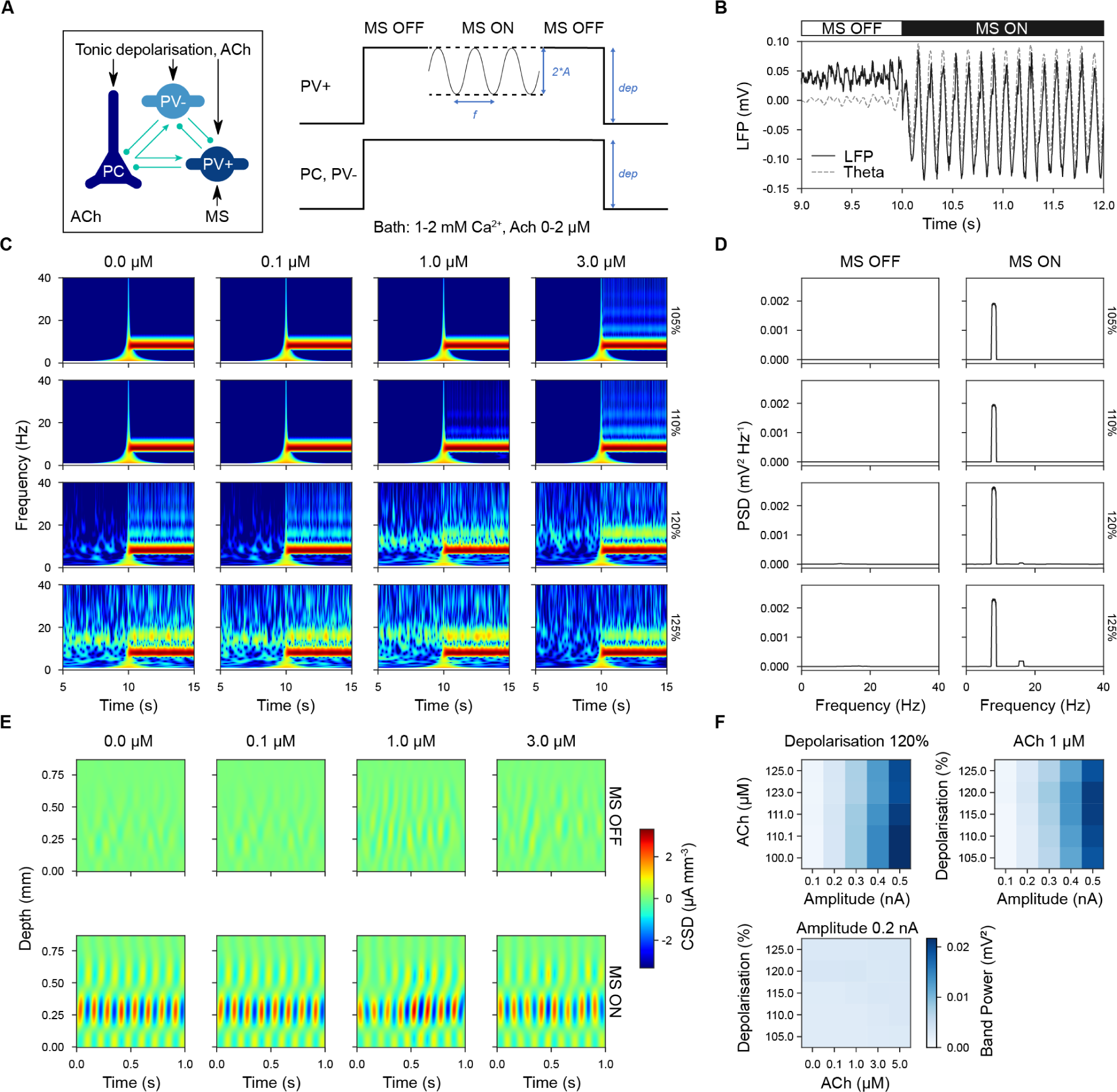
Medial septum (MS) disinhibition of parvalbumin-positive (PV+) interneurons induced theta oscillations in CA1. A. Simulation setup. All neurons received a tonic depolarizing current in the presence of ACh (”MS OFF” condition). For a given period an oscillatory hyperpolarizing current was injected into PV+ interneurons only (”MS ON” condition). B. Example of simulation before and after the onset of disinhibition. LFP in black and theta-filtered LFP in grey. C. Spectrogram for a range of ACh concentrations (top labels) and tonic depolarization levels (right labels). D. Power spectral density (PSD) across different levels of tonic depolarization (right labels) with and without disinhibition. E. Current source density (CSD) analysis across different ACh concentrations (top labels), with and without disinhibition. F. Theta band power as a function of the amplitude of oscillatory hyperpolarizing current, ACh concentration, and level of tonic depolarization. B-E: Stimulus disinhibitory modulation amplitude of 0.2 nA.

Prior to the onset of the disinhibitory stimulus (”MS OFF”), the global tonic depolarization resulted in weak, irregular beta-band LFP activity in CA1 but after its onset (”MS ON”), it induced a strong and sustained, regular theta oscillation matching the frequency of the hyperpolarizing stimulus (Figure 8B). For disinhibitory modulation amplitude of 0.2 nA, the LFP waveforms generated were close to symmetrical (mean asymmetry index = 0.25 *±* 0.11; see Figure S25). Over a range of ACh concentrations and tonic depolarization levels, this theta rhythm was robust, narrow banded (Figure 8B,C), and generated by a highly regular current source restricted to SP (Figure 8E). Higher ACh concentrations, while slightly reducing theta-band power, reduced the level of beta-band activity (Figure 8C). Increased levels of tonic depolarization enhanced both theta harmonics and higher frequency components (Figure 8C,D). We observed that theta-band power was more dependent on the amplitude of the disinhibitory oscillation than on either ACh concentration or tonic depolarization level (Figure 8B,F). Therefore, we found MS-mediated disinhibition could strongly induce CA1 theta oscillations.

During theta oscillations, the phase of spiking of different neuronal classes here divided into two main groups that were in anti-phase with each other (see Figure 9). As the level of tonic depolarization increased, more phase-locked cells were detected (Figure 9A) and only above 110% depolarization (where 100% represents the depolarization necessary to reach spike threshold) were there a sufficient number of active interneurons to discern this dual grouping. Increasing ACh concentration tended to weaken pyramidal phase locking (Figure 9A). For example, at 120% depolarisation and 1 µM ACh, the firing of SP_PC, SP_Ivy, and SP_CCKBC cells was broadly tuned around the theta trough and rising phase, while the firing of SP_AA, BS and SP_PVBC neuron was more narrowly tuned around the peak of the theta rhythm (Figure 9B left). Neurons with significant phase locking matched this pattern but were even more narrowly tuned (Figure 9B middle). The phase locking of SP_AA, SP_Ivy and PC closely matched *in vivo* recordings (see Table S23) but SP_BS and BC (CCK+ and PV+) were by comparison more than 90 degrees out of phase (Figure 9B right).

**Figure 9:**
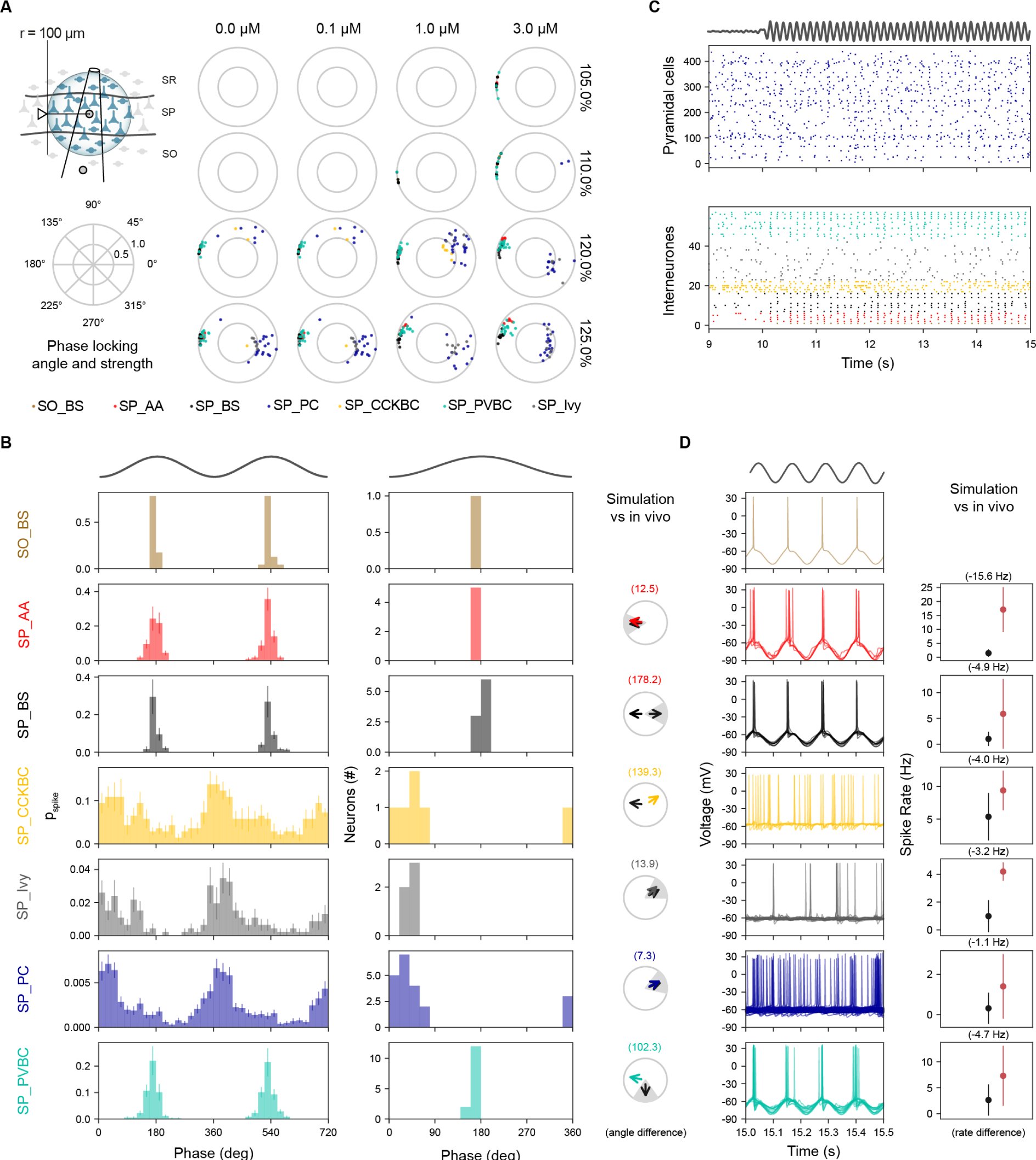
Medial septum (MS) disinhibition induced anti-phase modulation of CA1 neurons during theta cycles. A. Neurons for analysis were selected within 100 µm radius of the stratum pyramidale electrode location (top left). Phase locking angle and strength for range of ACh concentration (columns) and levels of tonic depolarization (rows, where 100% represents the spike threshold) for modulation amplitude of A = 0.2 nA. B. Phase Modulation. Spike discharge probability of all neurons grouped by morphological type (left). Phase-locked neurons tuning over theta cycle for each morphological class over a single theta cycle (middle). Experimental validation of phase locking against *in vivo* recordings (right). C. Spiking raster plots. SP_PC cell spiking (top panel); LFP theta rhythm (trace above plot); interneuron spiking (bottom panel). Disinhibition is switched on (”MS ON”) at time 10s. D. Intracellular traces from morphological cell types (left) and validation against in *in vivo* recordings. B-D: Stimulus disinhibitory modulation amplitude of A = 0.2 nA, 1 µM ACh and tonic depolarisation 120%.

The voltage traces and rate of spiking during theta oscillations for different neuronal types was also grouped. While SP_PC did not spike on every theta cycle their firing appeared weakly modulated by theta (Figure 9C top). Whereas for interneurons, SP_AA, BS, and SP_PVBC spiked tightly for most cycles, ivy cells spiked more rarely and SP_CCKBC more tonically (Figure 9C bottom). Intracellular voltage traces for SP_AA, BS, and SP_PVBC showed they spiked tightly on the rebound from the release of the hyperpolarizing stimulus, whereas SP_PC and other interneurons lacking this were less reactive to theta (Figure 9D left). Notably, all neurons spiked at a lower average rate than *in vivo* recordings (Klausberger et al., 2003, 2004; Klausberger, 2005; Fuentealba et al., 2008) (Figure 9D right). The population synchrony of SP_PC with theta trough was consistent with *in vivo* data (Csicsvari et al., 1998) for a range of disinhibitory stimulus amplitudes whereas for fast-spiking interneurons like SP_AA and SP_PVBC, synchronization with theta peak only occurred with lower stimulus amplitudes (Figure S26). Taken together, the MS-mediated disinhibition entrained theta oscillations under *in vivo*-like conditions, creating a diversity of firing phases between interneuron classes, close to what has been observed experimentally.

In summary, we used the reference model to investigate several possible mechanisms for theta oscillations. For intrinsic mechanisms, we found that spontaneous synaptic release and random afferent synaptic barrage did not induce detectable theta oscillations in the model, while tonic depolarization could induce a variable and unstable theta oscillation at 10-12 Hz. For extrinsic mechanisms, both CA3 and MS input induced a stable and stronger theta oscillation but in different ways, where MS disinhibition was more compatible with *in vivo* data.

#### 2.2.1 Other Frequencies

Next we asked whether the reference circuit was capable of propagating gamma oscillation using the commonly used *in vitro* experimental paradigm of inducing them using bath carbachol (CCh) (Bianchi & Wong, 1994; J. H. Williams & Kauer, 1997; Fisahn et al., 1998; Fellous & Sejnowski, 2000). Specifically, we replicated the setup of Zemankovics et al. (2013), where the authors added CCh to generate oscillations in CA3, which were transmitted to CA1 via SC. The simulation conditions were tailored to these *in vitro* experiments (i.e. 300 µm-thick slices, 2 mM extracellular *Ca*^2+^, 2 mM extracellular *Mg*^2+^, 10 µM ACh). We matched the shape and frequency of the input stimulus reported by Zemankovics et al. (2013) and followed the same LFP analysis methodology. We observed that gamma oscillatory SC input could entrain the entire CA1 network of the model slice to oscillate at the driving frequency (31 Hz) (Figure S27A). As well as inducing oscillations in CA3, *in vitro* CCh could alter the response of CA1 neurons to the SC input. Thus, to quantify the effect of CCh on CA1, we repeated the simulation without the influence of CCh. SC inputs, ranging from 15,000 to 100,000 stimulated fibers, were able to induce strong gamma oscillation in the absence of CCh. However, CCh increased the number of inputs needed for stable gamma oscillation, probably due to its weakening effect on synapses (Figure S27B). Therefore, at least for this experimental setup, the circuit was capable of propagating gamma oscillations.

Finally, we investigated how the reference circuit behaved across a much wider range of input frequencies. Because the nature of the input was less clear across this range, we used a sinusoidal modulated stimulus. In particular, we extended SC input for a wider range of cell (0.1-0.8 Hz) and modulation frequencies (0.5-200 Hz) and measured the corresponding input-output (I-O) gain and spike-spike correlation (Figure 10A). We found I-O gain was not uniform but depended on both cell and signal input frequencies (Figure 10B). The I-O responses of PC and interneurons were different, with interneuron gain greater at lower cell frequencies compared to PCs (Figure 10B). The strongest overall CA1 gain was obtained with a mean CA3 frequency of 0.4 Hz. The spike-spike correlation also depends on both cell and signal input frequencies (Figure 10C-E upper). In the case of input cell frequency of 0.4 Hz, we found CA1 spiking activity was strongly correlated for delta-to lower gamma-band input signal frequencies (i.e. between 1 and 30 Hz) but weaker outside this range. Internal CA1-CA1 spike correlation was similar but spanned higher frequencies of the gamma-band (Figure 10C-E lower). Therefore, the model is able to propagate oscillations better within the delta-to low gamma-band range.

**Figure 10:**
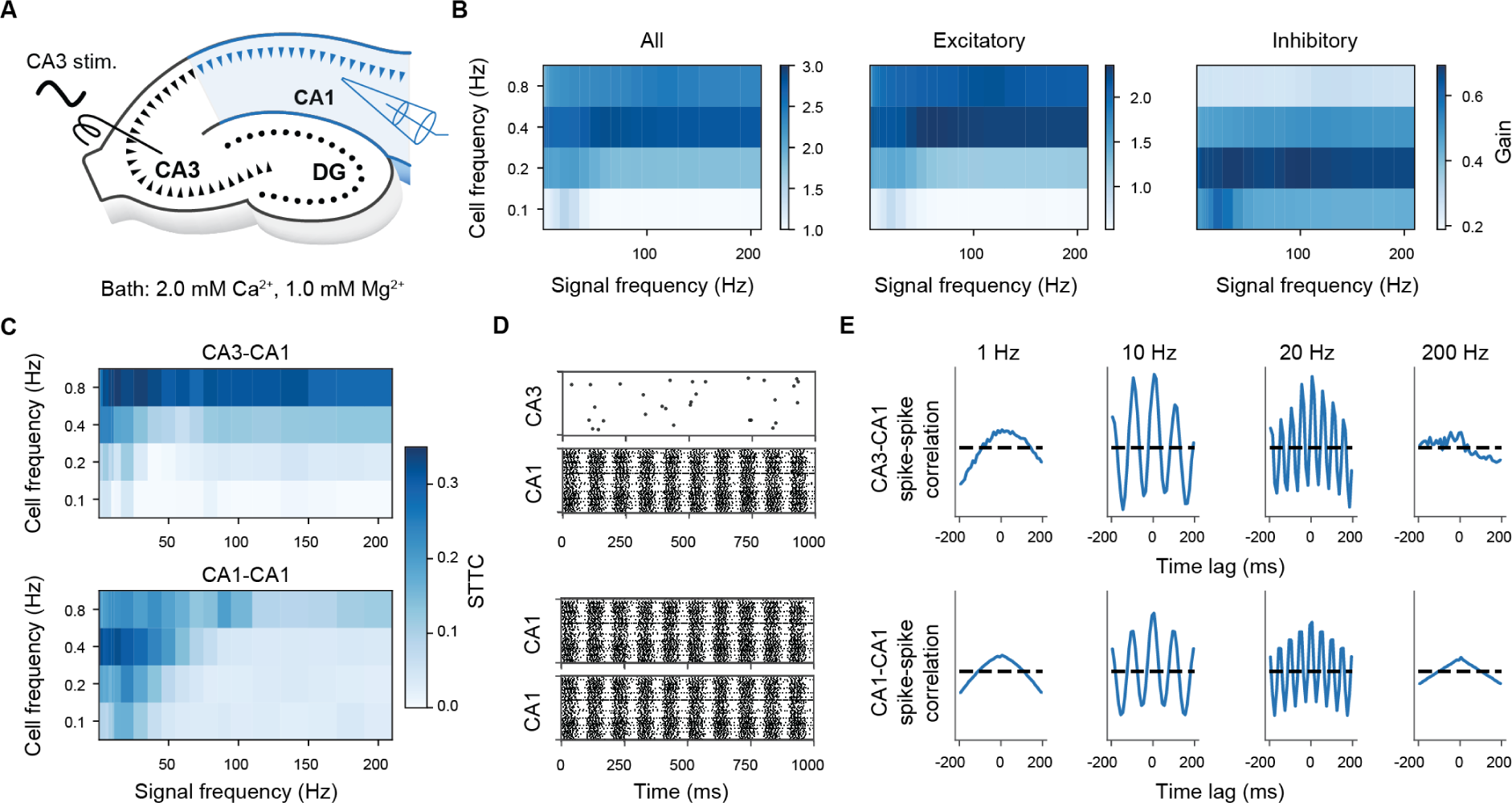
Intermediate frequencies propagate more efficiently through SC. A. *In silico* experimental setup. B. Ratio between the number of CA1 and CA3 spikes as a function of input cell and signal frequencies. We considered all CA1 neurons (left), CA1 PCs (center) and CA1 interneurons (right). C. Heatmaps representing the computed STTC values as a function of input cell and signal frequencies, for CA3-CA1 and CA1-CA1 neurons. D. Examples of CA3 and CA1 spike train (cell frequency of 0.4 Hz, signal frequency of 10 Hz). 100 random CA3 neurons (i.e., SC fibers) and CA1 neurons are selected for clarity. The same neurons are used to compute the STTC and spike-spike correlations (panels D-E). E. Spike-spike normalized correlation histograms for four signal frequencies (cell frequency of 0.4 Hz), for CA3-CA1 and CA1-CA1 neurons.

## 3 Discussion

### 3.1 Main summary

This study presents the reconstruction and simulation of a full-scale, atlas-based reference model of the rat hippocampal CA1 region driven by community data and collaboration. We extended and improved the framework of Markram et al. (2015) to curate and integrate a wide variety of anatomical and physiological experimental data from synaptic to network levels. We then systematically applied multiple validations for each level of the model. We augmented the resulting highly detailed intrinsic CA1 circuit with a reconstruction of its main input from CA3 and a phenomenological model of neuromodulation by acetylcholine. Importantly, the reference circuit model is, by definition, general: it was not created to reproduce a narrow spectrum of use cases but to be capable of addressing a wide range of research questions. To demonstrate its general utility, we were able to simulate different scales of circuits and investigate the generation and transmission of neuronal oscillations, with particular emphasis on theta rhythm, for a variety of stimulus conditions.

### 3.2 Previous work and limitations

While a full review of the many hippocampal circuit models is beyond the scope of the paper, we focus on the progression in both the size and level of detail of large-scale multiscale models of the rat hippocampus during the last three decades (for a comparison of their key features with the present model, see Table S2). These biologically realistic models aim to explain the complex dynamics of hippocampal activity, in particular the generation and control of rhythmic responses. However, all these models, including the one reported here, are incomplete descriptions of a hippocampal region or regions because of the paucity or even absence of some types of data necessary to constrain them. Moreover, the results of these models are difficult to compare because of fundamental differences in their composition, organization, and underlying assumptions. There is no commonly agreed-upon set of validations with which to benchmark a circuit model against experimental data.

The current model stands out by realistically constraining neurons and their connectivity by the highly curved shape of CA1 rather than by relying on an artificial space as found in other contemporary models. Additionally, it reflects both short-term plasticity and spontaneous synaptic release, well-established characteristics of central nervous system synapses. In addition, the morphologies and electrical properties of model neurons here are not just copies of the same class exemplars but their properties have been systematically varied to better capture the diverse nature of neuronal circuits and their responses to stimulation. However, compared with Bezaire et al. (2016), some elements are still missing from the current model such as neurogliaform cells, which did not exist in our available dataset, and GABA*_B_*R which are not included in our simulations.

Nonetheless, the current model includes the perforant path-associated and trilaminar interneurons, which were absent from all previous models. In addition, we modeled the NMDA synaptic currents observed in both hippocampal pyramidal cells and interneurons (with specific NMDAR conductance, rise and decay time constants for each pathway) that are absent in the model of Bezaire et al. (2016). Furthermore, the connectivity algorithm used for the current model generates an intrinsic connectome with more realistic high-order statistics than the more prescriptive approach used in the Bezaire et al. (2016) model (see Giacopelli et al. (2021) for the analysis). Unlike Yu et al. (2020), we did not replicate the topography of the afferent projections, which may play a role in patterning the circuit response, but did model the projections and circuit at a full rather than reduced scale. Overall, further improvement to our model requires additional experimental data.

While we incorporated key features distributed among previous models into a single, general model (see Table S2), it is important to recognise that our aims and approach were different, representing a step change in hippocampal modeling. The intention of the framework was first to curate and integrate community data into the model, preserving provenance for reproducibility, in a way that would allow the addition of new datasets from the large hippocampal community. Re-using these datasets and then making them publicly available through the hippocampushub.eu supports the 3R principles (replace, reduce, refine) for the reduction of animal experiments. Each circuit component and the final model was then systematically validated in an open and transparent way to a degree not previously attempted by other research groups. To increase the realism and utility of simulation experiments, we sought to approximate experimental conditions (e.g. slice thickness and location, bath calcium, magnesium and acetylcholine concentrations, and recording temperature) and to increase the capability to manipulate and record from the model (e.g. enable spontaneous synaptic release, alter connectivity, estimate extracellular LFP signals, and apply a variety of stimuli). In short, the aim was to offer a more realistic yet scalable and sustainable approach to model brain regions at full scale.

### 3.3 Lessons learned

In the context of a community effort, the process of curating and integrating available data to reconstruct a brain region and replicating the experimental conditions *in silico* proved instructive in a number of ways. First, assembling the components to reconstruct a brain region reveals surprising gaps in the available data and knowledge. Notably, for instance, while Schaffer collateral input to CA1 has received many decades of attention, especially in terms of long-term plasticity, we found the basic information needed to model this pathway quantitatively, was limited. To address this gap, we devised a multi-step algorithm constrained by the data that were available to parameterize these connections. Second, when data was available it often required further work before it could be used in the model. While an open-source rat hippocampal atlas (Ropireddy et al., 2012) was crucial to reconstruct CA1, the original volumetric reconstruction was too noisy for our purposes and required additional processing to give smooth layering. This smoothness was necessary to place and orient morphologies accurately in the atlas in relation to the layers. If the morphology was incorrectly placed or oriented, this had a knock-on effect for how the circuit was connected. Similarly, the completeness of morphological reconstructions also affected connectivity. For these reasons, some cell types in our available dataset could not be used in the circuit model, sacrificing a small amount of cell type diversity in favor of completeness. Third, setting up simulations to reproduce the desired experimental conditions required careful attention. We offer two examples from our research. When reproducing the I-O gain of SC afferent input reported in Sasaki et al. (2006), we initially sampled all neurons in the model slice to plot to the I-O curve. However, the result was poor. We later resolved this by following their experimental sampling of a subset of neurons with which we could closely match the empirical curve. When reproducing MS-induced theta oscillations, we initially simulated under default conditions of extracellular calcium concentration at 2 mM, resulting in theta oscillations that occurred episodically and only for a restricted parameter regime. However, when we lowered the extracellular calcium to *in vivo* levels (1 mM), sparser activity led to more robust and stable theta oscillations.

### 3.4 A community-driven modeling approach

This approach offers important features that make an attractive case for adoption by the wider hippocampus community. First, anchoring a circuit model in the volumetric space of a brain region atlas makes mapping experimental data for data integration, validation, and prediction easier than for more abstract spaces and permits investigations at different scales, e.g. a brain slice cut at an arbitrary angle. The Allen Brain Atlas has demonstrated the advantages of registering community experimental data in a common reference atlas (Wang et al., 2020). Extending this principle, a common framework appears advantageous for modeling as well. Second, the model components, validations, and circuit are openly available through a dedicated portal (hippocampushub.eu) to maximize transparency and reproducibility. We allow the community to examine how the reference circuit model was built, validated, simulated, and analyzed. We conceived the model to be adopted by the community, extended and improved. Third, while the traditional approach of constructing circuits for a specific use-case has short-term advantages for demonstrating proof-of-concept, in the long-term a community reference model must be valid across a range of use-cases. Finally, this circuit model can be extended to incorporate glia and vascular systems within the same framework (Zisis et al., 2021). These systems are fundamental to regulating neuronal activity and communication in health and disease (Giaume et al., 2010) and could be adapted to make atlas-based circuit models much more realistic embodiment of brain regions.

## 4 Methods

### 4.1 Experimental procedures

In this study, we used two datasets for single neuron morphological reconstructions and electrophysiological recordings from the region CA1: Sprague Dawley rat and Wistar rat. Both datasets also include the reconstruction of the layers that were used to estimate the layer thicknesses (see Layers Section) and guide the cell placement (see Soma placement Section). Note that, while the Wistar rat dataset also includes electrophysiological recordings, we did not use them for this model. The experimental procedures have been previously described in R. Migliore et al. (2018), but are summarized below.

#### 4.1.1 Sprague Dawley rat dataset

All procedures were carried out according to the British Home Office regulations (Animal Scientific Procedures Act 1986). Hippocampal slices were obtained from young adult rats (Sprague Dawley, 90-180g) as previously described (Ali et al., 1998; Ali et al., 1999; Pawelzik et al., 1999; Hughes et al., 2000; Thomson et al., 2000; Pawelzik et al., 2002; Mercer et al., 2006; Ali & Thomson, 2008; Fuentealba et al., 2008). Briefly, following deep anesthesia with euthatal solution, young adult rats (Sprague Dawley, 90-180 g) rats were perfused transcardially with ice-cold modified artificial cerebrospinal fluid) containing in mM: 248 sucrose, 25.5 *NaHCO*_3_, 3.3 KCl, 1.2 *KH*_2_*PO*_4_, 1 *MgSO*_4_, 2.5 *CaCl*_2_, 15 D-glucose, equilibrated with 95% *O*_2_/5% *CO*_2_. 450 to 500 µm coronal sections were cut, transferred to an interface recording chamber and maintained at 34 *−* 36*^◦^*C in modified ACSF solution for 1 hour and then in standard ASCF (in mM: 124 NaCl, 25.5 *NaHCO*_3_, 3.3 KCl, 1.2 *KH*_2_*PO*_4_, 1 *MgSO*_4_, 2.5 *CaCl*_2_, and 15 D-glucose, equilibrated with 95% *O*_2_/5% *CO*_2_) for another hour prior to starting electrophysiological recordings. Single intracellular recordings were made using sharp microelectrodes (tip resistance, 90-190 MΩ) filled with 2% biocytin in 2M *KMeSO*_4_ under current-clamp (Axoprobe; Molecular Devices, Palo Alto, CA). Electrophysiological characteristics of the recorded neurons were obtained from voltage responses to 400 ms current pulses between -1 and +0.8 nA and recorded with pClamp software (Axon Instruments, USA).

Following recording and biocytin filling, slices of 450-500 µm were fixed overnight [4% paraformaldehyde (PFA), 0.2% saturated picric acid solution, 0.025% glutaraldehyde solution in 0.1 M phosphate buffer)] as previously described (Hughes et al., 2000; Economides et al., 2018). Slices were then cut in 50-60 µm sections, cryoprotected with sucrose, freeze-thawed, incubated in 1% *H*_2_*O*_2_ and then in 1% sodium borohydride (NaBH4). Slices were incubated overnight in ABC (Vector laboratories) and then in DAB (3, 3’ diaminobenzidine, Sigma) to reveal the morphology of the recorded neurons. Following washes, sections were post-fixed in osmium tetroxide (*OsO*_4_), dehydrated, cleared with propylene oxide, mounted on slides (Durcupan epoxy resin, Sigma) and cured for 48 h at 56°C. For immunofluorescence staining, slices were incubated in 10% normal goat serum after incubation with *N aBH*_4_. Sections were then incubated overnight in a primary antibody solution containing mouse anti-PV (Sigma), rabbit anti-PV (Baimbridge & Miller, 1982) or mouse anti-gastrin/CCK (CURE, UCLA) made up in ABC and then in a solution of secondary antibodies (Avidin-AMCA, FITC and anti-rabbit Texas Red) for 3 hours. Following fluorescence visualization, slices were incubated in ABC, DAB and *OsO*_4_ prior to dehydration and mounting as described above. All neurons were then reconstructed in 3D using a Neurolucida software (MBF Bioscience) and a 100X objective as previously described (Economides et al., 2018).

#### 4.1.1 Wistar rat dataset

The project was approved by the Swiss Cantonal Veterinary Office following its ethical review by the State Committee for Animal Experimentation. All procedures were conducted in conformity with the Swiss Welfare Act and the Swiss National Institutional Guidelines on Animal Experimentation for the ethical use of animals. Hippocampal slices were obtained from young adult rats (Wistar, postnatal 14-23 days) as previously described (Markram et al., 2015; R. Migliore et al., 2018). In brief, *ex vivo* coronal preparations (300 µm thick) were cut in ice-cold aCSF (artificial cerebro-spinal fluid) with low *Ca*^2+^ and high *Mg*^2+^. The intracellular pipette solution contained (in mM) 110 potassium gluconate, 10 KCl, 4 ATP-Mg, 10 phosphocreatine, 0.3 GTP, 10 HEPES and 13 biocytin, adjusted to 290 *±* 300 mOsm/Lt with D-mannitol (2 *±* 35 mM) at pH 7.3. Chemicals were from Sigma Aldrich (Stenheim, Germany) or Merck (Darmstadt, Germany). A few somatic whole cell recordings (not used for this model) were performed with Axopatch 200B amplifiers in current clamp mode at 34 *±* 1 *^◦^*C bath temperature.

The 3D reconstructions of biocytin-stained cell morphologies were obtained from whole-cell patchclamp experiments on 300 *µ*m thick brain slices, following experimental and post-processing procedures as previously described (Markram et al., 1997). The neurons that were chosen for 3D reconstruction were high contrast, completely stained, and had few cut arbors. Reconstruction used the Neurolucida system (MicroBrightField Inc., USA) and a bright-field light microscope (DM-6B, Olympus, Germany) at a magnification of 100x (oil immersion objective, 1.4-0.7 NA). The finest line traced at the 100x magnification with the Neurolucida program was 0.15 *µ*m. The slice shrinkage due to the staining procedure was approximately 25% in thickness (Z-axis). Only the shrinkage of thickness was corrected at the time of reconstruction.

### 4.2 Morphological classification

We classified the morphologies into one of 12 different morphological types (m-types) based on the layer containing their somata and their morphological features. For the classification, we adopted three main assumptions. 1) Several subtypes of PCs have been described (Baimbridge & Miller, 1982; Bannister & Larkman, 1995; Deguchi et al., 2011; Mizuseki et al., 2011; Slomianka et al., 2011; Graves et al., 2012; Lee et al., 2014; Malik et al., 2016), but for simplicity we consider the pyramidal cells as a uniform class (SP_PC or simply PC). 2) SP_PVBC and SP_CCKBC were classified as two separate m-types. The two types of basket cells can be distinguished by biochemical markers and electrical properties, and they show different densities and connectivity (Bezaire & Soltesz, 2013). On the contrary, there is no strong evidence for differences in their morphologies and the small number of examples in our possession (3 SP_PVBCs, 1 SP_CCKBCs) prevented us from conducting any systematic classification. While we could have pulled the two cell types into one m-type, we kept them separated for the sake of simplicity. 3) We created supersets of m-types based on their principal biochemical marker. In particular, we defined PV+ cells as SP_PVBC, SP_BS, SP_AA, CCK+ cells as SP_CCKBC, SR_SCA, SLM_PPA and SOM+ (somatostatin) cells as SO_OLM, SO_BS, SO_BP, SO_Tri. When the layer is not specified (e.g., BS), we meant all the neurons of this type regardless their soma location.

### 4.3 Morphology curation

We curated the morphological reconstructions extensively prior to insertion in the network model. First, we translated the reconstructions to have their somas centered at coordinates (0,0,0). We reoriented the reconstructions so that their x, y, and z axes coincided respectively with the transverse, radial, and longitudinal axes they followed in the tissue. We considered the cells to be substantially complete, with only a few cuts, so we have not applied any corrections for cuts (Markram et al., 2015). The only exception is the PC axon. One particular reconstruction has a relatively long axon (3,646 µm) that spans 1,325 µm along the transverse axis. We assumed this axon to be relatively complete within the CA1 and we used it as a prototype for all the PCs. Validation of the number of synapses per connection, bouton density, and connection probability showed that this assumption is reasonable (see Building CA1 Section). A subsequent cloning procedure (see Section Morphology library Section) guaranteed that all the PC axons are unique. We removed two pyramidal cells due to the complete absence of an axon since at least the axon initial segment (AIS) needs to be present to create electrical models.

We applied a series of corrections as described in Markram et al. (2015). In brief, to comply with the NEURON and CoreNEURON standards, we repaired the reconstructed morphologies using NeuroR (Anwar et al., 2009, Table 1). In particular, we used this tool to remove unifurcations of dendrites, fix neurites whose type changes along its main branch, fix invalid formats of soma, and remove segments with close to zero length. We used this tool also to correct tissue shrinkage. The correction increased the neurite reach approximately 25% along the z-axis and approximately 10% along the x- and y-axes.

### 4.4 Morphology library

To produce morphology variants that fit well in different locations of the space, we applied scaling *±*15% of the original reconstructions. This range was a good compromise between the need to change the size but yet to not introduce too much distortion (Markram et al., 2015). To increase morphological diversity, we applied a cloning procedure as described in Markram et al. (2015). Some clones showed a wrong distribution of axons and dendrites, and we removed them. In particular, we annotated how different parts of the morphology were positioned within the layers (see Layers Section). We used these annotations to guide the cell placement, but it was also useful to discard cells that were too distorted by cloning and had an incorrect distribution of neurons within the layers. Clones were further validated by visual inspections. For OLM cells, a scaling of *±*15% was not sufficient to make sure that the axon correctly targeted SLM in all dorso-ventral positions. In this case, we introduced a synthetic axon. In the reconstruction of OLM cell, a chunk of 60 µm axon, which is placed at 85 to 145 µm from the soma, falls in SR and it is relatively simple. To produce morphologies with about 90% to 150% (step of 5%) of the height of the original morphology, we removed the aforementioned chunk of 60 µm axon (93%), only 45 µm of it (95%), or added a multiple of 45 µm (corresponding to a step of 5%) synthetic axon within SR until we reached a 150% scaling.

After we corrected the orientation of the initial set of reconstructions, the cloning procedure changed the branch lengths and rotations, which may have resulted in a change in the main orientation of the cells. Based on visual inspection, the effect on short neurites was negligible. In the case of PCs, the axon is relatively longer and has a clear orientation (S. Yang et al., 2014). In this case, the cloning may significantly alter the orientation of the axon, and this may result in a wrong placement in the circuit (see Soma placement Section). To avoid this problem, we renormalized the orientation of the cloned PCs. In particular, we used principal component analysis (PCA) to first determine the principal axis of all axonal points in each cloned morphology. Next, we used the direction of the principal axis to rotate the caudal portion of the arbor onto the x-axis so all pyramidal axon arbors were aligned in the same direction. Finally, to remove an unrealistic degree of variability, we filtered out cloned morphology outliers whose width (z-axis range) was 3.3 times greater than that of the original pyramidal axon arbor from which they were derived. This filtering step rejected about 14% of all pyramidal cell cloned morphologies.

### 4.5 Morphology library validation

Neuronal morphologies were validated at the end of the processing stage (cloning) by comparison of their morphological properties, as well as their topological profiles. Around 100 morphological features from dendritic and axonal neurites were extracted using the NeuroM package (Palacios et al., 2022). The median over visible spread (MVS) statistical score, defined as the ratio of the absolute value of the difference between the medians of the feature 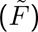 divided by the overall visible spread (OVS), was computed according to the following formula:

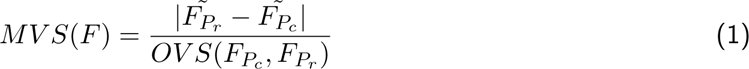

to identify the difference between populations of reconstructed *P_r_* and cloned *P_c_* neurons for each feature *F*. This score is close to zero for similar populations and increases as the difference between the populations increases. Scores below 0.3 indicate good agreement between the two populations as their differences are contained within three standard deviations (STD). Note that this measurement depends on the sample size, due to the computation of the overall visible spread, which remains small if the sample size is small and therefore the MVS score increases accordingly.

In addition to the traditional morphometrics, the topological profiles of the original reconstructions were compared to those of the processed morphologies (repaired and cloned). The topological morphological descriptor (TMD) encodes the start and the end radial distances of each branch from the soma surface (Kanari et al., 2019). The pairs of distances (start, end) are represented in a two dimensional plane, a representation known as the persistence diagram of the neurons (Figure S4). The Gaussian kernels of these points are averaged to generate the persistence images (Figures S5 and S6). The persistence images of two populations can be subtracted to study the precise differences between datasets. Note that the robustness of average persistence images depends on the sample size of the population, when only few (less than three) cells are available these images represent a small subset of the biological population and are therefore not quantitatively reliable.

### 4.6 Electrical type classification

We classified the cells into different electrical types (e-types) based on their firing patterns as defined by the Petilla nomenclature (Petilla Interneuron Nomenclature Group et al., 2008). Some of the firing patterns (i.e., irregular patterns) were too rare and we excluded them.

### 4.7 Morpho-electrical compositions

The 154 recordings were recorded from different m-types, and we observed that each m-type can show one or more firing patterns with different probabilities. We used this information to derive the morpho-electrical type (me-type) composition (Table S4).

### 4.8 Ion channels

We considered a set of active membrane properties which included a voltage-gated transient sodium current, four types of potassium current (DR-, A-, M-, and D-type), three calcium channels (CaN, CaL and CaT), a nonspecific hyperpolarization-activated cation current (Ih), two Ca-dependent potassium channels (KCa and Cagk), and a simple *Ca*^2+^-extrusion mechanism with a 100 ms time constant. We based channels’ kinetics on previously implemented cell models of hippocampal neurons (M. Migliore et al., 1999; M. Migliore et al., 2005; Ascoli et al., 2010; Morse et al., 2010).

### 4.9 Single neuron modeling

Single cell models are described in previous publications (R. Migliore et al., 2018; Ecker et al., 2020; Romani et al., 2022). In brief, we optimized cell parameters to match electrophysiological features rather than the traces directly. We extracted features using the open-source Electrophysiological Feature Extraction Library (eFEL, Table 1) or the Blue Brain Python E-feature extraction (BluePyEfe, Table 1). Initial optimizations considered only features from somatic recordings (R. Migliore et al., 2018), while a subsequent refinement included also the amplitude of the back-propagating action potential (in the apical trunk, 150 and 250 µm from the soma) for PCs (Ecker et al., 2020).

We assumed channels were uniformly distributed in all dendritic compartments except *K_A_*and *I_h_*, which in pyramidal cells are known to increase with distance from the soma (Hoffman & Johnston, 1999; Magee, 1999). Pyramidal cells include *K_M_* in the soma and axon (Shah et al., 2008), *K_A_* with different kinetics in dendrites, soma and axon (Hoffman et al., 1997; M. Migliore et al., 1999), *K_M_* with a different kinetics in the soma and the axon, Na and *K_DR_*, whereas they do not include *K_D_* since the delayed spiking is not a feature observed in PCs. Given the limited knowledge of the currents in interneurons, we applied the same currents of pyramidal cells with the following exceptions. Dendritic sodium channel densities decay exponentially with distance from the soma (with a length constant of 50 µm) based on Hu et al. (2010). We included *K_D_* since some interneurons show delayed firing. *K_A_* has the same kinetics in somas and dendrites because there is no experimental evidence of a different *K_A_* kinetics in the dendrites of interneurons. We distinguished two types of *K_A_* for proximal and distal dendrites (M. Migliore et al., 1999). For both pyramidal cells and interneurons, we optimized channel peak conductance independently in the different regions of a neuron (soma, axon, and dendrites).

We performed a multi-objective evolutionary optimization using the open source Blue Brain python optimization library (BluePyOpt, Table 1) (Van Geit et al., 2016) to obtain 39 single cell models. From this set, we excluded three models because their me-types are not used in the network model (cAC_SP_Ivy, cAC_SP_PVBC, cNAC_SR_SCA).

### 4.10 Single neuron model validations

We used HippoUnit (Sáray et al. (2021), Table 1) to validate the electrical models of pyramidal neurons, specifically considering the attenuation of PSP and BPAP.

#### 4.10.1 PSP attenuation

The PSP attenuation test evaluates how much PSP amplitude attenuates as it propagates from different dendritic locations to the soma. To get the behavior from the model, EPSC-like currents (i.e., double exponentials with rise time constant of 0.1 ms, decay time constant of 3 ms, and peak amplitude of 30 pA) were injected into the apical trunk of PCs at varying distances from the soma (100, 200, 300 *±* 50 *µm*), and PSP amplitudes were simultaneously measured at the local site of the injection and in the soma. Finally, the experimental and model data points were fitted using a simple exponential function. The space constants resulting from the fitting were then reported and compared with experimentally observed data in Magee and Cook (2000).

#### 4.10.1 BPAP attenuation

The BPAP test evaluates the strength of AP back-propagation in the apical trunk at locations of different distances from the soma. The AP is triggered by a step-current of 1 s with an amplitude for which the soma fires at *∼*15 Hz. The values were then averaged over distances of 50, 150, 250, 350 *±* 20 *µm* from the soma. We measured the amplitudes of the first AP at the four different dendritic locations and compared them with experimentally observed data in Golding et al. (2001).

### 4.11 Library of neuron models

Optimizing all the neurons in the morphology library is computationally expensive. Following Markram et al. (2015), we minimized the problem by using Blue Brain Python Cell Model Management (BluePyMM, Table 1) which combines the morphology library with initial single cell models to produce a library of cell models. The procedure accepts a new combination if the model and experimental features are within five STD of the experimental feature. If at least one feature is greater than this range, then the new combination is excluded. This strategy also has the advantage of increasing the model variability within the experimental data. In the case of bAC-SLM_PPA, the threshold for accepting new combinations had to be relaxed to 12 STD to obtain some valid models.

### 4.12 Rheobase estimation

For all the single cell model combinations, we estimated their rheobase with a bisection search until an accuracy of 10*^−^*^3^ nA was reached. The spikes were recorded at the axon initial segment, and the upper bound from the last step of the search was used, to ensure cells spiked in the axon at rheobase.

### 4.13 Atlas

#### 4.13.1 Volume

We based our annotated volume on a publicly available atlas of the CA1 region (Ropireddy et al., 2012) (http://cng.gmu.edu/hippocampus3d/). From the original atlas, we took all voxels labelled as CA1 without maintaining their subdivisions in CA1a, CA1b, and CA1c or the four layers (Figure S8A). The original file was converted from a .csv to a .nrrd format using voxcell (Table 1).

We undertook a series of post-processing steps to augment the atlas with a coordinate system and vector fields that followed the three hippocampal axes (longitudinal, transverse, radial). First, a substantial smoothing operation was necessary to render the surfaces within the CA1 regular enough for subsequent manipulations. To obtain smoother surfaces, we applied a Gaussian filter together with a morphological closing filter. A few manual checks were needed to ensure isolated voxels were removed and holes from missing tissue closed. The resulting volume has a minor difference in voxel counts compared with the original atlas (−3.7%) (Figure S8B).

We created a mesh for the boundary surface of the CA1 using Ultraliser (Table 1) and separated it into an upper and lower shell (Figure S8C). In practice, the operation of separating the mesh into two portions is challenging and we were not able to automate it. The curved structure of the hippocampus makes automatic solutions appear incorrect upon visual inspection, particularly around the ridges. Taking the results into account, we used an OpenGL 3D graphical user interface tool (atlas-direction-vectors, see Table 1) that allows one to paint voxels on the surface of the atlas in different colors. It enabled us to manually correct the voxel selection until the shell division appeared satisfactory. Two surface meshes were then derived from the selected voxel masks.

We generated a polygonal centerline along the innermost voxels of the CA1 from the dorsal to the ventral extremity (Figure S8D). In detail, this process started by computing the distance transform of the input volume by assigning each voxel the distance from its closest neighbor outside of the volume. The two extremity points were used as entry and exit locations to build a stochastic chain of points following the local maxima of the distance transform. A further step generated a graph of these possible points using proximity conditions and determined the overall shortest path between the extremities using a weighted Dijkstra algorithm (Sniedovich, 2006). The resulting skeleton of the centerline was then converted into a Bézier curve (Agoston, 2004) and its continuous derivative was used to orient a series of planes. These planes are oriented perpendicularly to the curve and cut the volume of the hippocampus in slices at regular intervals (Figure S8E).

#### 4.13.1 Coordinates system

To utilise the 3D volume fully, we created a set of parametric coordinates ranging from 0 to 1 and named them *l*, *t*, *r* as they follow the longitudinal, transverse and radial axes of the hippocampus (Figure S8G). The longitudinal coordinate was assigned first using the derivative of the centerline to sample the volume with many cross-section planes and assigning each CA1 voxel the [0, 1] value of the plane closest to it.

The transverse coordinate was obtained by considering the intersection between each of the cross-section planes and the meshes assigned previously as the upper and lower shells of the volume. Each plane cuts the meshes creating two lines of points which were fitted to spline functions and re-sampled to yield the same number of points each having a [0, 1] *u* coordinate. Connecting the upper to the lower line points resulted in a field of vectors that represent the natural orientation of pyramidal neurons in CA1. The transverse coordinate *t* was assigned from the *u* value of the spline points to all voxels found on the plane according to which was their closest vector (i.e. the line segment joining the upper *usp*(*u*) and lower *lsp*(*u*) line points with the same *u*).

The radial coordinate was assigned following the previous step and represents the relative [0, 1] location of each voxel of the plane along the segment connecting the *usp*(*u*) and *lsp*(*u*) points. It is defined as the ratio:

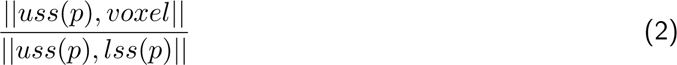

In summary, all voxels lying on the same *l* plane are assigned a *t* coordinate from the upper and lower splines and later those voxels on the same *t* segment are assigned a relative *r* position along it. Taken together, these three coordinates provide a coherent system to slice and parse the volume which is more robust to surface irregularities compared to other methods. Finally, for each voxel we re-computed its direction vector from the partial derivative of the (*l*, *t*, *r*) coordinates with respect to *r* to ensure perfect correspondence between the vector field and coordinate space (Figure S8H).

#### 4.13.1 Layers

Having achieved the necessary smoothing and orientations over the entire volume, we reintroduced the distinction into layers (Figure S8F) which are used to constrain cell density and the placement of specific portions of neural morphologies such as the axon and the apical dendritic tuft (Figure S9). For this reason, we used a combined approach to determine layer position: extracting thicknesses from morphology reconstructions and comparing the final layer volumes with voxel counts from the original atlas. During morphology reconstruction, we traced layer thicknesses from 23 images of pyramidal cells in CA1 slices and averaged them: SLM: 146 *±* 27 µm, SR: 279 *±* 40 µm, SP: 59 *±* 15 µm, SO: 168 *±* 30 µm.

The available morphological reconstructions came from the dorsal CA1, and thus these layer thicknesses reflect the dorsal portion rather than the entire extent of the region. To reintroduce layer labels back into the atlas, we used the radial coordinate computed above to assign a uniform proportion of voxels to each layer (SLM: 0.224, SR: 0.42791, SP: 0.090, SO: 0.258). This assumption returns the same relative layer thickness throughout the volume, automatically compensating for the different CA1 shape in each location thanks to the [0, 1] parametric range. As a final step, we computed the total volumes for each layer (SLM: 3.2143, SR: 6.8421, SP: 1.5789, SO: 4.8178 mm^3^).

### 4.14 Cell composition

We started by fixing the pyramidal cell (PC) density in SP to 264.0 *±* 32.6 *×* 10^3^ mm^3^ (Aika et al., 1994). By combining the PC density and SP volume as extracted from the atlas (1.5789 mm^3^), we estimated the total number of PCs as 416,842. To estimate the number of cells or cell densities for interneurons, we used the ratio between PCs and the different interneurons as predicted by Bezaire and Soltesz (2013) with several assumptions. In some cases, a cell type has been described as present in several layers. If we only have reconstructions of cells from one layer, we consider all the expected cells to be placed in this layer. In the case of bistratified (BS) cells, we have reconstructions from SO and SP, but Bezaire and Soltesz (2013) also described a small percentage of cells in SR. In this case, we considered the cells from SR to be placed in SP. Bezaire and Soltesz (2013) gave a rough estimation of 1,400 cells for trilaminar cells and radiatum-retrohippocampal cells together in SO. Without any other information, we considered these two cell types to have the same proportion, so we estimated the number of trilaminar cells as 700. We ignored the m-types for which we do not have at least one reconstruction. The lack of some cell types leads to a smaller number of interneurons. To maintain the same E/I ratio of 89:11 (Bezaire & Soltesz, 2013), we increased the number of interneurons accordingly. The resulting cell composition counts and densities are shown in Table S3.

### 4.15 Cell placement

#### 4.15.1 Soma placement

We expanded the basic algorithm of Markram et al. (2015) to take into account the more complex volume of the atlas and the particular constraints of the hippocampus. For each me-type, the cell positions are created given cell density volumetric data (using uniform distribution). The total cell count is calculated based on cell density values and the volume. Each voxel is populated with the desired count of cells, weighted by the contribution of that voxel to the total density so that the total count of cells is reached. For SR_SCA and SLM_PPA, we could not select morphologies for all the potential positions within the corresponding layers. For this reason, the soma placement was restricted to two narrow subvolumes within the layers. In the case of SCA, we placed the soma in the middle of SR *±* 5% of the layer thickness. In the case of PPA, we placed the cells in the lower part of the layer, from 2% to 10% of the layer.

#### 4.15.1 Cell orientation

Once we decided the soma positions, we oriented the neuronal morphologies to follow the curvature of the hippocampus and other additional constraints. For all the cells, we aligned their y-axis with the radial axis of the hippocampus, and this allowed us to compute the placement scores and select the morphologies that best fit the space. Cells may show a preferential orientation around the radial axis. For interneurons, the experimental evidence for a specific orientation in the transverse-longitudinal plane is scarce, and we applied a random rotation around the radial axis to avoid any bias in the orientation. On the contrary, literature has reported a particular orientation for pyramidal cell axons. Pyramidal cells normally have two main branches, roughly parallel to the transverse axis, one towards the subiculum and one towards the CA3. There is also a thinner branch that is roughly parallel to the longitudinal axis (Knowles & Schwartzkroin, 1981; S. Yang et al., 2014). To take this into account, we rotated the pyramidal cells so that their axons were parallel to the transverse axis and the most complex branch points closer to subiculum.

#### 4.15.1 Morphology selection

Once we identified the cell positions, we selected the morphologies that most closely matched a set of rules to ensure correct neurite targeting. We distinguished two types of rules: strict and optional ones (see later in this section). Optional rules are shown in Table S6, while we used only one strict rule, i.e., dendrites and axons should be below the top of SLM.

For each soma position, we have a candidate pool of morphologies to be selected. The candidate pool is computed as follows. For each soma position, we consider the radial axis passing through it and compute the relative position of the soma and the layer boundaries. For each morphology we associate a score 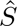 that describes how well the rules are matched. If 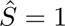 we have a perfect match, 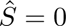 the rules are not matched and the morphology is excluded from the pool. The score combines a score for ‘optional’ rules 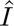 and ‘strict’ rules 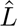

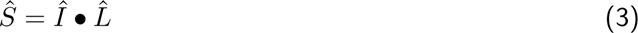

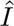 combined all the scores from the *n* ‘optional’ rules *I_j_*as follows:

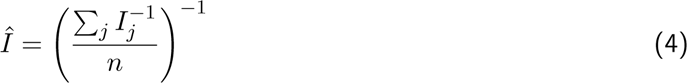

The use of a harmonic mean allows us to penalize low scores for a particular rule heavier than a simple mean, but still “to give it a chance” if other interval scores are high. If some optional score is close to zero (< +0.001), the aggregated optional score would be zero. If there are no optional scores or if optional scores are ignored 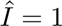.

Each rule *I_j_*is computed as follows:

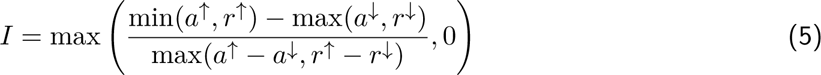

Where 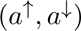 is the interval of the dendrites, axon or dendritic tuft (when applicable) corresponding to the rules. It corresponds to the interval as defined above that needs to be shifted by the soma position *y*_0_ of the morphology in the circuit

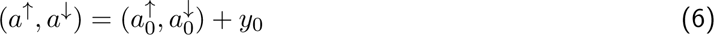

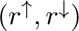 is the interval of the target region along the radial axis passing through *y*_0_. This interval corresponds to the rules expressed as relative intervals in the table above, converted to µm.

The numerator represents the overlap of the two intervals, while the denominator normalizes the overlap by the largest interval among 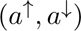 and 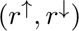. In this way, only a perfect overlap receives a score of 1, while if one interval is larger than the other, the score is lower than 1.

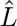 combines all the scores from the ‘strict’ rules *L_k_*as follows:

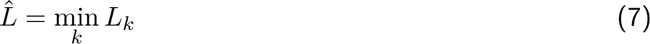

If there are no strict scores *L* = 1.

Given the multiple constraints described above, 2.6% soma positions could not have any associated morphologies, and the were ignored. This resulted fewer neurons than expected. However, validation of the cell composition and densities reassured us that this discrepancy is not significant. In fact, lower densities occurred mainly at the border of the circuit where the particular distorted shape of the layers makes the cell placement more complicated.

### 4.16 Circuit sections

We defined a series of sections of the circuit to be used in analyses and simulations. In particular, we defined nine cylinders equally spanned from position 0.1 to 0.9 along the longitudinal axis, in the middle of the transverse axis (0.5) and with a radius of 300 µm. In addition, we defined 47 consecutive non-overlapping transverse slices of thickness of 300 µm.

### 4.17 Synapses

#### 4.17.1 Local synapse anatomy

To derive the connectome, we adapted the algorithm described in Reimann et al. (2015). The first part of the algorithm finds all potential synapses (appositions or touches) among cells based on colocalization of presynaptic axon and postsynaptic cells. We allowed all the possible connections between m-types with the exception of AA cells that can form contacts only with PC. We allow synapses to occur on soma and dendrites, with the following two exceptions: PC-PC connections only occur on dendrites (Megias et al., 2001) and AA cells make synapses only on AIS of PCs (Freund & Buzsáki, 1996). Initially, SCA, Ivy, and BS were found to make too many synapses on the PC soma compared to the data reported in literature (Freund & Buzsáki, 1996; Vida et al., 1998; Klausberger, 2005; Fuentealba et al., 2008). Since the number of expected synapses on the soma is relatively low, and since the tool does not allow us to prune synapses on specific compartments, we decided not to allow SCA, Ivy, and BS to make synapses on PC somas.

An apposition occurs when the presynaptic axon is within a threshold distance (maximum touch distance) from the postsynaptic cell. In the somatosensory cortex microcircuit, this distance was set to 2.5 µm and 0.5 µm for synapses on PCs and INTs respectively (Markram et al., 2015). However, when we applied the same thresholds to the hippocampus, we could not match some of the experimental data. In particular, compared to experimental data there were too many connections among PCs and too few between PCs and INTs (Takács et al., 2012). Using a grid search approach, we found that maximum touch distances of 1.0 µm and 6.0 µm for synapses on PCs and INTs respectively represent the minimum values that guarantee a sufficient number of synapses to match experimental data (Takács et al., 2012) and a certain room for the subsequent pruning step.

The second part of the algorithm discards synapses (pruning) in order to match experimental data on bouton densities (mean and standard deviation) and numbers of synapses per connection (mean). As experimental data were available for only a few pathways, we had to make additional assumptions for the uncharacterized pathways. According to Reimann et al. (2015), the number of appositions per connection can be used to predict the number of synapses per connection. We plotted the number of appositions as found by the model and the number of synapses for characterized pathways, and found that we can describe a good relationship among the two by splitting the data points into two groups (I-I connections and the rest) and fitting them separately (*y* = 0.1096*x* for I-I, *R* = 0.401, *p* = 0.325; *y* = 1.1690*x* for the rest, *R* = 0.267, *p* = 0.828) (Figure S11). For mean bouton density, we applied an average bouton density from characterized pathways to uncharacterized ones. Finally, the standard deviation of bouton density is estimated from the mean bouton density for the given pathway and the coefficient of variation (CV) estimated from a well-characterized pathway which we can generalize to all the other pathways. For somatosensory cortex, Reimann et al. (2015) used a CV of 0.32. For the hippocampus, the best source is Sik et al. (1995) which reported that the 64 PV postsynaptic cells receive 99 boutons from PVBC. They also reported that 51 neurons received one synapse, while 13 neurons received from two to four synapses, for a total of 99 - 51 = 48 synapses. So, the 13 neurons have on average 48/13 = 3.69 synapses per connection. We estimated the standard deviation to be 1.08 by considering 51 neurons with one synapse, and 13 neurons with 3.69 synapses. The resulting coefficient of variation is then 1.08/1.55 = 0.70. We scanned several CVs between these two extremes and checked how well the resulting connectomes matched experimental data on several parameters. We obtained the best results with a CV of 0.50, and we applied it to all the pathways.

As reported in Reimann et al. (2015) a bouton may form synapses onto two different postsynaptic neurons and we need to take this into account to better use data on bouton density to constrain the connectome. We replaced the value of 1.2 for synapses per bouton used by Reimann et al. (2015) with 1.15 taking into account newer unpublished data.

#### 4.17.1 Local synapse physiology

The methodology for the design of the synapse models, the extraction of the parameters, and their implementation is described in detail in Ecker et al. (2020) with the only exception of increasing the sample size of the number of pairs (i.e., 10,000 pre-post neurons) to increase the robustness of the calibration (see below). In brief, synapses were modeled with a stochastic version of the Tsodyks-Markram model (Tsodyks & Markram, 1997; Markram et al., 1998; Fuhrmann et al., 2002), featuring multivesicular release (Barros-Zulaica et al., 2019). We used sparse data from the literature (Tables S13 and S14) to parameterize the model in a pathway-specific manner. We aimed at characterizing the physiological properties of synapses like PSC rise and decay time constants, receptor ratios (NMDA/AMPA), and short-term plasticity (STP) profiles, which can be directly input into the model after some corrections (e.g., for calcium concentration, temperature, and liquid junction potential). We set the synaptic reversal potential (*E_rev_*) to 0 mV for AMPA and NMDA receptors, and to *−*80 mV for *GABA_A_*receptors; *τ_rise_*= 0.2 ms for AMPA and *GABA_A_* and 2.93 ms for NMDA receptors (Maccaferri et al., 2000; Andrásfalvy & Magee, 2001; Neu et al., 2007; Elfant et al., 2008; Fuentealba et al., 2008; Földy et al., 2010; Lee et al., 2010; Moradi & Ascoli, 2020).

We sampled 10,000 connected pairs from the circuit and replicated paired recordings *in silico* to calibrate N*_RRP_* (the number of vesicles in the release-ready pool) and peak synaptic conductance to match *in vitro* PSC CVs and PSP amplitudes, respectively (Barros-Zulaica et al., 2019; Ecker et al., 2020). Because of the sparsity of experimental data, at the end of this exercise, we only had six and 14 pathways (out of 130) with calibrated N*_RRP_* s and synaptic peak conductances, respectively. For the uncharacterized pathways, we had to generalize the values found.

As in a previous study, the release probability of the synapses scales nonlinearly with the extracellular calcium concentration, enriching their dynamical regime even further (Markram et al., 2015). The scaling of the release probability is made using a Hill function with three possible coefficients (steep, intermediate, shallow). NMDA/AMPA receptor ratios are pathway-dependent. Physiological evidence about release probability scaling and NMDA/AMPA ratios exists only for a minimal number of pathways. Thus, as the last step, we grouped the characterized pathways into 22 classes based on neurochemical markers, STP profiles, and peak synaptic conductances and used class average values predicatively for the remaining uncharacterized pathways.

### 4.18 Schaffer collaterals (SC)

#### 4.18.1 SC synapse anatomy

In the case of Schaffer collaterals, we did not model presynaptic CA3 pyramidal cells, and their axons, but directly the synapses on the postsynaptic neurons. For this reason, we followed an approach different from the one used for CA1 internal synapses, and we used the tool Projectionizer (Table 1), already adopted to model thalamo-cortical and cortical-thalamic projections (Markram et al., 2015; Reimann et al., 2022; Iavarone et al., 2023). In brief, the generation of projections was a multi-step workflow as described below.

The number of CA3 PCs was constrained considering the physiological ratios between CA3 PCs and CA1 PCs, as reported by Bezaire and Soltesz (2013), an approach consistent with the rest of the cell composition. Having estimated the number of presynaptic neurons (i.e., 267,238), we determined the number of afferent synapses on CA1 PCs to be 20,878 (the average of the range reported in Table 22 of Bezaire and Soltesz (2013), and the number of afferent synapses on interneurons to be 12,714 (the average of the range reported in Table 26 of Bezaire and Soltesz (2013). Details on the experimental data used to constrain SC anatomy are reported in Table S15.

To connect CA3 PC axons (i.e., SCs) with CA1 neurons, for each region in CA1, we gathered all the dendrite segment samples in the voxelized atlas. Neuron somas were not considered viable targets. Each region-wise candidate segment pool was then subsampled. Each region was assigned a number of synapses based on the synapse distribution (i.e., SLM: 0.3%, SR: 67.9%, SP: 7.1%, SO: 24.7%) and the total afferent synapses. The drawing from the pool was done with replacement, and sampling was weighted by the segment length not to oversample short segments. The sampled segments were considered as candidates for placing synapses. For each sampled segment, a random synapse position was chosen along the segment. Then, the candidate synapses were randomly assigned to each of the CA3 PCs presynaptic neurons. Finally, synaptic physiology parameters were drawn from distributions with specific means and standard deviations (see SC physiology Section). To match the average number of SC on PC and INT, the workflow was run separately for SC to PC and SC to INT.

#### 4.18.1 SC physiology

To define the synaptic parameters, we distinguish two pathways: SC-PC and SC-INT. In both cases, we used the Tsodyks-Markram model parameters estimated by Wierenga and Wadman (2003) (Tables S17 and S18), since we did not have experimental traces to estimate the parameters (as done with the internal synapses). We also set the NMDA/AMPA ratio, rise and decay time constant for NMDAR according to available data (Tables S17 and S18). The remaining parameters were not available, and we optimized them using two-steps procedure that is schematically reported in Figure S17B. In the first step, we set placeholder values for rise and decay time constant for AMPAR, and optimized maximum synaptic conductance and N*_RRP_*, while in the second step, we optimized the AMPAR time constants. In each step, we aimed to match a set of experimental measures that depend on the considered pathway. We used a grid search and selected the model parameters that minimized a cost function defined as:

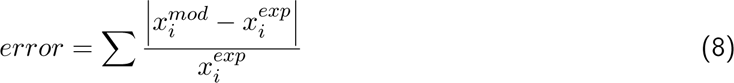

Where *x^exp^* is the experimental value we want to match, and *x^mod^* is the value produced by the model. Because of the strong parameter interdependence, after each step, the other step was rerun in order to verify if both held true. The cycle ended when the two steps converged toward a satisfactory solution (i.e. model and experiments are not statistically different). As done for the definition of the internal CA1 synaptic parameters, we ran all the steps selecting 10,000 random pairs of pre- and postsynaptic neurons.

##### Definition of SC-PC synaptic parameters

To constrain the peak synaptic conductance, N*_RRP_*, rise and decay time of AMPA and NMDA receptors, we aimed to match mean, CV, rise and decay time constant, and half-width of the PSP as reported in Sayer et al. (1990) (Table S17). We set up the simulations to best approximate the same experimental conditions (2.0 mM *Mg*^2+^ and 2.0 mM *Ca*^2+^).

##### Definition of SC-INT synaptic parameters

To constrain the peak synaptic conductance and N*_RRP_*, we used Glickfeld and Scanziani (2006) as a reference. The authors distinguished between cannabinoid receptor type 1 negative (CB1R-) basket cell (BC) and CB1R+ BC. Each cell type has a very distinctive response to SC stimulation, measured as the ratio between EPSC from SC to BC and the EPSC from SC to PC (*EPSC_BC_/EPSC_P C_*) (Table S18). We considered the CB1R- and CB1R+ BC to be representative respectively of all the CB1R- and CB1R+ INTs. We considered CB1R+ INTs to include PPA, CCKBC, and SCA, while CB1R-INTs the rest, according to the molecular marker profiles reported on www.hippocampome.org (Wheeler et al., 2015; Sanchez-Aguilera et al., 2021).

Following this strategy, we treated the two populations, CB1R-and CB1R+ INTs, separately, and we optimized the peak synaptic conductance and N*_RRP_* to match the corresponding EPSC ratio. We set up the simulations to best reproduce the experimental conditions (i.e., 1.3 mM *Mg*^2+^, 2.5 mM *Ca*^2+^, and NMDAR blocked). For each combination of parameters, we simulated 1,000 voltage-clamp experiments (neurons clamped at *−*85 mV as in the experiments) where we stimulated with a single spike one SC connected to one PC and one interneuron. For each of the 1,000 triplets SC-PC-interneuron, we have simulated 100 trials to have a sufficient number of traces to compute robust statistics of EPSCs. PSC ratios were computed only when there was a PSC in both PC and INT of the triplet. For each triplet, the mean PSC values were computed.

To optimize rise and decay time constants of AMPAR, we aimed to match the EPSP-IPSP latency as reported by Pouille and Scanziani (2001) (Table S18). We set up the simulations to reproduce the same experimental conditions (1.3 mM *Mg*^2+^, 2.5 mM *Ca*^2+^, and NMDAR blocked, with and without inhibition). For the sake of simplicity, we optimized the parameters of PVBC given their important role in feedforward inhibition (Pouille & Scanziani, 2001; Glickfeld & Scanziani, 2006) and then generalized the resulting parameters to the other INTs. To identify a feedforward loop SC-PV-PVBC, we proceeded as follows: 1) randomly select one PVBC; 2) select one PC connected to the PVBC; 3) select 200 SC fibers that innervate the PVBC. A simultaneous stimulation of 200 SC fibers is necessary to trigger an AP in PVBC; and 4) check whether at least one of these SC fibers also innervates the PC. If this is not the case, we repeat the procedure. We selected 1,000 SC-PC-PVBC triplets. For each triplet, we simulated 35 trials (different seeds). Each trial was 900 ms long, with spikes simultaneously delivered to the 200 SC fibers at 800 ms. We then made an average of the included voltage traces for each triplet and computed the EPSP with and without the inhibition. We derived the IPSP trace by subtracting the trace with inhibition from the trace without inhibition (Figure S17E1). Traces with EPSP or IPSP failures, without PC or INT spikes, or more than one spike were excluded from the analysis. Since some parameter combinations led to few valuable traces and this could bias the result, we included the number of surviving traces in our cost function (Equation 8), where *x^mod^* becomes the number of usable traces and *x^exp^* the total number of traces (i.e. 35,000).

#### 4.18.1 SC validation

The reconstruction of the SC were validated against the results of Sasaki et al. (2006) on input-output (I-O) characteristics of SC projections *in vitro*. We set up the simulations to mimic the same experimental conditions (slice of 300 µm, *Ca*^2+^ 2.4 m M, *Mg*^2+^ 2.4 m M, 32 °C) (Figure 4A). As in the experiment, we randomly selected 101 SP neurons in the slice. We randomly chose 350 SC inputs to activate simultaneously, evoking an AP at t = 1000 ms. This input was able to make all the 101 neurons spike corresponding to 100% of input/output (Figure 4B). We activated a different percentage of the 350 SC fibers, ranging from 5% to 100%, and quantified the number of spiking neurons. We repeated the protocol by blocking the GABAergic synapses, and repeated each condition (with and without GABAR) in five different slices. We computed the Pearson correlation coefficient R in control conditions to assess the linearity of IO curve in control condition (with GABAR).

### 4.19 Cholinergic modulation

To build a model of the effects of cholinergic release we began by collecting and curating literature findings from bath application experiments in which ACh, carbachol (CCh), or muscarine were used. We extracted data on their effects on neuron excitability (membrane potential, firing rate, Table S19), synaptic transmission (PSP, PSC, Table S20) and network activity (extracellular recordings, Table S21). We subsequently verified that the other experimental conditions (such as cell type, connection type and mouse vs rat provenance) did not produce further data stratification.

Since for each experiment, data was available for both control and drug-applied conditions, we estimated the relative amount of depolarizing current at different concentrations of ACh. For sub-threshold data, we computed the cell-type specific amount of current causing a change in the resting membrane potential equal to the mean value reported in the experiments. For supra-threshold data, we computed the amount of current needed to increase the firing rate from the baseline frequency to the frequency in ACh conditions.

The data points show a dose-dependent increase in depolarizing current. To describe this effect, we fitted a Hill function assuming the ACh-induced current at control condition is zero (*R*^2^ = 0.691, *N* = 28).

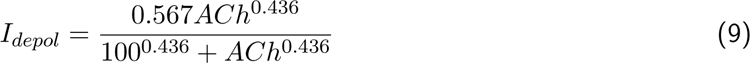

where *I_depol_*is the depolarizing current (in nA) and ACh is the neuromodulator concentration in µM (Figure 5A,C).

At the synaptic level, the data points show a dose-dependent decrease in the amplitude of the voltage/current response to pre-synaptic stimulation for all pathways analyzed (Table S20). Since ACh affects synaptic transmission principally at the level of release probability (Tsodyks & Markram, 1997; M. E. Hasselmo, 2006; D. Yang et al., 2021), we used the Tsodyks-Markram model (Tsodyks & Markram, 1997; Markram et al., 1998; Fuhrmann et al., 2002) and introduced scaling factor to make the parameter *U_SE_* (neurotransmitter release probability) dependent on ACh concentration with a Hill function fitted to experimental data (*R*^2^ = 0.667, *N* = 27).

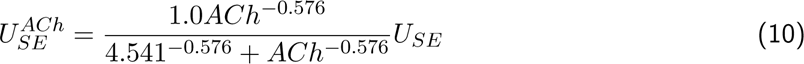

where *U_SE_*is the release probability (without ACh), and 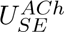 is the release probability with the dependency on ACh concentration (Figure 5B,D).

As a control, we verified that the values of PSP or PSC are proportional to *U_SE_*.

For each network simulation, we applied these equations to compute the effect of ACh concentration on cells and synapses. In particular, we inject the same amount of depolarizing current (equation 9) to all the cells and apply the same *U_SE_* scaling factor (Equation 10) to all pathways including Schaffer collaterals. All the simulations shown in figure 5E-K were run on the cylinder microcircuit with the extracellular calcium concentration set to 2 mM and a duration of 10 s. To disregard the initial ramping activity, we used the last 9 s of the simulation to compute correlation metrics.

### 4.20 Simulation

#### 4.20.1 Model instantiation and execution

For running the hippocampus simulations, the NEURON simulator is used and leverages the CoreNEU-RON optimized solver for improved efficiency (Awile et al., 2022). The model is instantiated using existing scripting methods for NEURON to construct the components of the virtual tissue and add support constructs such as stimuli and reports. Users can use the introspection feature of NEURON to adapt parameters based on certain features and identifiers. Once all components have their final values, the data structures are serialized to disk so that they can be loaded by the CoreNEURON solver into optimized data structures. This results in a 4-7*×* reduction in memory utilization by CoreNEURON, allowing larger simulation on the same hardware. The memory layout can better utilize hardware features such as vectorized instructions and can have a 2-7*×* improvement to execution time.

### 4.21 Simulation experiments

For simulations in Section 2.2, spike time, intracellular membrane potential, and extracellular voltage were recorded and analyzed.

#### 4.21.1 CA1 spontaneous synaptic release

The aim of these simulations was to discover whether spontaneous synaptic release in the intrinsic circuit would be sufficient to induce rhythmic activity in CA1. So for these simulations, the stochastic synaptic release was enabled for each model intrinsic synapse and the SC input disconnected for the entire period simulated, typically 10 s. *In vitro* CA1 spontaneous synaptic release rate mPSPs are generally estimated from recording postsynaptic events (Table S22), which is the summation of presynaptic release from the many converging axons. To determine what presynaptic rate parameter value to assign to the synapse models, we ran single neuron simulations for a range of mean presynaptic release rates. We verified that the presynaptic rate range produced a postsynaptic rate range that included ranges reported in literature (Figure S18A). We ran circuit simulations using this presynaptic rate range with both *in vitro*-and *in vivo*-like calcium concentrations. All other parameters remained constant across simulations.

#### 4.21.1 CA1 random synaptic barrage

The aim of these simulations was to discover whether random synaptic activity of the intrinsic circuit would induce oscillatory activity in CA1. Therefore, in these simulations, SC input was connected while stochastic synaptic release for intrinsic and extrinsic synapses was disabled for the entire period simulated, typically 10 s. SC spike times were generated independently for each input implemented using a Poisson random process with a constant (homogeneous) mean firing rate (Hz).To match the input firing rate of SC input that was able to generate regular theta activity in their CA1 model, we downloaded and analyzed example afferent input data provided with their Figure 5 of Bezaire et al. (2016). While Bezaire et al. (2016) quoted 0.65 Hz as the critical input rate of synaptic barrage required to generate theta activity, we found their actual spike data had a much lower rate *∼*0.14 Hz (*∼*64k spikes, the summation of their “proximal” and “distal” afferent sources arrived per second from *∼*450k afferents). To cover this range, we ran simulations with constant random spiking rates of 0.05-0.60 Hz and repeated these simulations at *in vitro*-and *in vivo*-like calcium levels. All other parameters remained constant across simulations.

#### 4.21.1 CA1 bath concentration of extracellular calcium and potassium ions

The aim of these simulations was to discover whether tonic depolarization of the intrinsic circuit led to the emergence of oscillatory activity in CA1. Hence, stochastic synaptic release was disabled for each model intrinsic synapse and SC input disconnected for the entire period simulated, typically 10 s. We modeled the bath effect of calcium as explained previously and the effect of extracellular potassium ion (*K*^+^) concentration as an constant current injected into the somatic compartment of each neuron (115-140% relative to each neuron’s rheobase current) (Markram et al., 2015). Simulations for this range of injected currents were repeated for a range of calcium concentrations between *in vitro*-and *in vivo*-like levels. All other parameters remained constant across simulations.

#### 4.21.1 CA3 theta oscillatory input

The aim of these simulations was to discover whether regular theta activity delivered via SC input to the intrinsic circuit would induce theta activity in CA1. So for these simulations, SC input was connected while stochastic synaptic release for intrinsic and extrinsic synapses was disabled for the entire period simulated, typically 10 s. SC spike times were generated independently for a subset of input axons (typically 15,000) using a Poisson random process with a sinusoidal (inhomogeneous) firing rate (Hz) using the Elephant software package functions (Denker et al., 2018). Here, the individual mean rate of firing (cell frequency, 0.1-0.4 Hz) defines the offset and amplitude that was sinusoidally modulated over time to represent theta-band (4-10 Hz) oscillation (signal frequency). Simulations spanning the range of combinations of cell frequency and signal frequency were repeated at *in vitro*-and *in vivo*-like calcium levels. All other parameters remained constant across simulations.

#### 4.21.1 Medial septum input

In this set of simulations, we aimed to study how MS input can induce theta oscillations in CA1. All the simulations run on the cylinder microcircuit with the extracellular calcium concentration set to 1 mM (*in vivo*-like level). All the neurons received a background depolarization, expressed as a percentage of the voltage threshold. Since the background depolarization is not completely known during theta oscillation, we tested values from 100% (threshold) to 130%. Above 130%, we observed non-physiological behaviors of the cells. The sinusoidal hyperpolarizing current injected in PV+ interneurons can be described by three parameters: frequency, mean, and amplitude. To reduce the parameter space to be scanned, we considered only the frequency of 8 Hz, an intermediate value in the theta range (4-12 Hz). In addition, we set *mean = -amplitude*. The disinhibition amplitude is unknown and we tested values in a range that produces physiological hyperpolarization in the PV+ interneuron models (0.1-0.5 nA). We scanned also different acetycholine concentrations which are biological plausible (0-2 *µ*M). The simulations ran for 20 s and disinhibition started at time 10 s.

#### 4.21.1 Propagation of oscillatory inputs

To examine whether CA3 gamma oscillations propagated to CA1, we aimed to reproduce the *in vitro* slice experimental results in Zemankovics et al. (2013). During these CCh-induced gamma oscillations the time course of CA3 spiking activity did not follow a pure sinusoidal waveform, so we had to create a custom inhomogeneous rate function. We did this by first manually reconstructing the probability of discharge of CA3 pyramidal cells over time (shown in Figure 4D, orange trace) and then mapped a smoothed version of these values from phase (radians) to time (seconds) coordinates to create a temporal rate function modulated at the reported gamma frequency of 31 Hz. The spike trains were generated from this custom rate function using the Elephant software package (Denker et al., 2018). Since the circuit did not model the topography of SC connections to CA1 neurons, it did not constrain how many active SCs project to the simulated slice. Therefore, we ran multiple simulations varying the number of activated SC axons with overall mean rate matching the reported average firing rate of CA3 PCs (Zemankovics et al., 2013). To mimic the same experimental conditions, we simulated a slice circuit with extracellular *Ca*^2+^ 2 m M and *Mg*^2+^ 2 m M. The effect of bath application of 10 *µ*M CCh effect on CA1 neurons was modeled using the approach described earlier (see Cholinergic modulation Section). These simulations ran for 5 s, during which stochastic synaptic release for intrinsic and extrinsic synapses was disabled, with SC input applied 2 s to 5 s. All other parameters remained constant across simulations.

In a final set of simulations, we studied how the CA1 circuit responds to CA3 oscillatory input covering a broad range of frequencies (0.5 - 200 Hz). We simulated an oscillatory input at CA3, with four different signal strengths (0.1, 0.2, 0.4, and 0.8 Hz, as mean firing rate) and spike trains were generated using Elephant (Denker et al., 2018). We measured the spiking response of CA1, checking if it was constant regardless of the oscillation frequency of CA3. Then, we computed the correlation between CA3 and CA1 and within CA1 neurons using the metrics explained below in Section 4.22.2, with 10,000 randomly selected cell pairs (CA3-CA1 or CA1-CA1, respectively). All spike train correlation measurements were repeated using standard covariance and cross-correlation functions. All the simulations run on the cylinder microcircuit with the extracellular calcium concentration set to 2 mM and a duration of 6 seconds. We used 1 s of activity (i.e., from 3 s to 4 s of simulation time) to compute gains and correlation metrics.

### 4.22 Analysis

Simulations showed an initial transient due to variable initiation and it appeared as an initial high activity of the network. This transient lasted for few ms, but we normally discarded the first 1000 ms in all the simulation analyses to be sure that the parameters converged into a stable regime.

#### 4.22.1 LFP analysis

For all simulations where extracellular voltage was recorded, the LFP analysis was performed in the same way unless otherwise stated. The raw extracellular voltage signal was estimated from multiple locations (channels) in the circuit model to mimic experimental electrode positions of a linear probe (Figure 2C). Following a standard electrophysiological processing protocol (e.g. Bokil et al., 2010), the raw extracellular signal was divided into two components using 6th-order Butterworth filtering implemented in the Elephant package (doi:10.5281/zenodo.1186602; RRID:SCR_003833) (Denker et al., 2018): a low-pass filtered signal (< 400 Hz cutoff) referred to as the local field potential (LFP) and a high-pass filtered signal (> 400 Hz). Here we analyzed the low-pass filtered LFP signal only. In the case of simulations reproducing the results of Zemankovics et al. (2013), LFP was band-pass filtered between 10-45 Hz to match their analysis protocol focusing on CA1 low gamma-band oscillations. Typically the first second of recording was discarded to eliminate onset transient artifacts. The remaining signal was detrended and tested for stationarity before being analyzed further (see Statistical analysis Section). For time-frequency analysis, this detrended signal was then downsampled from 2000 (1000/0.5 ms) to 400 Hz. The downsampled LFP signal was next filtered into separate frequency bands: delta (1-3 Hz), theta (4-12 Hz), and gamma (30-120 Hz). A multitaper method was used to estimate power spectral density (PSD) of the downsampled LFP signal with a frequency resolution of 1.5 Hz (Prieto et al., 2009, mtspec package: https://pypi.org/project/mtspec/). To estimate the spatiotemporal LFP spectral response, a complex Morlet wavelet transform (CWT) was applied with n_cycles = 7 for 1-150 Hz range in steps of 0.25 Hz using the Elephant package function and reported in decibel (dB) units (Addison et al., 2009). To identify cross-laminar source-sink relationships, current source density (CSD) analysis was applied to the theta-band filtered signal across all channels of the virtual electrode using the Elephant package function for KCSD1D method (Potworowski et al., 2012).

For simulations that generated regular theta oscillations, spike-LFP phase coupling and theta waveform analyses were performed. To quantify phase preference of neurons during stable periods of theta oscillations (defined as at least three 2 second periods of theta in Klausberger et al., 2003), the Hilbert analytic transform of single channel theta-band signal (unless otherwise stated in SP) was calculated to estimate the instantaneous phase using the Elephant package function. The instantaneous phase was used to assign a (theta) phase angle to individual spike times for a period over two consecutive theta cycles (0-720 degree period) with trough of theta cycle set as 0 degrees (Klausberger et al., 2003). For each neuron, the resultant (summed) vector angle for its spike train was used to determine the preferred phase of firing and the vector norm estimated the degree of phase locking (see Statistical analysis Section). To quantify waveform asymmetry, we measured the asymmetry index for a low-pass LFP signal filtered in 1-80 Hz range (Belluscio et al., 2012). To identify peak and trough locations, we used z-score thresholding, as the Hilbert method yielded too many spurious locations, and from this we computed the duration of rise (*t_rise_*) and decay (*t_decay_*) for each oscillation. The asymmetry index per oscillation was calculated from log(*t_rise_/t_decay_*).

#### 4.22.1 Correlation

To have a quantitative measurement of correlations between spike trains, we mainly used the spike time tiling coefficient (STTC) and the mean spike-spike correlations, computed for a defined number of cell pairs (normally 10,000, for robustness) as the histogram of intervals between all spike times of two different cells. For both STTC and spike-spike correlations we used a bin size of 10 ms, considering 1 s of simulated activity.

To verify the results obtained with the STTC method, we computed with the same input data the correlation using standard covariance and cross-correlation functions. All correlation analyses have been done with the Elephant library (Denker et al., 2018).

### 4.23 Statistical analysis

To calculate the slopes in linear fits to the appositions data, we used least-squares solution from the python toolbox (numpy.linalg.lstsq function).

For the fitting of experimental data points of cholinergic modulation we used the non-linear least squares solution (scipy.optimize.curve_fit function). Values are expressed as *R*^2^ coefficients.

For LFP analysis, the Augmented Dickey-Full (ADF) unit root test was used to determine whether a single channel continuous signal was stationary before analyzing its spectral properties (statsmodels.tsa. stattools.adfuller function). To determine whether single neurons were phase-locked, the Rayleigh Test of randomness of circular data was used to reject the null hypothesis that phase angles of a spike train were uniformly distributed (Wilkie, 1983). To compare with empirical results (e.g., Klausberger et al., 2003), phase analysis results were described by mean *±* angular deviation in degrees.

### 4.24 Statistical comparison methodology

Our statistical comparisons between the model and experimental data were guided by the experimental data availability. To streamline the process, we performed statistical comparisons at an aggregate level rather than conducting comparisons for each individual cell type or pathway.

There are two main reasons to perform statistical comparisons at an aggregate level. Firstly, there may be a lack of data available at a granular level, such as for cell composition validation. Secondly, for certain parameters, such as bouton density, because we used a multi-objective optimization approach to match experimental data at an aggregate level, it is more appropriate to validate at an aggregate level as well. For these comparisons, we typically use Pearson’s Correlation Coefficient (scipy.stats.pearsonr). Values are expressed as (Pearson correlation coefficient, *p*-value).

In cases where we needed to compare a large number of features for a metric, such as in morphology cloning validation, we used similarity scores in addition to correlation analysis. However, in other cases, when only a mean and standard deviation value were available for comparison, as in Schaffer collateral comparison, we used a z-test or t-test. Values are expressed as (*p*-value).

Finally, if the experimental data had a very small sample size, such as in the case of population synchrony, we did not perform any statistical and we relied only on qualitative assessments.

### 4.25 Visualization

#### 4.25.1 Brayns

Hippocampus circuit and simulations were visualized using Brayns software and its internally developed web interfaces: WebBrayns and Brayns Circuit Studio (Table 1).

Brayns is a visualization software based on ray tracing techniques. It allows rendering 3D scenes and producing high quality images and videos. Brayns offers programming interfaces in C++ and Python that make it highly customizable, while its web-based interfaces allow the interactive exploration of the scene and make the platform accessible to a wider community (no programming skills needed). In order to create high-quality, high-resolution visuals, Brayns makes use of different rendering engines, like Intel OSPRay (CPU-based ray tracing library, https://www.ospray.org/) or NVIDIA Optix (GPU-based ray-tracing framework https://developer.nvidia.com/rtx/ray-tracing/optix).

The capability of Brayns to render large-scale models, like the whole hippocampus circuit, is key to supporting scientific visualization needs and providing insight on scientific aspects that otherwise are very complex to analyze (e.g. signal propagation at circuit level).

The morphology collage images were also produced with Brayns. A set of clipping planes were used to determine the locations of the slices. For each plane, a second, parallel plane was placed 100 µm further from the origin to create a slice of 100 µm thickness. Then, for each slice and for each neuron morphological type, a small set of neurons of the chosen morphological type and physically located inside the volume determined by the slice were picked and rendered with Brayns together with the polygonal meshes that define the outline of CA1 layers. The set of neurons displayed in every slice was chosen in a way to maximize its physical distribution within the slice (Figure S9A).

Some of the morphology collage images were post-processed with media design tools (e.g., Adobe Illustrator, Adobe Photoshop) to match the visual style of other figures.

#### 4.25.1 NeuroMorphoVis

NeuroMorphoVis (NMV, see Table 1) was used to visualize single morphologies (Abdellah et al., 2018). NMV is a Blender plug-in that allows the visualization and analysis of neuronal morphology skeletons that are digitally reconstructed. NMV presents many features, including the manual repair of broken morphology skeletons and the creation of accurate meshes that represent the membranes of the morphologies.

### 4.26 List of assumptions

#### 4.26.1 General

- This list is not exhaustive. We describe the assumptions that are specific to this work and the ones that could be revised in subsequent refinements
- We do not list here the assumptions that are already included in and derive from the adoption of specific other models (e.g., Markram et al. (2015); animal models)
- Certain assumptions involve the exclusion of specific features. For example, gap junctions, rare cell types, glial cells, vasculature, are excluded from the current datasets and model.
- In addition to the preceding point, certain limitations were imposed by both data quality and accessibility.

4.26.1 Data

- Most of the data came from the dorsal CA1. We approximated the entire CA1 using data from dorsal CA1.
- We mixed data from different labs, experimental conditions, and animal models. We assumed that the inter-datasets variability is less than the intra-individual variability.

4.26.1 Volume

- We approximated the layer anatomy by dividing the overall CA1 volume into parallel layers with a fixed ratio between their thicknesses. The ratio was taken from the analyses of slice reconstructions from dorsal CA1 of adult rats. We presumed that any potential error introduced by this approximation was comparatively smaller than the inherent noise present in the original atlas.

#### 4.26.1 Morphologies

- We considered the reconstructed cell morphologies to be fairly complete.
- One PC reconstruction showed an axon with a length of 3.646 µm and an extension of 1.325 µm along the transverse axis. We deemed this axon to be relatively complete within the CA1 and therefore used it as a prototype for all the PCs. We presumed that using other axons would introduce a larger problem into connectivity with relatively little gain in diversity.
- We treated PCs as one homogeneous cell type.
- SP_PVBC and SP_CCKBC were classified as two separate classes.
- We defined PV+ cells as SP_PVBC, SP_BS, SP_AA, CCK+ cells as SP_CCKBC, SR_SCA, SLM_PPA and SOM+ (somatostatin) cells as SO_OLM, SO_BS, SO_BP, SO_Tri.
- Scaling and cloning compensated for the small sample size (Markram et al., 2015).

#### 4.26.1 Electrical types

- We assumed only four e-types: classical accommodating (cAC), bursting accommodating (bAC) and classical non-accommodating (cNAC) for interneurons, and classical accommodating for pyramidal cells (cACpyr).

#### 4.26.1 Cell composition

- For a given cell type, we assumed that all the expected cells in CA1 are located in the layers for which we had the corresponding morphological reconstructions.
- Trilaminar cells and radiatum-retrohippocampal cells in SO had the same proportion.
- To compensate for the lack of some inhibitory types, we increased the interneurons to match the expected E/I ratio of 11:89 (Bezaire & Soltesz, 2013). In doing so, we assumed that matching E/I ratio was more important that maintaining the expected number of cell types present in the model.

#### 4.26.1 Cell positioning

- The cell somas could have been located in any point of the layers (random placement).
- Cells had a principal axis that was parallel to the radial axis and perpendicular to the layers.
- Lacking precise evidence, we allowed random rotations around the y-axis for the interneurons.

#### 4.26.1 Connectome

- Axo-axonic cells contacted only pyramidal cells.
- We did not allow SCA, Ivy, and BS to form synapses on PC somas to prevent the number of somatic synapses from far exceeding expected values. In doing so, we assumed that the exclusion of few synapses would result in a smaller error compared to the one generated by a large number of somatic synapses.

#### 4.26.1 Single cell models

- The considered electrical features were the ones that described most of the behavior of the cells (R. Migliore et al., 2018).
- We considered channels to be uniformly distributed in all dendritic compartments except KA and Ih.
- We applied the same currents of pyramidal cells to interneurons with exceptions supported by experimental evidence.

#### 4.26.1 Synapse model

- Synapses between A and B had the same parameters.
- Synapses between m-types A and B had parameters extracted from the same distributions (truncated Gaussian).
- Generalization: the synapse type and dynamics were defined, first of all, by pre- and post-synaptic m-type.
- The NMDA/AMPA ratio was kept constant through all compartments of a given neuron.

#### 4.26.1 Network

- The action potential was propagated stereotypically from AIS to synapse with a fixed velocity of 300 mm/s (Markram et al., 2015).
- In slices, neurons preserved their integrity. We did not model cut neurons.

#### 4.26.1 Schaffer collaterals (SC)

- SC synapses were uniformly distributed along the transverse and longitudinal axes.
- SC synapses could have been placed at all locations on the target neuron (except on the soma), including apical tuft dendrites.
- While Glickfeld and Scanziani (2006) measured EPSC only on basket cells, then divided into cannabinoid receptor type 1 negative (CB1R-) and positive (CB1R+), we applied the experimental measurements to all interneurons, dividing them into two categories according to their positivity/negativity to CB1 markers, as reported in https://hippocampome.org/php/markers.php.
- STP parameters (i.e., *U*, *D*, *F*, in the Tsodyks-Markram Model (Tsodyks & Markram, 1997)) were uniform for all interneurons, and they followed the values identified by Wierenga and Wadman (2003).

#### 4.26.1 Acetylcholine (ACh)

- The dose-response curves for cholinergic modulation was homogeneous for all cell and synapse types.
- The dose-response was well described using the Hill equation.
- ACh, CCh, and muscarine had the same effects on neuronal excitability and synaptic transmission.
- The change in membrane excitability caused by ACh was assumed to be equivalent to a tonic current injection at the soma.
- Perisynaptic effects of volumetric transmission (i.e., non-synaptic release) of ACh was neglected.

## Acknowledgments

The authors would like to thank all the people involved in the rat CA1 hippocampus project over the last years, in particular Attila Gulyás, Luc Guyot, and Arseny Povolotsky. We would also like to thank those who offered valuable advice or data: Giorgio Ascoli, Norbert Hájos, Jesse Jackson, Corette Wierenga, Sylvain Williams. Experimentalists who generated data for this project or who recorded, dye-filled and/or reconstructed neurons: A.B. Ali, A.P Bannister, R. Begum, N. Botcher, J. Deuchars, K. Eastlake, D. I. Hughes, M. Ilia, J. Kerkhoff, S. Kirchhecker, H. Pawelzik, and H. Trigg. D.C. West designed data collection and analysis software. We thank those who worked on the Explore feature of the Hippocampus hub: Anil Tuncel, Pavlo Getta, Caitlin Monney, Stefano Antonel, Alexander Dietz, and Liviu Soltuzu. Finally, we are indebted to Karin Holm for copy editing and publication advice and support.

## 5.1 Author contributions

H.M. conceived and led the study. S.K., M.M., A.T., F.S., E.M., and A.M. co-led the study. A.M.T. and A.M. planned, performed and supervised electrophysiological experiments and neuron reconstructions. J.F. and S.L. performed reconstructions. A.R., J.B., A.A., D.B., K.K. planned and supervised on data integration, strategies and algorithms, model building, simulation experiments, and analysis. A.A., J.B., D.B., A.R. reconstructed Schaffer collaterals. A.A., C.C., J.B., A.R. modeled Acetylcholine. J.B. and A.R. worked on theta. A.A., J.B. and A.R. worked on oscillation propagation. F.S. and J-D.C. planned and supervised the development of algorithms, software and workflows, computing infrastructure, and technical integration. A.R., J.B., A.A., D.B., C.C., K.K. wrote the manuscript. A detailed listing of author contributions is available in the Supplemental materials.

## 6 Software used

Table 1 gives a list of software used in the paper.

Table 1: List of software used
| Software Name | Source | Identifier |
| --- | --- | --- |
| atlas-direction-vectors | BBP/EPFL software package | https://github.com/BlueBrain/atlas-direction-vectors |
| bbp-workflow | BBP/EPFL software package | not yet open source |
| BluePy | BBP/EPFL software package | not yet open source |
| BluePySNAP | BBP/EPFL software package | https://github.com/BlueBrain/snap |
| BluePyEfe | BBP/EPFL software package | https://github.com/BlueBrain/BluePyEfe |
| BluePyOpt | BBP/EPFL software package | https://github.com/BlueBrain/BluePyOpt |
| BluePyMM | BBP/EPFL software package | https://github.com/BlueBrain/BluePyMM |
| Brainbuilder | BBP/EPFL software package | not yet open source |
| Brayns | BBP/EPFL software package | https://github.com/BlueBrain/Brayns |
| circuit-build | BBP/EPFL software package | not yet open source |
| CoreNEURON | BBP/EPFL software package | https://github.com/BlueBrain/CoreNeuron |
| eFEL | BBP/EPFL software package | https://github.com/BlueBrain/eFEL |
| Elephant | Elephant authors and contributors | https://doi.org/10.5281/zenodo.1186602 |
| EMSim | BBP/EPFL software package | https://github.com/BlueBrain/EMSim |
| Hippounit | KOKI software package | https://github.com/KaliLab/hippounit |
| ITK-SNAP | University of Pennsylvania | http://www.itksnap.org/ |
| morphology-workflows | BBP/EPFL software package | https://github.com/BlueBrain/morphology-workflows |
| mtspec | pypi python package | https://pypi.org/project/mtspec/ |
| neo | The NeuralEnsemble Initiative | https://github.com/NeuralEnsemble/python-neo |
| Neurodamus | BBP/EPFL software package | https://github.com/BlueBrain/neurodamus |
| NeuroMorphoVis | BBP/EPFL software package | https://github.com/BlueBrain/NeuroMorphoVis |
| NeuroM | BBP/EPFL software package | https://github.com/BlueBrain/NeuroM |
| NeuroR | BBP/EPFL software package | https://github.com/BlueBrain/NeuroR |
| projectionizer | BBP/EPFL software package | not yet open source |
| psp-validation | BBP/EPFL software package | not yet open source |
| regiodesics | BBP/EPFL software package | not yet open source |
| TMD | BBP/EPFL software package | https://github.com/BlueBrain/TMD |
| voxcell | BBP/EPFL software package | https://github.com/BlueBrain/voxcell |

## References

Jung, R., & Kornmüller, A. E. (1938). Eine methodik der ableitung iokalisierter potentialschwankungen aus subcorticalen hirngebieten. Archiv für Psychiatrie und Nervenkrankheiten, 109 (1), 1–30. 10.1007/BF02157817

Green, J. D., & Arduini, A. A. (1954). Hippocampal electrical activity in arousal. J. Neurophysiol., 17 (6), 533–557. 10.1152/jn.1954.17.6.533

Grastyan, E., Lissak, K., Madarasz, I., & Donhoffer, H. (1959). Hippocampal electrical activity during the development of conditioned reflexes. Electroencephalogr. Clin. Neurophysiol., 11 (3), 409–430. 10.1016/0013-4694(59)90040-9

Vanderwolf, C. H. (1969). Hippocampal electrical activity and voluntary movement in the rat. Electroencephalogr. Clin. Neurophysiol., 26 (4), 407–418. 10.1016/0013-4694(69)90092-3

O’Keefe, J., & Dostrovsky, J. (1971). The hippocampus as a spatial map. preliminary evidence from unit activity in the freely-moving rat. Brain Res., 34 (1), 171–175. 10.1016/0006-8993(71)90358-1

Knowles, W. D., & Schwartzkroin, P. A. (1981). Local circuit synaptic interactions in hippocampal brain slices [Publisher: Society for Neuroscience Section: Articles]. Journal of Neuroscience, 1 (3), 318–322. 10.1523/JNEUROSCI.01-03-00318.1981

Baimbridge, K. G., & Miller, J. J. (1982). Immunohistochemical localization of calcium-binding protein in the cerebellum, hippocampal formation and olfactory bulb of the rat. Brain Research, 245 (2), 223–229. 10.1016/0006-8993(82)90804-6

Morris, R. G., Garrud, P., Rawlins, J. N., & O’Keefe, J. (1982). Place navigation impaired in rats with hippocampal lesions. Nature, 297 (5868), 681–683. 10.1038/297681a0

Wilkie, D. (1983). Rayleigh test for randomness of circular data. J. R. Stat. Soc. Ser. C Appl. Stat., 32 (3), 311. 10.2307/2347954

Buzsáki, G., Rappelsberger, P., & Kellényi, L. (1985). Depth profiles of hippocampal rhythmic slow activity (‘theta rhythm’) depend on behaviour. Electroencephalogr. Clin. Neurophysiol., 61 (1), 77–88. 10.1016/0013-4694(85)91075-2

Traub, R. D., Miles, R., & Wong, R. S. (1988). Large scale simulations of the hippocampus. IEEE Eng. Med. Biol. Mag., 7 (4), 31–38. 10.1109/51.20378

Amaral, D. G., & Witter, M. P. (1989). The three-dimensional organization of the hippocampal formation: A review of anatomical data. Neuroscience, 31 (3), 571–591. 10.1016/0306-4522(89)90424-7

Sayer, R. J., Friedlander, M. J., & Redman, S. J. (1990). The time course and amplitude of EPSPs evoked at synapses between pairs of CA3/CA1 neurons in the hippocampal slice. The Journal of Neuroscience: The Official Journal of the Society for Neuroscience, 10 (3), 826–836.

Sheridan, R. D., & Sutor, B. (1990). Presynaptic m1 muscarinic cholinoceptors mediate inhibition of excitatory synaptic transmission in the hippocampus in vitro. Neuroscience Letters, 108 (3), 273–278. 10.1016/0304-3940(90)90653-Q

Williams, S., & Johnston, D. (1990). Muscarinic depression of synaptic transmission at the hippocampal mossy fiber synapse. Journal of Neurophysiology, 64 (4), 1089–1097. 10.1152/jn.1990.64.4.1089

Traub, R. D., & Miles, R. (1991). Neuronal networks of the hippocampus. Cambridge University Press. 10.1017/cbo9780511895401

Traub, R. D., Miles, R., & Buzsáki, G. (1992). Computer simulation of carbachol-driven rhythmic population oscillations in the CA3 region of the in vitro rat hippocampus. J. Physiol. 10.1113/jphysiol.1992.sp019184

Sik, A., Tamamaki, N., & Freund, T. F. (1993). Complete axon arborization of a single CA3 pyramidal cell in the rat hippocampus, and its relationship with postsynaptic parvalbumin-containing interneurons. European Journal of Neuroscience, 5 (12), 1719–1728. 10.1111/j.1460-9568.1993.tb00239.x

Aika, Y., Ren, J., Kosaka, K., & Kosaka, T. (1994). Quantitative analysis of GABA-like-immunoreactive and parvalbumin-containing neurons in the CA1 region of the rat hippocampus using a stereological method, the disector. Experimental Brain Research, 99 (2). 10.1007/BF00239593

Bianchi, R., & Wong, R. K. (1994). Carbachol-induced synchronized rhythmic bursts in CA3 neurons of guinea pig hippocampus in vitro. Journal of Neurophysiology, 72 (1), 131–138. 10.1152/jn.1994.72.1.131

Buhl, E. H., Han, Z. S., Lorinczi, Z., Stezhka, V. V., Karnup, S. V., & Somogyi, P. (1994). Physiological properties of anatomically identified axo-axonic cells in the rat hippocampus. Journal of Neurophysiology, 71 (4), 1289–1307. 10.1152/jn.1994.71.4.1289

Buhl, E. H., Halasy, K., & Somogyi, P. (1994). Diverse sources of hippocampal unitary inhibitory postsynaptic potentials and the number of synaptic release sites. Nature, 368 (6474), 823–828. 10.1038/368823a0

Hasselmo, M., & Schnell, E. (1994). Laminar selectivity of the cholinergic suppression of synaptic transmission in rat hippocampal region CA1: Computational modeling and brain slice physiology. The Journal of Neuroscience, 14 (6), 3898–3914. 10.1523/JNEUROSCI.14-06-03898.1994

Li, X.-G., Somogyi, P., Ylinen, A., & Buzsáki, G. (1994). The hippocampal CA3 network: An in vivo intracellular labeling study: THE HIPPOCAMPAL CA3 NETWORK. Journal of Comparative Neurology, 339 (2), 181–208. 10.1002/cne.903390204

Bannister, N. J., & Larkman, A. U. (1995). Dendritic morphology of CA1 pyramidal neurones from the rat hippocampus: I. branching patterns. The Journal of Comparative Neurology, 360 (1), 150–160. 10.1002/cne.903600111

Debanne, D., Guérineau, N. C., Gähwiler, B. H., & Thompson, S. M. (1995). Physiology and pharmacology of unitary synaptic connections between pairs of cells in areas CA3 and CA1 of rat hippocampal slice cultures. J. Neurophysiol., 73 (3), 1282–1294. 10.1152/jn.1995.73.3.1282

Sik, A., Penttonen, M., Ylinen, A., & Buzsáki, G. (1995). Hippocampal CA1 interneurons: An in vivo intracellular labeling study. The Journal of Neuroscience, 15 (10), 6651–6665. 10.1523/JNEUROSCI.15-10-06651.1995

Ylinen, A., Soltész, I., Bragin, A., Penttonen, M., Sik, A., & Buzsáki, G. (1995). Intracellular correlates of hippocampal theta rhythm in identified pyramidal cells, granule cells, and basket cells. Hippocampus, 5 (1), 78–90. 10.1002/hipo.450050110

Deuchars, J., & Thomson, A. (1996). CA1 pyramid-pyramid connections in rat hippocampus in vitro: Dual intracellular recordings with biocytin filling. Neuroscience, 74 (4), 1009–1018.10.1016/0306-4522(96)00251-5

Freund, T., & Buzsáki, G. (1996). Interneurons of the hippocampus. Hippocampus, 6 (4), 347–470. 10.1002/(SICI)1098-1063(1996)6:4<347::AID-HIPO1>3.0.CO;2-I

Kawasaki, H., & Avoli, M. (1996). Excitatory effects induced by carbachol on bursting neurons of the rat subiculum. Neuroscience Letters, 219 (1), 1–4. 10.1016/S0304-3940(96)13175-X

Hájos, N., & Mody, I. (1997). Synaptic communication among hippocampal interneurons: Properties of spontaneous IPSCs in morphologically identified cells. J. Neurosci., 17 (21), 8427–8442.

Hoffman, D. A., Magee, J. C., Colbert, C. M., & Johnston, D. (1997). K+ channel regulation of signal propagation in dendrites of hippocampal pyramidal neurons. Nature, 387 (6636), 869– 875. 10.1038/43119

Markram, H., Lübke, J., Frotscher, M., Roth, A., & Sakmann, B. (1997). Physiology and anatomy of synaptic connections between thick tufted pyramidal neurones in the developing rat neocortex. The Journal of Physiology, 500, 409–440. Retrieved September 21, 2022, from https://www.ncbi.nlm.nih.gov/pmc/articles/PMC1159394/

Tsodyks, M., & Markram, H. (1997). The neural code between neocortical pyramidal neurons depends on neurotransmitter release probability. Proceedings of the National Academy of Sciences, 94 (2), 719–723. 10.1073/pnas.94.2.719

Williams, J. H., & Kauer, J. A. (1997). Properties of carbachol-induced oscillatory activity in rat hippocampus. Journal of Neurophysiology, 78 (5), 2631–2640. 10.1152/jn.1997.78.5.2631

Ali, A. B., Deuchars, J., Pawelzik, H., & Thomson, A. M. (1998). CA1 pyramidal to basket and bistratified cell EPSPs: Dual intracellular recordings in rat hippocampal slices. J. Physiol., 507 (Pt 1), 201–217.

Csicsvari, J., Hirase, H., Czurko, A., & Buzsáki, G. (1998). Reliability and state dependence of pyramidal cell-interneuron synapses in the hippocampus: An ensemble approach in the behaving rat. Neuron, 21 (1), 179–189. 10.1016/s0896-6273(00)80525-5

Fisahn, A., Pike, F. G., Buhl, E. H., & Paulsen, O. (1998). Cholinergic induction of network oscillations at 40 hz in the hippocampus in vitro. Nature, 394 (6689), 186–189. 10.1038/28179

Markram, H., Wang, Y., & Tsodyks, M. (1998). Differential signaling via the same axon of neocortical pyramidal neurons. Proceedings of the National Academy of Sciences of the United States of America, 95 (9), 5323–5328. 10.1073/pnas.95.9.5323

Vida, I., Halasy, K., Szinyei, C., Somogyi, P., & Buhl, E. H. (1998). Unitary IPSPs evoked by interneurons at the stratum radiatum-stratum lacunosum-moleculare border in the CA1 area of the rat hippocampus *in vitro*. The Journal of Physiology, 506 (3), 755–773. 10.1111/j.1469-7793.1998.755bv.x

Ali, A. B., Bannister, A. P., & Thomson, A. M. (1999). IPSPs elicited in CA1 pyramidal cells by putative basket cells in slices of adult rat hippocampus: Basket cell IPSPs in CA1 pyramidal cells. European Journal of Neuroscience, 11 (5), 1741–1753. 10.1046/j.1460-9568.1999.00592.x

Csicsvari, J., Hirase, H., Czurkó, A., Mamiya, A., & Buzsáki, G. (1999). Oscillatory coupling of hippocampal pyramidal cells and interneurons in the behaving rat. J. Neurosci., 19 (1), 274–287. 10.1523/JNEUROSCI.19-01-00274.1999

Esclapez, M., Hirsch, J. C., Ben-Ari, Y., & Bernard, C. (1999a). Newly formed excitatory pathways provide a substrate for hyperexcitability in experimental temporal lobe epilepsy. J. Comp. Neurol., 408 (4), 449–460. 10.1002/(sici)1096-9861(19990614)408:4<449::aid-cne1>3.0.co;2-r

Esclapez, M., Hirsch, J. C., Ben-Ari, Y., & Bernard, C. (1999b). Newly formed excitatory pathways provide a substrate for hyperexcitability in experimental temporal lobe epilepsy. The Journal of Comparative Neurology, 408 (4), 449–460. 10.1002/(SICI)1096 - 9861(19990614)408:4<otherinfo><449::AID-CNE1>3.0.CO;2-R</otherinfo>

Hoffman, D. A., & Johnston, D. (1999). Neuromodulation of dendritic action potentials. J. Neurophysiol., 81 (1), 408–411. 10.1152/jn.1999.81.1.408

Magee, J. C. (1999). Dendritic lh normalizes temporal summation in hippocampal CA1 neurons. Nat. Neurosci., 2 (6), 508–514. 10.1038/9158

McQuiston, A. R., & Madison, D. V. (1999). Muscarinic receptor activity has multiple effects on the resting membrane potentials of CA1 hippocampal interneurons. The Journal of Neuroscience, 19 (14), 5693–5702. 10.1523/JNEUROSCI.19-14-05693.1999

Migliore, M., Hoffman, D. A., Magee, J. C., & Johnston, D. (1999). Role of an a-type k+ conductance in the back-propagation of action potentials in the dendrites of hippocampal pyramidal neurons. Journal of Computational Neuroscience, 7 (1), 5–15. 10.1023/a:1008906225285

Pawelzik, H., Bannister, A. P., Deuchars, J., Ilia, M., & Thomson, A. M. (1999). Modulation of bistratifed cell IPSPs and basket cell IPSPs by pentobarbitone sodium, diazepam and zn2+: Dual recordings in slices of adult rat hippocampus. European Journal of Neuroscience, 13.

Cossart, R., Hirsch, J. C., Cannon, R. C., Dinoncourt, C., Wheal, H. V., Ben-Ari, Y., Esclapez, M., & Bernard, C. (2000). Distribution of spontaneous currents along the somato-dendritic axis of rat hippocampal CA1 pyramidal neurons. Neuroscience, 99 (4), 593–603. 10.1016/s0306-4522(00)00231-1

Fellous, J. M., & Sejnowski, T. J. (2000). Cholinergic induction of oscillations in the hippocampal slice in the slow (0.5-2 hz), theta (5-12 hz), and gamma (35-70 hz) bands. Hippocampus, 10 (2), 187–197. 10.1002/(SICI)1098-1063(2000)10:2<187::AID-HIPO8>3.0.CO;2-M

Hughes, D. I., Bannister, A., Pawelzik, H., & Thomson, A. M. (2000). Double immunofluorescence, peroxidase labelling and ultrastructural analysis of interneurones following prolonged electrophysiological recordings in vitro. Journal of Neuroscience Methods, 101 (2), 107–116. 10.1016/S0165-0270(00)00254-5

Maccaferri, G., David, J., Roberts, B., Szucs, P., Cottingham, C. A., & Somogyi, P. (2000). Cell surface domain specific postsynaptic currents evoked by identified GABAergic neurones in rat hippocampus *in vitro*. The Journal of Physiology, 524 (1), 91–116. 10.1111/j.1469-7793.2000.t01-3-00091.x

Magee, J. C., & Cook, E. P. (2000). Somatic EPSP amplitude is independent of synapse location in hippocampal pyramidal neurons. Nat. Neurosci., 3 (9), 895–903. 10.1038/78800

Thomson, A. M., Bannister, A. P., Hughes, D. I., & Pawelzik, H. (2000). Differential sensitivity to zolpidem of IPSPs activated by morphologically identified CA1 interneurons in slices of rat hippocampus: Zolpidem enhancement of basket cell IPSPs in CA1. European Journal of Neuroscience, 12 (2), 425–436. 10.1046/j.1460-9568.2000.00915.x

Traub, R. D., Bibbig, A., Fisahn, A., LeBeau, F. E., Whittington, M. A., & Buhl, E. H. (2000). A model of gamma-frequency network oscillations induced in the rat CA3 region by carbachol in vitro. Eur. J. Neurosci., 12 (11), 4093–4106. 10.1046/j.1460-9568.2000.00300.x

Andrásfalvy, B. K., & Magee, J. C. (2001). Distance-dependent increase in AMPA receptor number in the dendrites of adult hippocampal CA1 pyramidal neurons. The Journal of Neuroscience, 21 (23), 9151–9159. 10.1523/JNEUROSCI.21-23-09151.2001

Golding, N. L., Kath, W. L., & Spruston, N. (2001). Dichotomy of action-potential backpropagation in CA1 pyramidal neuron dendrites. J. Neurophysiol., 86 (6), 2998–3010. 10.1152/jn.2001.86.6.2998

Harris, E., & Stewart, M. (2001). Propagation of synchronous epileptiform events from subiculum backward into area CA1 of rat brain slices. Brain Res., 895 (1-2), 41–49. 10.1016/s0006-8993(01)02023-6

Megias, M., Emri, Z., Freund, T., & Gulyas, A. (2001). Total number and distribution of inhibitory and excitatory synapses on hippocampal CA1 pyramidal cells. Neuroscience, 102 (3), 527–540. 10.1016/S0306-4522(00)00496-6

Pouille, F., & Scanziani, M. (2001). Enforcement of temporal fidelity in pyramidal cells by somatic feed-forward inhibition. Science (New York, N.Y.), 293 (5532), 1159–1163. 10.1126/science.1060342

Buzsáki, G. (2002). Theta oscillations in the hippocampus. Neuron, 33 (3), 325–340. 10.1016/s0896-6273(02)00586-x

Fuhrmann, G., Segev, I., Markram, H., & Tsodyks, M. (2002). Coding of temporal information by activity-dependent synapses. Journal of Neurophysiology, 87 (1), 140–148. 10.1152/jn.00258.2001

Pawelzik, H., Hughes, D. I., & Thomson, A. M. (2002). Physiological and morphological diversity of immunocytochemically defined parvalbumin-and cholecystokinin-positive interneurones in CA1 of the adult rat hippocampus. The Journal of Comparative Neurology, 443 (4), 346–367. 10.1002/cne.10118

Sevilla, D. F., Cabezas, C., Prada, A. N. O., Sánchez-Jiménez, A., & Buño, W. (2002). Selective muscarinic regulation of functional glutamatergic schaffer collateral synapses in rat CA1 pyramidal neurons. The Journal of Physiology, 545 (1), 51–63. 10.1113/jphysiol.2002.029165

Klausberger, T., Magill, P. J., Márton, L. F., Roberts, J. D. B., Cobden, P. M., Buzsáki, G., & Somogyi, P. (2003). Brain-state-and cell-type-specific firing of hippocampal interneurons in vivo. Nature, 421 (6925), 844–848. 10.1038/nature01374

Wierenga, C. J., & Wadman, W. J. (2003). Excitatory inputs to CA1 interneurons show selective synaptic dynamics. Journal of Neurophysiology, 90 (2), 811–821. 10.1152/jn.00865.2002

Agoston, M. K. (2004). Computer graphics and geometric modeling: Implementation and algorithms. Springer.

Klausberger, T., Márton, L. F., Baude, A., Roberts, J. D. B., Magill, P. J., & Somogyi, P. (2004). Spike timing of dendrite-targeting bistratified cells during hippocampal network oscillations in vivo. Nature Neuroscience, 7 (1), 41–47. 10.1038/nn1159

Biro, A. A. (2005). Quantal size is independent of the release probability at hippocampal excitatory synapses. Journal of Neuroscience, 25 (1), 223–232. 10.1523/JNEUROSCI.3688-04.2005

Buzsáki, G. (2005). Theta rhythm of navigation: Link between path integration and landmark navigation, episodic and semantic memory. Hippocampus, 15 (7), 827–840. 10.1002/hipo.20113

Klausberger, T. (2005). Complementary roles of cholecystokinin-and parvalbumin-expressing GABAergic neurons in hippocampal network oscillations. Journal of Neuroscience, 25 (42), 9782–9793. 10.1523/JNEUROSCI.3269-05.2005

Migliore, M., Ferrante, M., & Ascoli, G. A. (2005). Signal propagation in oblique dendrites of CA1 pyramidal cells. J. Neurophysiol., 94 (6), 4145–4155. 10.1152/jn.00521.2005

Siapas, A. G., Lubenov, E. V., & Wilson, M. A. (2005). Prefrontal phase locking to hippocampal theta oscillations. Neuron, 46 (1), 141–151. 10.1016/j.neuron.2005.02.028

Glickfeld, L. L., & Scanziani, M. (2006). Distinct timing in the activity of cannabinoid-sensitive and cannabinoid-insensitive basket cells. Nature Neuroscience, 9 (6), 807–815. 10.1038/nn1688

Hasselmo, M. E. (2006). The role of acetylcholine in learning and memory. Current Opinion in Neuro-biology, 16 (6), 710–715. 10.1016/j.conb.2006.09.002

Lawrence, J. J., Statland, J. M., Grinspan, Z. M., & McBain, C. J. (2006). Cell type-specific dependence of muscarinic signalling in mouse hippocampal stratum oriens interneurones: mAChR modulation of str. oriens interneurones. The Journal of Physiology, 570 (3), 595–610. 10.1113/jphysiol.2005.100875

Mercer, A., Bannister, A. P., & Thomson, A. M. (2006). Electrical coupling between pyramidal cells in adult cortical regions. Brain Cell Biol., 35 (1), 13–27. 10.1007/s11068-006-9005-9

Sasaki, T., Kimura, R., Tsukamoto, M., Matsuki, N., & Ikegaya, Y. (2006). Integrative spike dynamics of rat CA1 neurons: A multineuronal imaging study. The Journal of Physiology, 574, 195–208. 10.1113/jphysiol.2006.108480

Sniedovich, M. (2006). Dijkstra’s algorithm revisited: The dynamic programming connexion. Control and Cybernetics, 599–620. Retrieved August 2, 2022, from https://www.infona.pl//resource/bwmeta1.element.baztech-article-BAT5-0013-0005

Neu, A., Földy, C., & Soltesz, I. (2007). Postsynaptic origin of CB1-dependent tonic inhibition of GABA release at cholecystokinin-positive basket cell to pyramidal cell synapses in the CA1 region of the rat hippocampus. The Journal of Physiology, 578, 233–247. 10.1113/jphysiol.2006.115691

Wittner, L., Henze, D. A., Záborszky, L., & Buzsáki, G. (2007). Three-dimensional reconstruction of the axon arbor of a CA3 pyramidal cell recorded and filled in vivo. Brain Structure and Function, 212 (1), 75–83. 10.1007/s00429-007-0148-y

Ali, A. B., & Thomson, A. M. (2008). Synaptic alpha 5 subunit-containing GABAA receptors mediate IPSPs elicited by dendrite-preferring cells in rat neocortex. Cereb. Cortex, 18 (6), 1260–1271.

Elfant, D., Pál, B. Z., Emptage, N., & Capogna, M. (2008). Specific inhibitory synapses shift the balance from feedforward to feedback inhibition of hippocampal CA1 pyramidal cells. The European Journal of Neuroscience, 27 (1), 104–113. 10.1111/j.1460-9568.2007.06001.x

Fuentealba, P., Begum, R., Capogna, M., Jinno, S., Márton, L. F., Csicsvari, J., Thomson, A., Somogyi, P., & Klausberger, T. (2008). Ivy cells: A population of nitric-oxide-producing, slow-spiking GABAergic neurons and their involvement in hippocampal network activity. Neuron, 57 (6), 917–929. 10.1016/j.neuron.2008.01.034

Petilla Interneuron Nomenclature Group, Ascoli, G. A., Alonso-Nanclares, L., Anderson, S. A., Barrionuevo, G., Benavides-Piccione, R., Burkhalter, A., Buzsáki, G., Cauli, B., Defelipe, J., Fairén, A., Feldmeyer, D., Fishell, G., Fregnac, Y., Freund, T. F., Gardner, D., Gardner, E. P., Goldberg, J. H., Helmstaedter, M.,…Yuste, R. (2008). Petilla terminology: Nomenclature of features of GABAergic interneurons of the cerebral cortex. Nat. Rev. Neurosci., 9 (7), 557–568. 10.1038/nrn2402

Shah, M. M., Migliore, M., Valencia, I., Cooper, E. C., & Brown, D. A. (2008). Functional significance of axonal kv7 channels in hippocampal pyramidal neurons. Proc. Natl. Acad. Sci. U. S. A., 105 (22), 7869–7874. 10.1073/pnas.0802805105

Sirota, A., Montgomery, S., Fujisawa, S., Isomura, Y., Zugaro, M., & Buzsáki, G. (2008). Entrainment of neocortical neurons and gamma oscillations by the hippocampal theta rhythm. Neuron, 60 (4), 683–697. 10.1016/j.neuron.2008.09.014

Addison, P., Walker, J., & Guido, R. (2009). Time–frequency analysis of biosignals. IEEE Eng. Med. Biol. Mag., 28 (5), 14–29. 10.1109/MEMB.2009.934244

Anwar, H., Riachi, I., Hill, S., Schürmann, F., & Markram, H. (2009). An approach to capturing neuron morphological diversity. In Computational modeling methods for neuroscientists (pp. 211–232). The MIT Press. 10.7551/mitpress/9780262013277.003.0010

Goutagny, R., Jackson, J., & Williams, S. (2009). Self-generated theta oscillations in the hippocampus. Nature Neuroscience, 12 (12), 1491–1493. 10.1038/nn.2440

Hangya, B., Borhegyi, Z., Szilágyi, N., Freund, T. F., & Varga, V. (2009). GABAergic neurons of the medial septum lead the hippocampal network during theta activity. The Journal of Neuroscience: The Official Journal of the Society for Neuroscience, 29 (25), 8094–8102. 10.1523/JNEUROSCI.5665-08.2009

Ito, H. T., & Schuman, E. M. (2009). Distance-dependent homeostatic synaptic scaling mediated by a-type potassium channels. Front. Cell. Neurosci., 3, 15. 10.3389/neuro.03.015.2009

Pouille, F., Marin-Burgin, A., Adesnik, H., Atallah, B. V., & Scanziani, M. (2009). Input normalization by global feedforward inhibition expands cortical dynamic range. Nature Neuroscience, 12 (12), 1577–1585. 10.1038/nn.2441

Prieto, G. A., Parker, R. L., & Vernon, F. L., III. (2009). A fortran 90 library for multitaper spectrum analysis. Comput. Geosci., 35 (8), 1701–1710. 10.1016/j.cageo.2008.06.007

Rockström, J., Steffen, W., Noone, K., Persson, Å., Chapin, F. S., Lambin, E. F., Lenton, T. M., Scheffer, M., Folke, C., Schellnhuber, H. J., Nykvist, B., de Wit, C. A., Hughes, T., van der Leeuw, S., Rodhe, H., Sörlin, S., Snyder, P. K., Costanza, R., Svedin, U.,…Foley, J. A. (2009). A safe operating space for humanity. Nature, 461 (7263), 472–475. 10.1038/461472a

Ascoli, G. A., Gasparini, S., Medinilla, V., & Migliore, M. (2010). Local control of postinhibitory rebound spiking in CA1 pyramidal neuron dendrites. The Journal of Neuroscience: The Official Journal of the Society for Neuroscience, 30 (18), 6434–6442. 10.1523/JNEUROSCI.4066-09.2010

Bokil, H., Andrews, P., Kulkarni, J. E., Mehta, S., & Mitra, P. P. (2010). Chronux: A platform for analyzing neural signals. J. Neurosci. Methods, 192 (1), 146–151. 10.1016/j.jneumeth.2010.06.020

Buchanan, K. A., Petrovic, M. M., Chamberlain, S. E., Marrion, N. V., & Mellor, J. R. (2010). Facilitation of long-term potentiation by muscarinic m1 receptors is mediated by inhibition of SK channels. Neuron, 68 (5), 948–963. 10.1016/j.neuron.2010.11.018

Cutsuridis, V., Cobb, S., & Graham, B. P. (2010). Encoding and retrieval in a model of the hippocampal CA1 microcircuit. Hippocampus, 20 (3), 423–446. 10.1002/hipo.20661

Földy, C., Lee, S.-H., Morgan, R. J., & Soltesz, I. (2010). Regulation of fast-spiking basket cell synapses by the chloride channel ClC-2. Nature Neuroscience, 13 (9), 1047–1049. 10.1038/nn.2609

Giaume, C., Koulakoff, A., Roux, L., Holcman, D., & Rouach, N. (2010). Astroglial networks: A step further in neuroglial and gliovascular interactions. Nat. Rev. Neurosci., 11 (2), 87–99. 10.1038/nrn2757

Hu, H., Martina, M., & Jonas, P. (2010). Dendritic mechanisms underlying rapid synaptic activation of fast-spiking hippocampal interneurons. Science (New York, N.Y.), 327 (5961), 52–58. 10.1126/science.1177876

Lee, S.-H., Foldy, C., & Soltesz, I. (2010). Distinct endocannabinoid control of GABA release at perisomatic and dendritic synapses in the hippocampus. Journal of Neuroscience, 30 (23), 7993–8000. 10.1523/jneurosci.6238-09.2010

Morse, T. M., Carnevale, N. T., Mutalik, P. G., Migliore, M., & Shepherd, G. M. (2010). Abnormal excitability of oblique dendrites implicated in early alzheimer’s: A computational study. Frontiers in Neural Circuits, 4, 16. 10.3389/fncir.2010.00016

Szabó, G. G., Holderith, N., Gulyás, A. I., Freund, T. F., & Hájos, N. (2010). Distinct synaptic properties of perisomatic inhibitory cell types and their different modulation by cholinergic receptor activation in the CA3 region of the mouse hippocampus: Synaptic properties of perisomatic interneurons. European Journal of Neuroscience, 31 (12), 2234–2246. 10.1111/j.1460-9568.2010.07292.x

Zhang, H., Lin, S.-C., & Nicolelis, M. A. L. (2010). Spatiotemporal coupling between hippocampal acetylcholine release and theta oscillations in vivo. The Journal of Neuroscience, 30 (40), 13431– 13440. 10.1523/jneurosci.1144-10.2010

Ali, A. B. (2011). CB1 modulation of temporally distinct synaptic facilitation among local circuit interneurons mediated by n-type calcium channels in CA1. Journal of Neurophysiology, 105 (3), 1051–1062. 10.1152/jn.00831.2010

Cea-del Rio, C. A., Lawrence, J. J., Erdelyi, F., Szabó, G., & McBain, C. J. (2011). Cholinergic modulation amplifies the intrinsic oscillatory properties of CA1 hippocampal cholecystokinin-positive interneurons: Cholinergic modulation and CA1 cholecystokinin-positive interneurons. The Journal of Physiology, 589 (3), 609–627. 10.1113/jphysiol.2010.199422

Dasari, S., & Gulledge, A. T. (2011). M1 and m4 receptors modulate hippocampal pyramidal neurons. Journal of Neurophysiology, 105 (2), 779–792. 10.1152/jn.00686.2010

Deguchi, Y., Donato, F., Galimberti, I., Cabuy, E., & Caroni, P. (2011). Temporally matched subpopulations of selectively interconnected principal neurons in the hippocampus. Nature Neuroscience, 14 (4), 495–504. 10.1038/nn.2768

Mizuseki, K., Diba, K., Pastalkova, E., & Buzsáki, G. (2011). Hippocampal CA1 pyramidal cells form functionally distinct sublayers. Nature Neuroscience, 14 (9), 1174–1181. 10.1038/nn.2894

Slomianka, L., Amrein, I., Knuesel, I., Sørensen, J. C., & Wolfer, D. P. (2011). Hippocampal pyramidal cells: The reemergence of cortical lamination. Brain Structure & Function, 216 (4), 301–317. 10.1007/s00429-011-0322-0

Belluscio, M. A., Mizuseki, K., Schmidt, R., Kempter, R., & Buzsáki, G. (2012). Cross-frequency phase-phase coupling between theta and gamma oscillations in the hippocampus. J. Neurosci., 32 (2), 423–435. 10.1523/JNEUROSCI.4122-11.2012

Graves, A. R., Moore, S. J., Bloss, E. B., Mensh, B. D., Kath, W. L., & Spruston, N. (2012). Hippocampal pyramidal neurons comprise two distinct cell types that are countermodulated by metabotropic receptors. Neuron, 76 (4), 776–789. 10.1016/j.neuron.2012.09.036

Potworowski, J., Jakuczun, W., Leski, S., & Wojcik, D. (2012). Kernel current source density method. Neural Comput., 24 (2), 541–575. 10.1162/NECO\_a\_00236

Ropireddy, D., Bachus, S., & Ascoli, G. (2012). Non-homogeneous stereological properties of the rat hippocampus from high-resolution 3d serial reconstruction of thin histological sections. Neuroscience, 205, 91–111. 10.1016/j.neuroscience.2011.12.055

Szirmai, I., Buzsáki, G., & Kamondi, A. (2012). 120 years of hippocampal schaffer collaterals. Hippocampus, 22 (7), 1508–1516. 10.1002/hipo.22001

Takács, V. T., Klausberger, T., Somogyi, P., Freund, T. F., & Gulyás, A. I. (2012). Extrinsic and local glutamatergic inputs of the rat hippocampal CA1 area differentially innervate pyramidal cells and interneurons. Hippocampus, 22 (6), 1379–1391. 10.1002/hipo.20974

Bezaire, M. J., & Soltesz, I. (2013). Quantitative assessment of CA1 local circuits: Knowledge base for interneuron-pyramidal cell connectivity: Quantitative assessment of ca1 local circuits. Hippocampus, 23 (9), 751–785. 10.1002/hipo.22141

Colgin, L. L. (2013). Mechanisms and functions of theta rhythms. Annual Review of Neuroscience, 36, 295–312. 10.1146/annurev-neuro-062012-170330

Hood, L., & Rowen, L. (2013). The human genome project: Big science transforms biology and medicine. Genome Medicine, 5 (9), 79. 10.1186/gm483

Le Roux, N., Cabezas, C., Böhm, U. L., & Poncer, J. C. (2013). Input-specific learning rules at excitatory synapses onto hippocampal parvalbumin-expressing interneurons. The Journal of Physiology, 591 (7), 1809–1822. 10.1113/jphysiol.2012.245852

Teles-Grilo Ruivo, L. M., & Mellor, J. R. (2013). Cholinergic modulation of hippocampal network function. Front. Synaptic Neurosci., 5, 2. 10.3389/fnsyn.2013.00002

Zemankovics, R., Veres, J. M., Oren, I., & Hajos, N. (2013). Feedforward inhibition underlies the propagation of cholinergically induced gamma oscillations from hippocampal CA3 to CA1. Journal of Neuroscience, 33 (30), 12337–12351. 10.1523/JNEUROSCI.3680-12.2013

Lee, S.-H., Marchionni, I., Bezaire, M., Varga, C., Danielson, N., Lovett-Barron, M., Losonczy, A., & Soltesz, I. (2014). Parvalbumin-positive basket cells differentiate among hippocampal pyramidal cells. Neuron, 82 (5), 1129–1144. 10.1016/j.neuron.2014.03.034

Pietersen, A. N. J., Ward, P. D., Hagger-Vaughan, N., Wiggins, J., Jefferys, J. G. R., & Vreugdenhil, M. (2014). Transition between fast and slow gamma modes in rat hippocampus area CA1 in vitro is modulated by slow CA3 gamma oscillations. The Journal of Physiology, 592 (4), 605–620. 10.1113/jphysiol.2013.263889

Sun, Y., Nguyen, A. Q., Nguyen, J. P., Le, L., Saur, D., Choi, J., Callaway, E. M., & Xu, X. (2014). Cell-type-specific circuit connectivity of hippocampal CA1 revealed through cre-dependent rabies tracing. Cell Reports, 7 (1), 269–280. 10.1016/j.celrep.2014.02.030

Szigeti, B., Gleeson, P., Vella, M., Khayrulin, S., Palyanov, A., Hokanson, J., Currie, M., Cantarelli, M., Idili, G., & Larson, S. (2014). OpenWorm: An open-science approach to modeling caenorhabditis elegans. Front. Comput. Neurosci., 8, 137. 10.3389/fncom.2014.00137

Vandecasteele, M., Varga, V., Berényi, A., Papp, E., Barthó, P., Venance, L., Freund, T. F., & Buzsáki, G. (2014). Optogenetic activation of septal cholinergic neurons suppresses sharp wave ripples and enhances theta oscillations in the hippocampus. Proceedings of the National Academy of Sciences, 111 (37), 13535–13540. 10.1073/pnas.1411233111

Yang, S., Yang, S., Moreira, T., Hoffman, G., Carlson, G. C., Bender, K. J., Alger, B. E., & Tang, C.-M. (2014). Interlamellar CA1 network in the hippocampus. Proceedings of the National Academy of Sciences of the United States of America, 111 (35), 12919–12924. 10.1073/pnas.1405468111

Aad, G., & Abbott, B. (2015). Combined measurement of the Higgs boson mass in pp collisions at 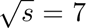 and 8 TeV with the ATLAS and CMS experiments. Physical Review Letters, 114 (19). 10.1103/physrevlett.114.191803

Markram, H., Muller, E., Ramaswamy, S., Reimann, M. W., Abdellah, M., Sanchez, C. A., Ailamaki, A., Alonso-Nanclares, L., Antille, N., Arsever, S., Kahou, G. A. A., Berger, T. K., Bilgili, A., Buncic, N., Chalimourda, A., Chindemi, G., Courcol, J.-D., Delalondre, F., Delattre, V.,…Gidon, A. (2015). Reconstruction and simulation of neocortical microcircuitry. Cell, 163 (2), 456–492. 10.1016/j.cell.2015.09.029

Milstein, A. D., Bloss, E. B., Apostolides, P. F., Vaidya, S. P., Dilly, G. A., Zemelman, B. V., & Magee, J. C. (2015). Inhibitory gating of input comparison in the CA1 microcircuit. Neuron, 87 (6), 1274–1289. 10.1016/j.neuron.2015.08.025

Reimann, M. W., King, J. G., Muller, E. B., Ramaswamy, S., & Markram, H. (2015). An algorithm to predict the connectome of neural microcircuits. Frontiers in Computational Neuroscience, 9. 10.3389/fncom.2015.00120

Wheeler, D. W., White, C. M., Rees, C. L., Komendantov, A. O., Hamilton, D. J., & Ascoli, G. A. (2015). Hippocampome.org: A knowledge base of neuron types in the rodent hippocampus (F. K. Skinner, Ed.) [Publisher: eLife Sciences Publications, Ltd]. eLife, 4, e09960. 10.7554/eLife.09960

Bezaire, M. J., Raikov, I., Burk, K., Vyas, D., & Soltesz, I. (2016). Interneuronal mechanisms of hippocampal theta oscillations in a full-scale model of the rodent CA1 circuit. eLife, 5. 10.7554/eLife.18566

Malik, R., Dougherty, K. A., Parikh, K., Byrne, C., & Johnston, D. (2016). Mapping the electrophysiological and morphological properties of CA1 pyramidal neurons along the longitudinal hippocampal axis. Hippocampus, 26 (3), 341–361. 10.1002/hipo.22526

Van Geit, W., Gevaert, M., Chindemi, G., Rössert, C., Courcol, J.-D., Muller, E. B., Schürmann, F., Segev, I., & Markram, H. (2016). BluePyOpt: Leveraging open source software and cloud infrastructure to optimise model parameters in neuroscience. Front. Neuroinform., 10, 17. 10.3389/fninf.2016.00017

Dannenberg, H., Young, K., & Hasselmo, M. (2017). Modulation of hippocampal circuits by muscarinic and nicotinic receptors. Front. Neural Circuits, 11, 102. 10.3389/fncir.2017.00102

Ferguson, K. A., Chatzikalymniou, A. P., & Skinner, F. K. (2017). Combining theory, model, and experiment to explain how intrinsic theta rhythms are generated in an in vitro whole hippocampus preparation without oscillatory inputs. eNeuro, 4 (4). 10.1523/ENEURO.0131-17.2017

Gal, E., London, M., Globerson, A., Ramaswamy, S., Reimann, M. W., Muller, E., Markram, H., & Segev, I. (2017). Rich cell-type-specific network topology in neocortical microcircuitry. Nature Neuroscience, 20 (7), 1004–1013. 10.1038/nn.4576

Pelkey, K. A., Chittajallu, R., Craig, M. T., Tricoire, L., Wester, J. C., & McBain, C. J. (2017). Hippocampal GABAergic inhibitory interneurons. Physiol. Rev., 97 (4), 1619–1747. 10.1152/physrev.00007.2017

Abdellah, M., Hernando, J., Eilemann, S., Lapere, S., Antille, N., Markram, H., & Schürmann, F. (2018). NeuroMorphoVis: A collaborative framework for analysis and visualization of neuronal morphology skeletons reconstructed from microscopy stacks. Bioinformatics (Oxford, England), 34 (13), i574–i582. 10.1093/bioinformatics/bty231

Denker, M., Yegenoglu, A., & Grün, S. (2018). Collaborative HPC-enabled workflows on the HBP collaboratory using the elephant framework [Publisher: G-Node]. 10.12751/INCF.NI2018.0019

Dumas, T. C., Uttaro, M. R., Barriga, C., Brinkley, T., Halavi, M., Wright, S. N., Ferrante, M., Evans, R. C., Hawes, S. L., & Sanders, E. M. (2018). Removal of area CA3 from hippocampal slices induces postsynaptic plasticity at schaffer collateral synapses that normalizes CA1 pyramidal cell discharge. Neuroscience Letters, 678, 55–61. 10.1016/j.neulet.2018.05.011

Economides, G., Falk, S., & Mercer, A. (2018). Biocytin recovery and 3D reconstructions of filled hippocampal CA2 interneurons. J. Vis. Exp., (141). 10.3791/58592

Gerkin, R. C., Jarvis, R. J., & Crook, S. M. (2018). Towards systematic, data-driven validation of a collaborative, multi-scale model of caenorhabditis elegans. Philos. Trans. R. Soc. Lond. B Biol. Sci., 373 (1758). 10.1098/rstb.2017.0381

Kanari, L., Dłotko, P., Scolamiero, M., Levi, R., Shillcock, J., Hess, K., & Markram, H. (2018). A topological representation of branching neuronal morphologies. Neuroinformatics, 16 (1), 3–13. 10.1007/s12021-017-9341-1

Migliore, R., Lupascu, C. A., Bologna, L. L., Romani, A., Courcol, J.-D., Antonel, S., Van Geit, W. A. H., Thomson, A. M., Mercer, A., Lange, S., Falck, J., Rössert, C. A., Shi, Y., Hagens, O., Pezzoli, M., Freund, T. F., Kali, S., Muller, E. B., Schürmann, F.,…Migliore, M. (2018). The physiological variability of channel density in hippocampal CA1 pyramidal cells and interneurons explored using a unified data-driven modeling workflow (W. W. Lytton, Ed.). PLOS Computational Biology, 14 (9), e1006423. 10.1371/journal.pcbi.1006423

Müller, C., & Remy, S. (2018). Septo–hippocampal interaction. Cell and Tissue Research, 373 (3), 565–575. 10.1007/s00441-017-2745-2

Soltesz, I., & Losonczy, A. (2018). CA1 pyramidal cell diversity enabling parallel information processing in the hippocampus [Number: 4 Publisher: Nature Publishing Group]. Nature Neuroscience, 21 (4), 484–493. 10.1038/s41593-018-0118-0

Barros-Zulaica, N., Rahmon, J., Chindemi, G., Perin, R., Markram, H., Muller, E., & Ramaswamy, S. (2019). Estimating the readily-releasable vesicle pool size at synaptic connections in the neocortex. Frontiers in Synaptic Neuroscience, 11, 29. 10.3389/fnsyn.2019.00029

Colangelo, C., Shichkova, P., Keller, D., Markram, H., & Ramaswamy, S. (2019). Cellular, synaptic and network effects of acetylcholine in the neocortex. Frontiers in Neural Circuits, 13, 24. 10.3389/fncir.2019.00024

Cornford, J. H., Mercier, M. S., Leite, M., Magloire, V., Häusser, M., & Kullmann, D. M. (2019). Dendritic NMDA receptors in parvalbumin neurons enable strong and stable neuronal assemblies (M. Bartos, G. L. Westbrook, C.-C. Lien, & J. C. Poncer, Eds.). eLife, 8, e49872. 10.7554/eLife.49872

Fan, X., & Markram, H. (2019). A brief history of simulation neuroscience. Front. Neuroinform., 13, 32. 10.3389/fninf.2019.00032

Kanari, L., Ramaswamy, S., Shi, Y., Morand, S., Meystre, J., Perin, R., Abdellah, M., Wang, Y., Hess, K., & Markram, H. (2019). Objective morphological classification of neocortical pyramidal cells. Cereb. Cortex, 29 (4), 1719–1735. 10.1093/cercor/bhy339

Ecker, A., Romani, A., Sáray, S., Káli, S., Migliore, M., Falck, J., Lange, S., Mercer, A., Thomson, A. M., Muller, E., Reimann, M. W., & Ramaswamy, S. (2020). Data-driven integration of hippocampal CA1 synaptic physiology in silico. Hippocampus, 30 (11), 1129–1145. 10.1002/hipo.23220

Goyal, A., Miller, J., Qasim, S. E., Watrous, A. J., Zhang, H., Stein, J. M., Inman, C. S., Gross, R. E., Willie, J. T., Lega, B., Lin, J.-J., Sharan, A., Wu, C., Sperling, M. R., Sheth, S. A., McKhann, G. M., Smith, E. H., Schevon, C., & Jacobs, J. (2020). Functionally distinct high and low theta oscillations in the human hippocampus. Nature Communications, 11 (1), 2469. 10.1038/s41467-020-15670-6

Hasselmo, M. E., Alexander, A. S., Dannenberg, H., & Newman, E. L. (2020). Overview of computational models of hippocampus and related structures: Introduction to the special issue. Hippocampus, 30 (4), 295–301. 10.1002/hipo.23201

Moradi, K., & Ascoli, G. A. (2020). A comprehensive knowledge base of synaptic electrophysiology in the rodent hippocampal formation. Hippocampus, 30 (4), 314–331. 10.1002/hipo.23148

Wang, Q., Ding, S.-L., Li, Y., Royall, J., Feng, D., Lesnar, P., Graddis, N., Naeemi, M., Facer, B., Ho, A., Dolbeare, T., Blanchard, B., Dee, N., Wakeman, W., Hirokawa, K. E., Szafer, A., Sunkin, S. M., Oh, S. W., Bernard, A.,…Ng, L. (2020). The allen mouse brain common coordinate framework: A 3D reference atlas. Cell, 181 (4), 936–953.e20. 10.1016/j.cell.2020.04.007

Yu, G. J., Bouteiller, J.-M. C., & Berger, T. W. (2020). Topographic organization of correlation along the longitudinal and transverse axes in rat hippocampal CA3 due to excitatory afferents. Front. Comput. Neurosci., 14, 588881. 10.3389/fncom.2020.588881

Giacopelli, G., Tegolo, D., Spera, E., & Migliore, M. (2021). On the structural connectivity of large-scale models of brain networks at cellular level. Scientific Reports, 11 (1), 4345. 10.1038/s41598-021-83759-z

Palacios-Filardo, J., Udakis, M., Brown, G. A., Tehan, B. G., Congreve, M. S., Nathan, P. J., Brown, A. J. H., & Mellor, J. R. (2021). Acetylcholine prioritises direct synaptic inputs from entorhinal cortex to CA1 by differential modulation of feedforward inhibitory circuits. Nature Communications, 12 (1), 5475. 10.1038/s41467-021-25280-5

Sanchez-Aguilera, A., Wheeler, D. W., Jurado-Parras, T., Valero, M., Nokia, M. S., Cid, E., Fernandez-Lamo, I., Sutton, N., Garcia-Rincon, D., de la Prida, L. M., & Ascoli, G. A. (2021). An update to hippocampome.org by integrating single-cell phenotypes with circuit function in vivo. PLoS biology, 19 (5), e3001213. 10.1371/journal.pbio.3001213

Sáray, S., Rössert, C. A., Appukuttan, S., Migliore, R., Vitale, P., Lupascu, C. A., Bologna, L. L., Van Geit, W., Romani, A., Davison, A. P., Muller, E., Freund, T. F., & Káli, S. (2021). HippoUnit: A software tool for the automated testing and systematic comparison of detailed models of hippocampal neurons based on electrophysiological data. PLoS Comput. Biol., 17 (1), e1008114. 10.1371/journal.pcbi.1008114

Sutton, N. M., & Ascoli, G. A. (2021). Spiking neural networks and hippocampal function: A web-accessible survey of simulations, modeling methods, and underlying theories. Cognitive Systems Research, 70, 80–92. 10.1016/j.cogsys.2021.07.008

Yang, D., Ding, C., Qi, G., & Feldmeyer, D. (2021). Cholinergic and adenosinergic modulation of synaptic release. Neuroscience, 456, 114–130. 10.1016/j.neuroscience.2020.06.006

Zisis, E., Keller, D., Kanari, L., Arnaudon, A., Gevaert, M., Delemontex, T., Coste, B., Foni, A., Ab-dellah, M., Calı, C., Hess, K., Magistretti, P. J., Schürmann, F., & Markram, H. (2021). Digital reconstruction of the Neuro-Glia-Vascular architecture. Cereb. Cortex, 31 (12), 5686–5703. 10.1093/cercor/bhab254

Awile, O., Kumbhar, P., Cornu, N., Dura-Bernal, S., King, J. G., Lupton, O., Magkanaris, I., McDougal, R. A., Newton, A. J. H., Pereira, F., Savulescu, A., Carnevale, N. T., Lytton, W. W., Hines, M. L., & Schurmann, F. (2022). Modernizing the NEURON simulator for sustainability, portability, and performance. Front. Neuroinform., 16, 884046. 10.3389/fninf.2022.884046

Palacios, J., Lida Kanari, Zisis, E., MikeG, Benoit Coste, Asanin-Epfl, Vanherpe, L., Jdcourcol, Arnaudon, A., Berchet, A., Haleepfl, Getta, P., Povolotsky, A. V., A Sato, Alex4200, Bbpgithubaudit, Ficarelli, G., Amsalem, O., Stefanoantonel, & Tomdele. (2022). Bluebrain/neurom: V3.2.2. 10.5281/ZENODO.597333

Reimann, M. W., Bolaños-Puchet, S., Courcol, J.-D., Santander, D. E., Arnaudon, A., Coste, B., Delemontex, T., Devresse, A., Dictus, H., Dietz, A., Ecker, A., Favreau, C., Ficarelli, G., Gevaert, M., Hernando, J. B., Herttuainen, J., Isbister, J. B., Kanari, L., Keller, D.,…Markram, H. (2022). Modeling and Simulation of Rat Non-Barrel Somatosensory Cortex. Part I: Modeling Anatomy. bioRxiv, 2022.08.11.503144. 10.1101/2022.08.11.503144

Romani, A., Schürmann, F., Markram, H., & Migliore, M. (2022). Reconstruction of the hippocampus. Advances in Experimental Medicine and Biology, 1359, 261–283. 10.1007/978-3-030-89439-9\_11

Iavarone, E., Simko, J., Shi, Y., Bertschy, M., Garcıa-Amado, M., Litvak, P., Kaufmann, A.-K., O’Reilly, C., Amsalem, O., Abdellah, M., Chevtchenko, G., Coste, B., Courcol, J.-D., Ecker, A., Favreau, C., Fleury, A. C., Van Geit, W., Gevaert, M., Guerrero, N. R.,…Hill, S. L. (2023). Thalamic control of sensory processing and spindles in a biophysical somatosensory thalamoreticular circuit model of wakefulness and sleep. Cell Rep., 42 (3), 112200. 10.1016/j.celrep.2023.112200

Isbister, J. B., Ecker, A., Pokorny, C., Bolaños-Puchet, S., Santander, D. E., Arnaudon, A., Awile, O., Barros-Zulaica, N., Alonso, J. B., Boci, E., Chindemi, G., Courcol, J.-D., Damart, T., Delemontex, T., Dietz, A., Ficarelli, G., Gevaert, M., Herttuainen, J., Ivaska, G.,…Reimann, M. W. (2023). Modeling and simulation of neocortical micro-and mesocircuitry. part II: Physiology and experimentation. bioRxiv, 2023.05.17.541168. 10.1101/2023.05.17.541168

