## Supplementary material for "Community-based Reconstruction and Simulation of a Full-scale Model of Region CA1 of Rat Hippocampus"

### Supplementary Figures

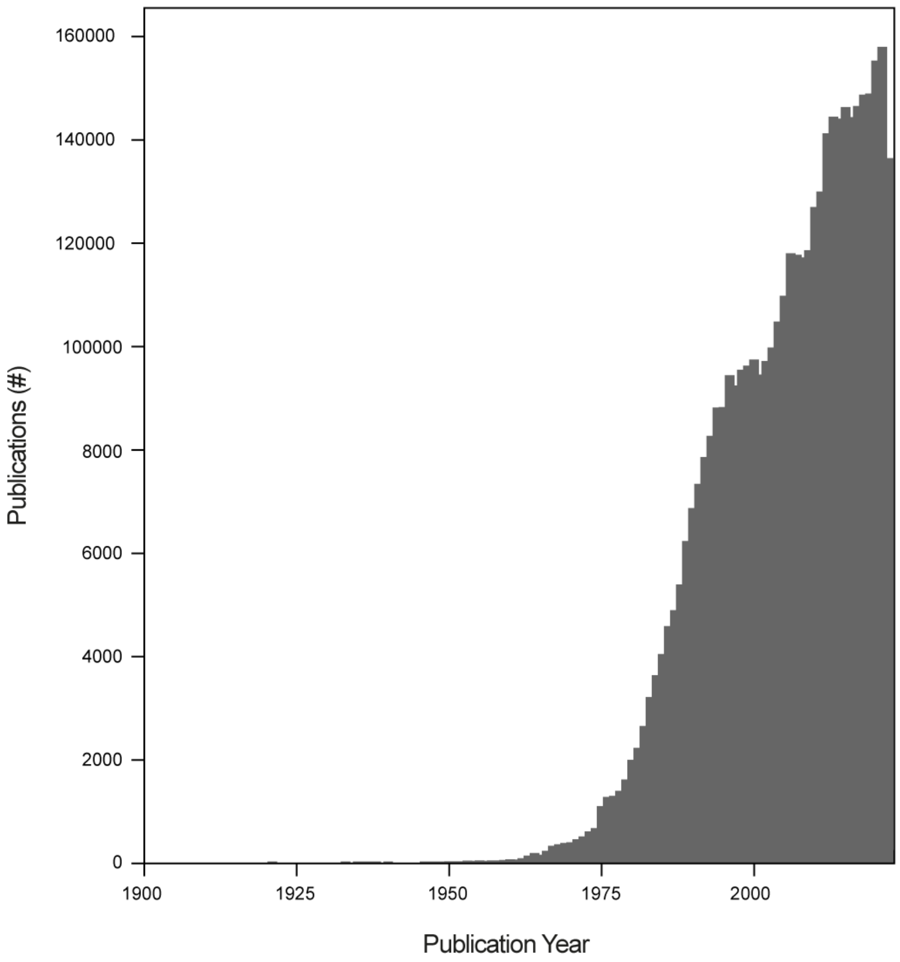


Figure S1: **Scientific publications on hippocampus progressively increase**. Counts per year of the number of publications based on a pubmed search for 'hippocampus|cornu Ammonis|CA1|CA2|CA3' from 1900-present indicates the large size of neuroscientific community researching hippocampus.

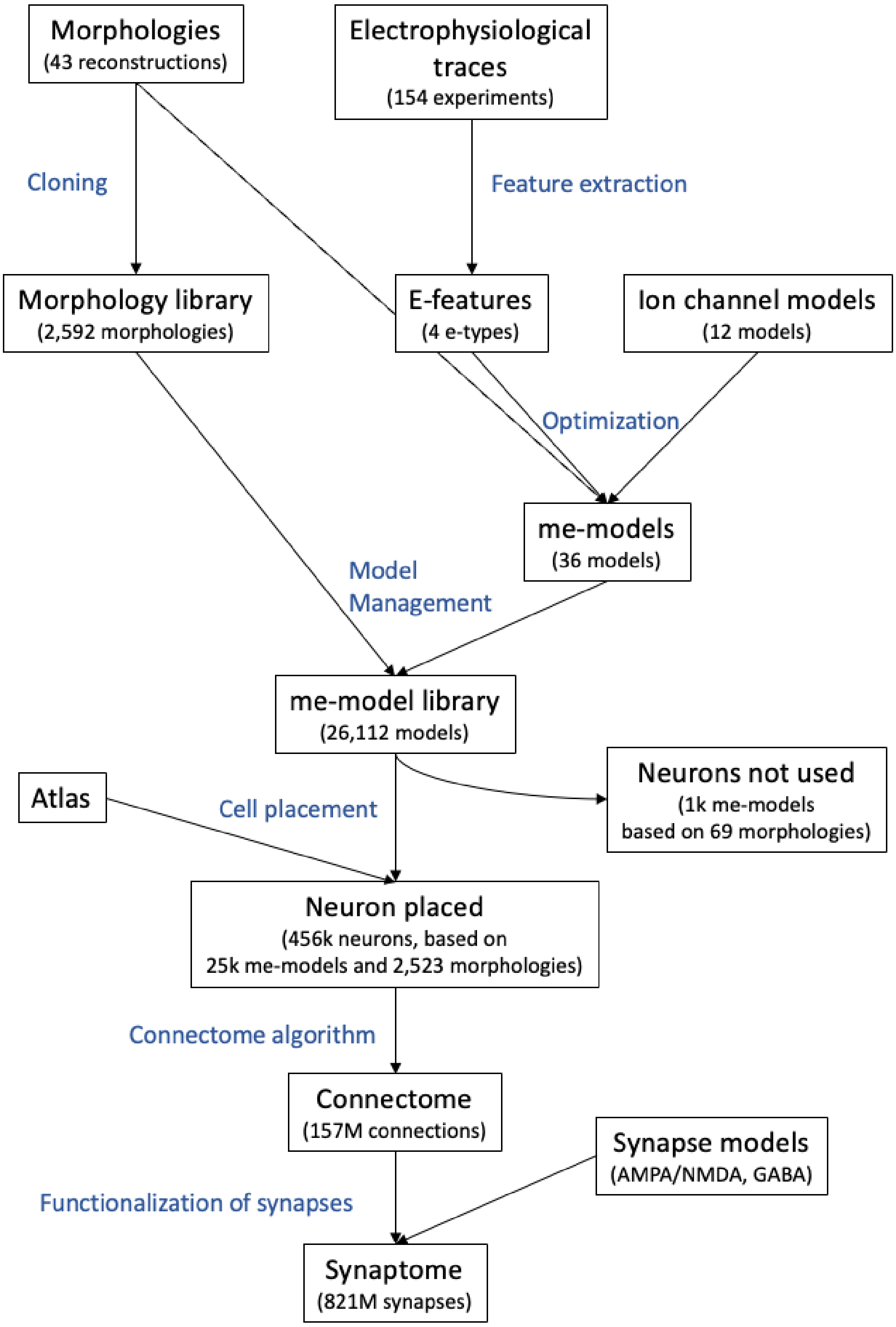


Figure S2: **Circuit building workflow**. Simplified workflow of the circuit building. Boxes represent the different building blocks, while blue labels are the operations between blocks.

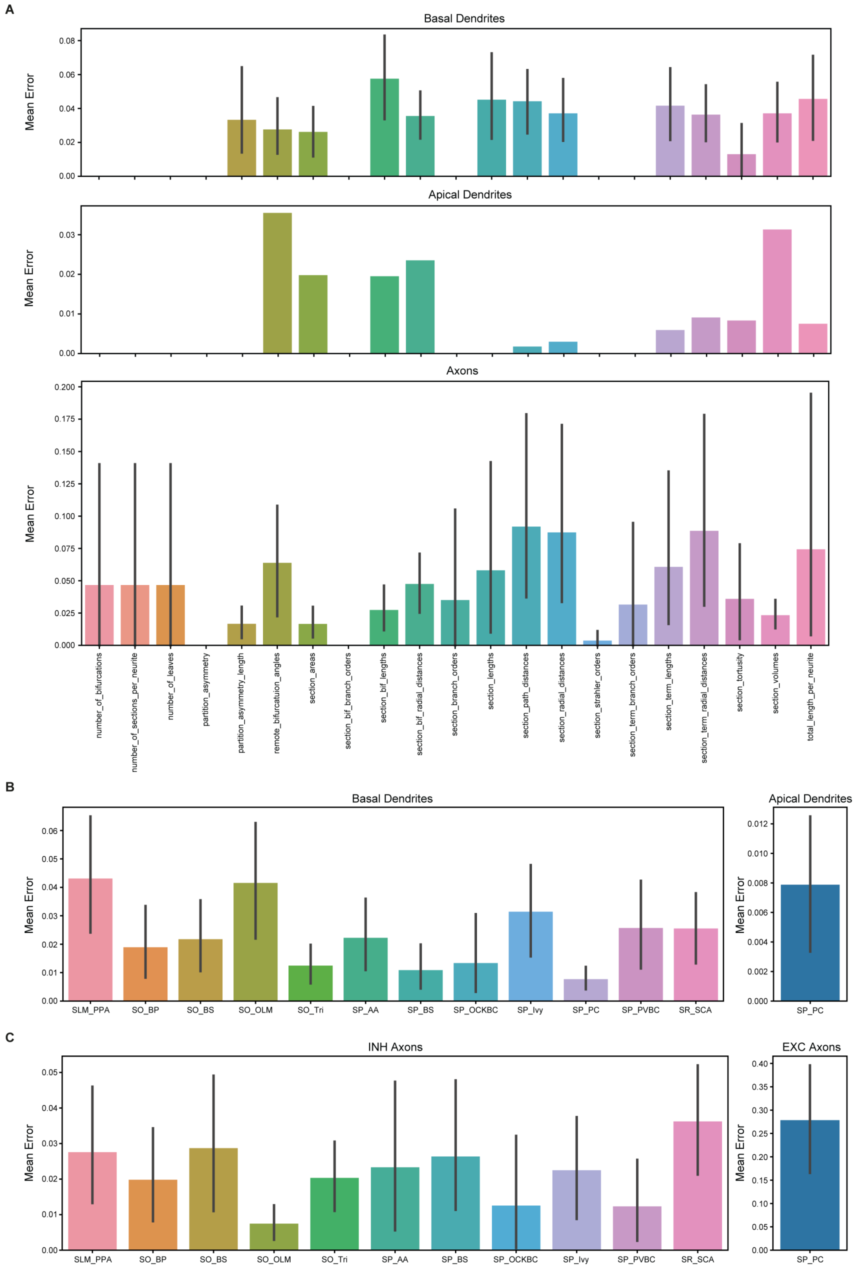


Figure S3: **Validation of cloning**. Validations of cloning methods with a similarity metric (differences between median values divided by variance). A. The mean scores for 21 unique morphometrics averaged across m-types shown for basal and apical dendrites, and axons in each row. B. The same scoring grouped by m-types instead of metrics for basal and apical dendrites. Since inhibitory neurons did not have apical dendrites, we show its score only for pyramidal cells. C. Similar to B, but calculated for axons. Excitatory and inhibitory axons are grouped separately.

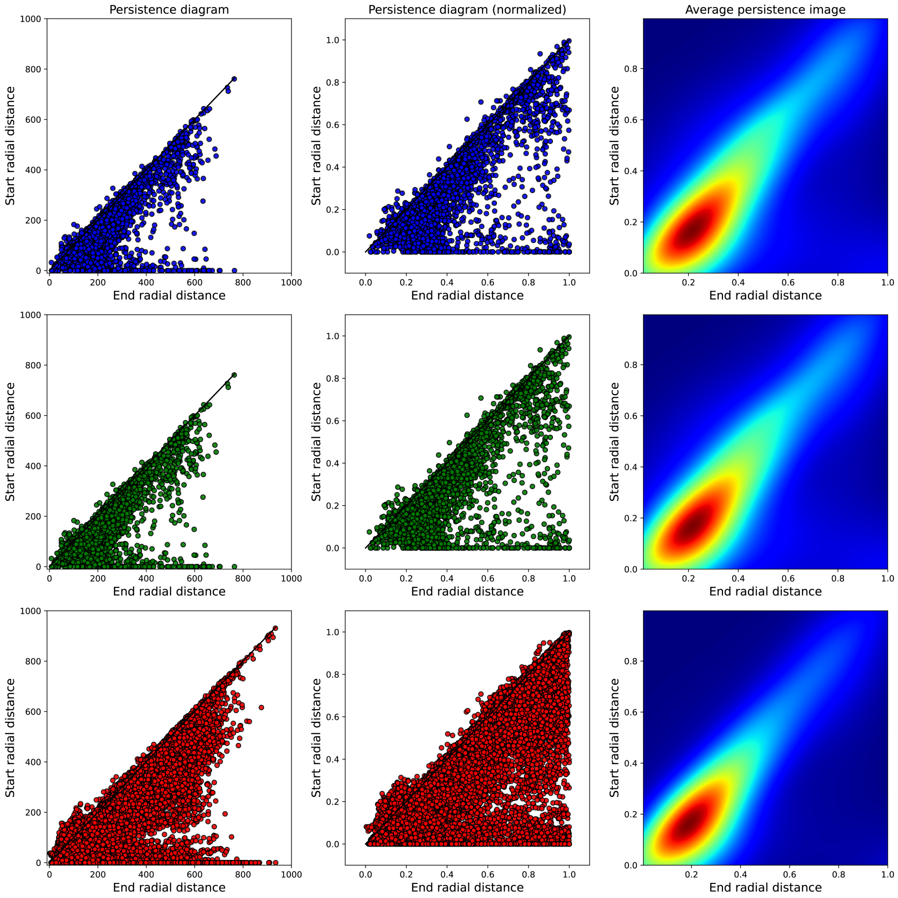


Figure S4: **Increasing morphological diversity**. A. Persistence diagrams of original (blue), repaired (green) and cloned (red) dendrites. Persistence diagrams encode the start (y-axis) and end (x-axis) radial distances of all dendritic trees in the respective populations of neurons. B. Persistence diagrams of normalized radial distances (to one) from original (blue), repaired (green) and cloned (red) dendrites. C. Persistence images are the Gaussian averages of the respective persistence diagrams (B) from original (top), repaired (middle) and cloned (bottom) dendrites. Calculations are computed for the all cell groups. For more detailed analyses, see Figures S5 and S6.

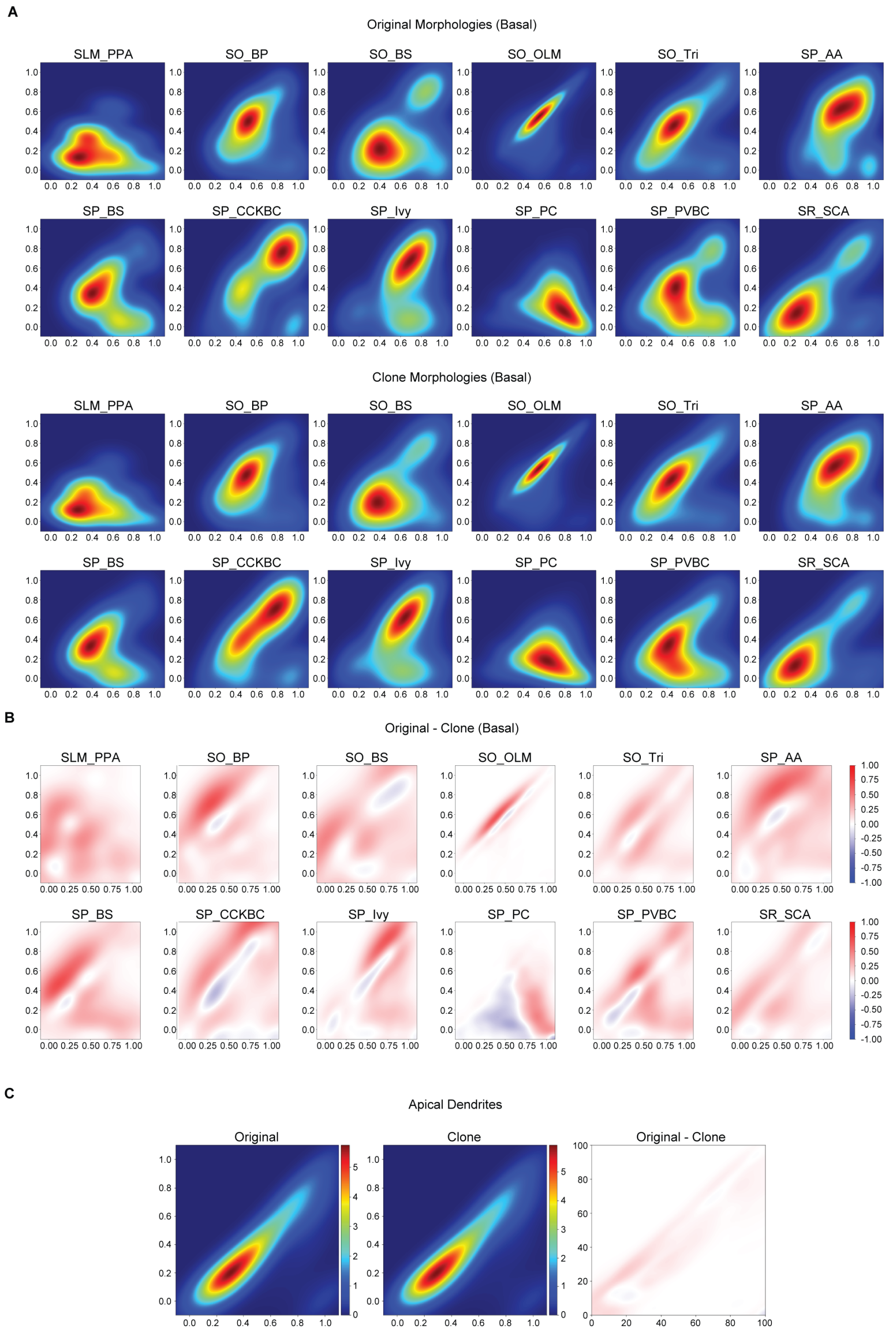


Figure S5: **Persistent images for the basal and apical dendrites of each m-type**. A. Persistent images averaged across each m-type for original morphologies and for their cloned counterparts with respect to the basal dendrites, and (B) their differences. C. The same process applied for apical dendrites which was only present in the excitatory cell group (e.g., pyramidal cells). Note that the radial distance is synonymous with Euclidean distance, and should not be mixed with hippocampal radial axis.

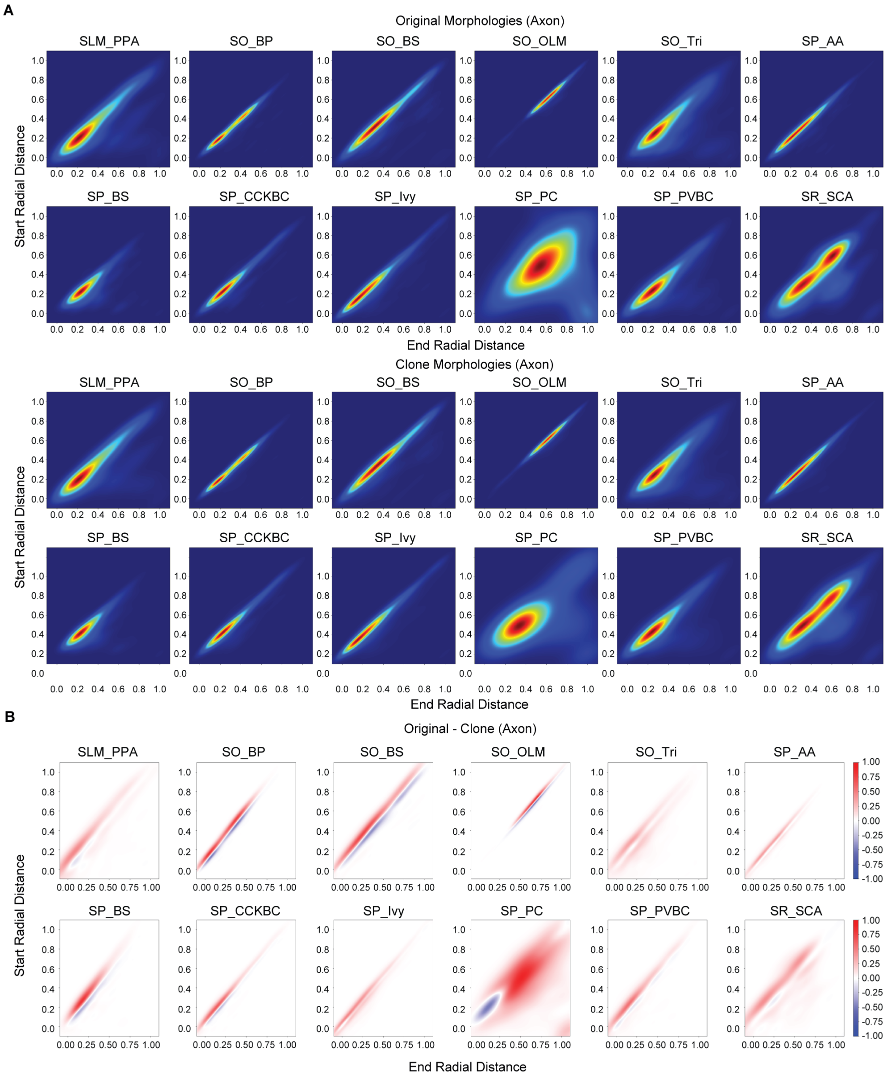


Figure S6: **Persistent images for the axons of each m-type**. Persistent images averaged across each cell type for A. original axons, B. cloned versions, and C. their difference.

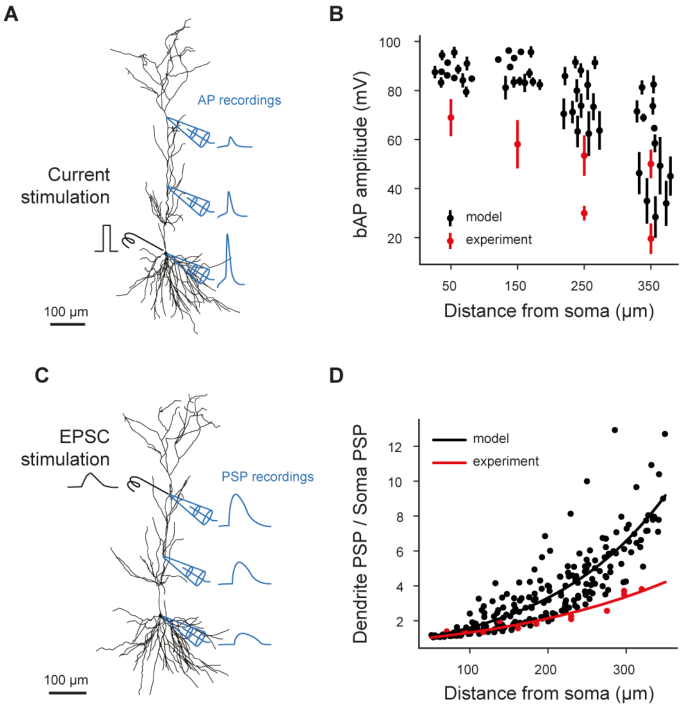


Figure S7: **Validation of single-cell models**. A. Graphical illustration of the back-propagating action potential protocol. In a pyramidal cell model, the soma is stimulated to elicit an AP, which is then measured at different distances from the soma. B. *In silico* measurements (black dots) are reported and compared with experimental data from Golding et al. (2001) (red dots and whiskers, indicating mean and standard deviation). C. Graphical illustration of the post-synaptic potential attenuation protocol. Dendrites on the apical trunk of a pyramidal cell are stimulated with bi-exponential currents; then, PSP's amplitudes are measured as it travels toward the soma. D. *In silico* measurements of PSP attenuation (black dots) are expressed as a ratio between the PSP measured in the dendrite and the one measured in the soma. Pyramidal cell models are compared with experimental data from Magee and Cook (2000). The two distributions have been fitted with exponential equations, resulting in the following space constants: $\tau$model $= 155.6\,\mu$m and $\tau$experiment $= 235.2\,\mu$m.

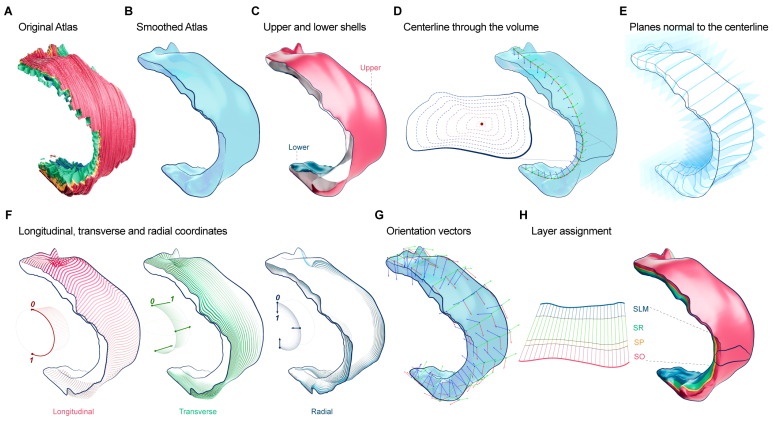


Figure S8: **CA1 atlas overview**. A. Original atlas. B. Smoothed atlas. C. Upper and lower shells. D. Centerline through the volume. E. Planes normal to the centerline. F. Layer assignment. G. Longitudinal, transverse and radial coordinates. H. Orientation vectors.

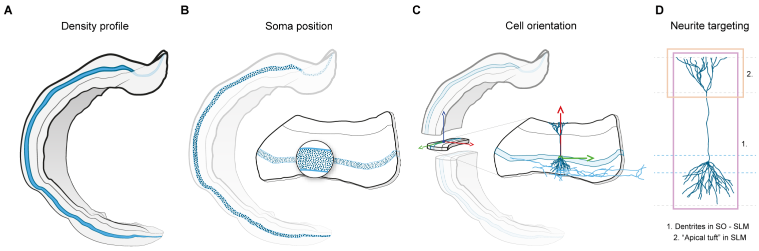


Figure S9: **Cell placement**. A pyramidal cell (PC) is used to illustrate the cell placement in CA1 volume. A. We set a density profile for each cell type. In the case of PC, we set an uniform density in stratum pyramidal (SP). B. We randomly identify soma positions matching the given cell density. C. For each soma position, we assign a morphology and orient it using the vector fields. For PC, we align the main axis of the dendrites with a vector (red arrow) parallel to the radial axis, while we align the axon to a vector (green arrow) parallel to the transverse axis. Furthermore, the more complex branch of the axon points to the Subiculum. D. We select a morphology that respects a set of rules. In the case of PC, we specify two rules regarding the dendrites.

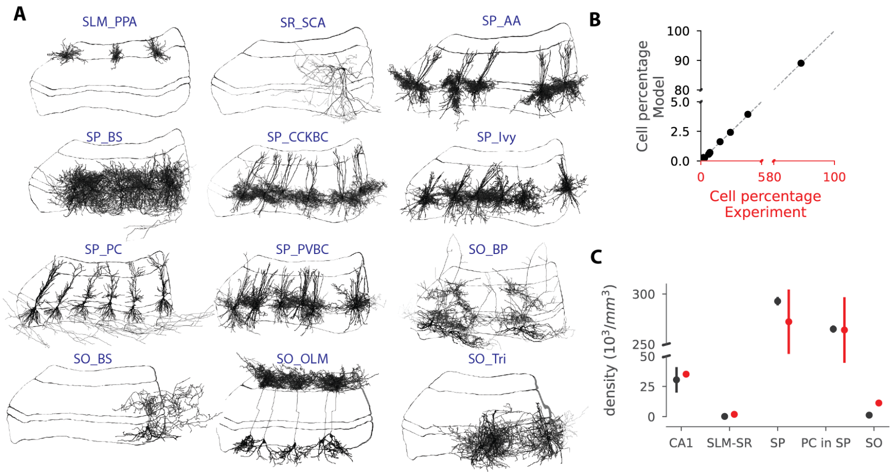


Figure S10: **Validation of cell placement**. A. Validation of the cell placement. A subset of cells from each m-type is displayed within each of the 100 slices of $100\,\mu m$ thickness equally spanned along the longitudinal axis. B. Total cell density in CA1, in the layers SLM + SR, SP, pyramidal cell density in SP, total cell density in SO. Neuron density validation is intrinsic (PC in SP) and extrinsic. The density is sampled in different subvolumes (9 cylinders of $300\,\mu m$ of radius equally spanned along the longitudinal axis). C. Cell composition is sampled in different subvolumes (9 cylinders of $300\,\mu m$ of radius equally spanned along the longitudinal axis) and compared with desired composition (Bezaire & Soltesz, 2013) ($R = 0.999998$, $p < 0.0001$)

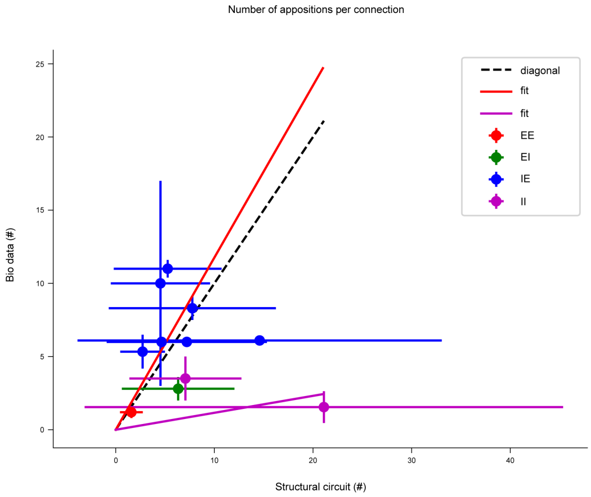


Figure S11: **Prediction of the number of synapses per connection from the number of appositions per connections**. We compare the available data on the number of synapses per connection of a given pathway to the corresponding number of appositions per connection. Data can be grouped in two sets and fit separately (purple line $y = 0.1096x$ for I-I, red line $y = 1.1690x$ for the rest). The fitting lines can be use to predict how much the appositions should be pruned to match or predict synapses per connections. E: excitatory neuron, I: inhibitory neuron.

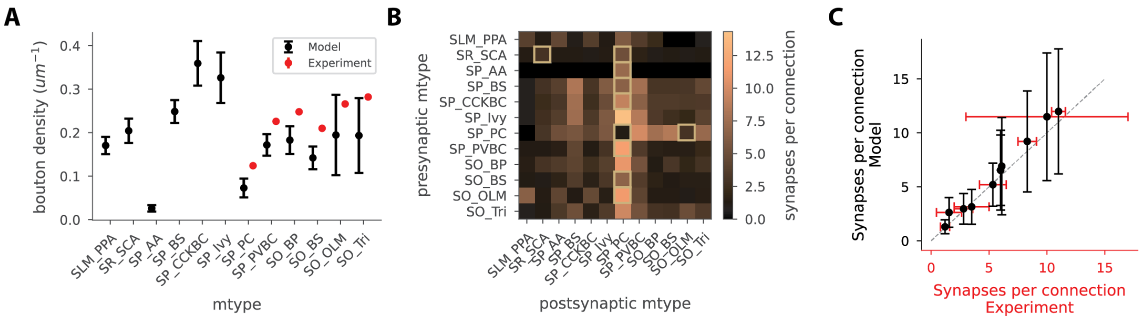


Figure S12: **Bouton density and synapses per connection.** A. Mean and std of bouton density values per individual m-type. B. Mean synapses per connection for each m-type pair. Experimental values are given in square brackets. C. Comparisons of mean and std of synapses per connection for experimental data points and corresponding values in the model.

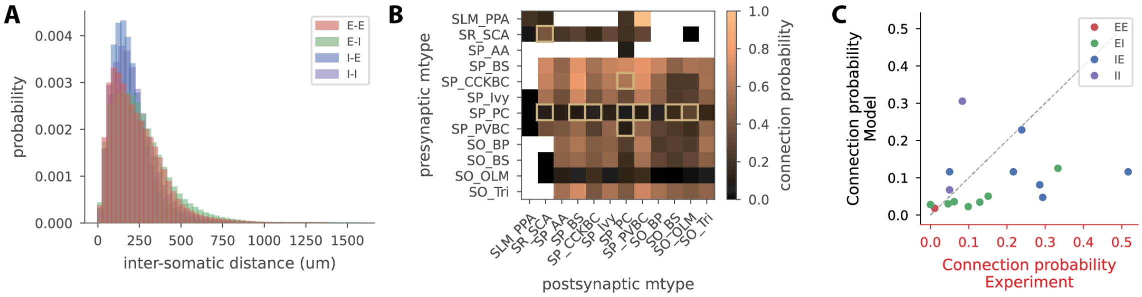


Figure S13: **Connection probability** A. Connection probability with respect to the intersomatic distance within and between Excitatory (E) and Inhibitory (I) groups. B. Mean connection probability for each m-type pair. Non-existing connections are left blank and experimental observations are shown with square brackets. C. Comparison of experimental and model connection probabilities within E/I group pairs.

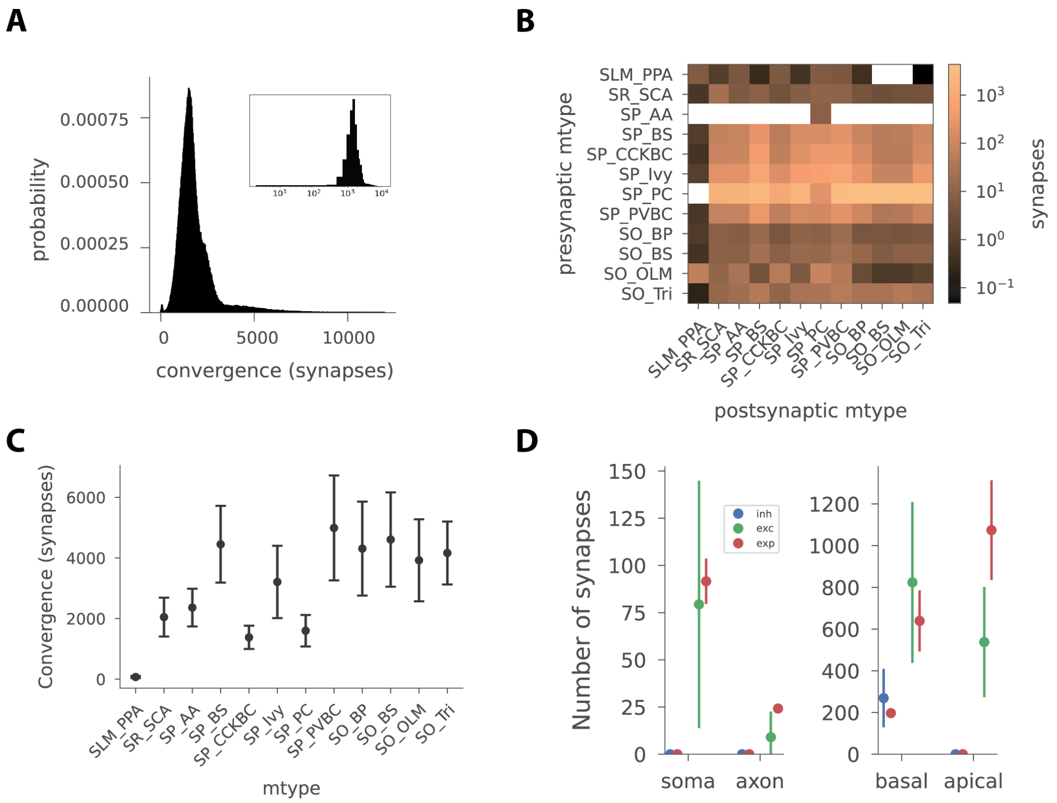


Figure S14: **Convergence on neurons and neuron groups.** A. Indegree distribution for neurons in this model. The inset shows the distribution on logarithmic scale. B. Number of synapses made on an average post-synaptic cell from each afferent m-type group. Colorbar in log-normal scale. C. Mean and std of synapses made onto an average neuron for each m-type. D. Number of synapses made onto each neurite type for excitatory and inhibitory classes.

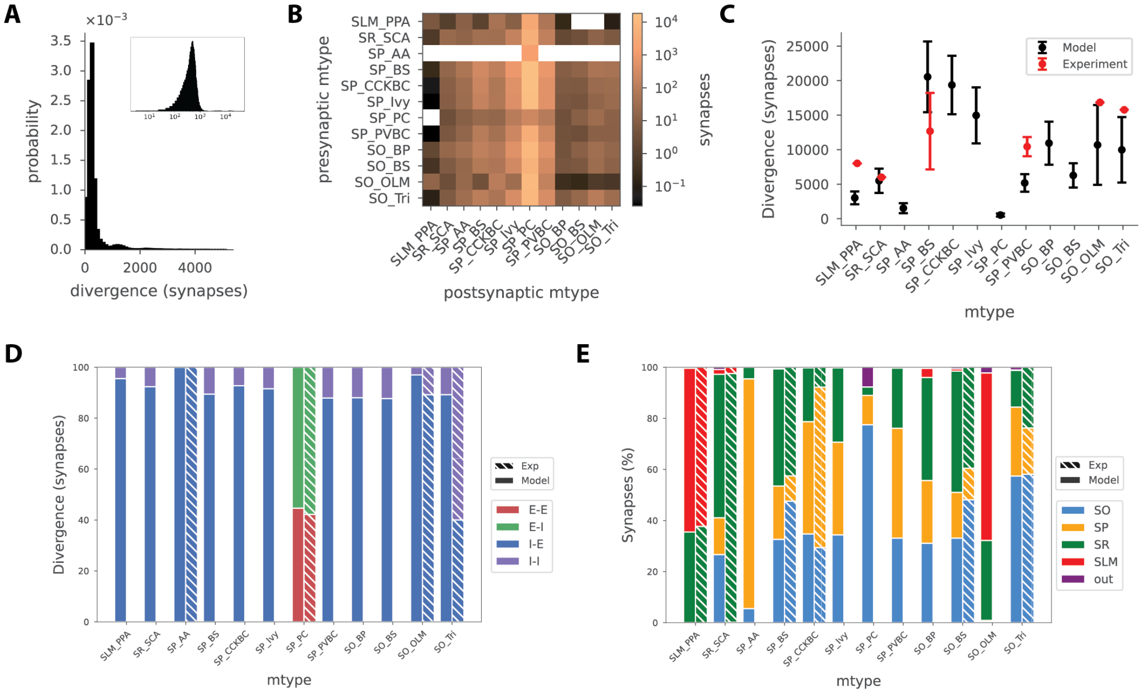


Figure S15: **Divergence of neurons and neuron groups.** A. Outdegree distribution for neurons in this model. The inset shows the distribution on logarithmic scale. B. Number of synapses made by an average neuron from presynaptic m-type to postsynaptic group. C. Comparison of divergence per m-type with the available experimental values. D. Prediction of synapse divergence broken down into efferent synaptic group. E. Percentage of synapses made onto individual layers or outside the CA1 mesh. For D and E, hatched bars indicate the experimental values.

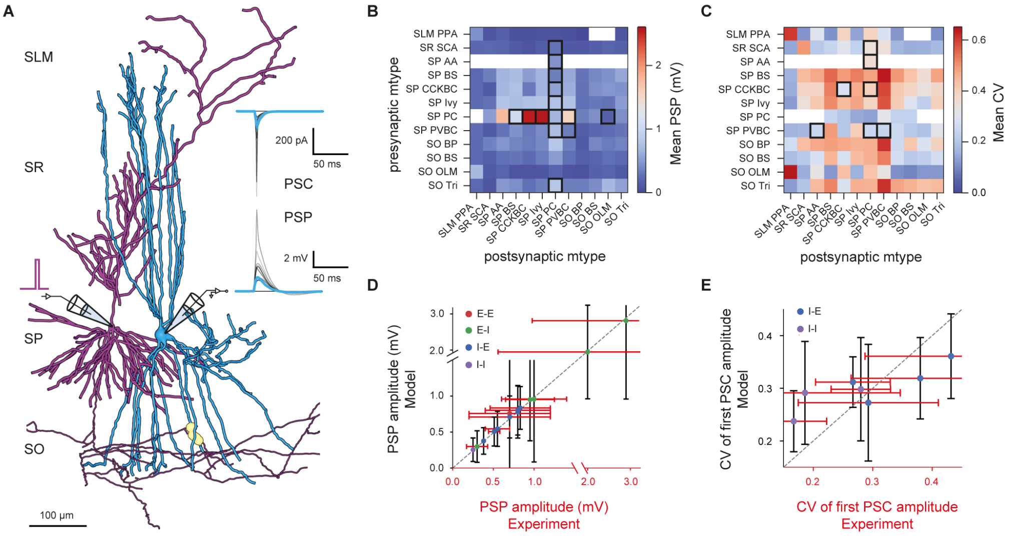


Figure S16: **Prediction and validation of synapses**. A. Example of *in silico* paired recording experiment between a PC (purple morphology) and a PVBC (light blue morphology), synapses between PC axon and PVBC dendrites in SO are depicted as yellow spheres. The presynaptic neuron is stimulated with a step current to elicit an action potential. Somatic recordings of EPSP and EPSC are depicted on the right. Each grey line represents one of the 35 trials, while the average response is shown in light blue. B. Prediction of the PSP amplitudes (B) and PSC CVs (C) for the 130 possible pathways. C. The combinations with bold borders indicate pathways that have been validated (panels C and D). Validation of PSP amplitudes (C) and PSC CVs (D). Dots represent mean values and whiskers the standard deviation of experimental (horizontal, in red) and model (vertical, in black) data. The dashed gray line indicates the diagonal (i.e., the target).

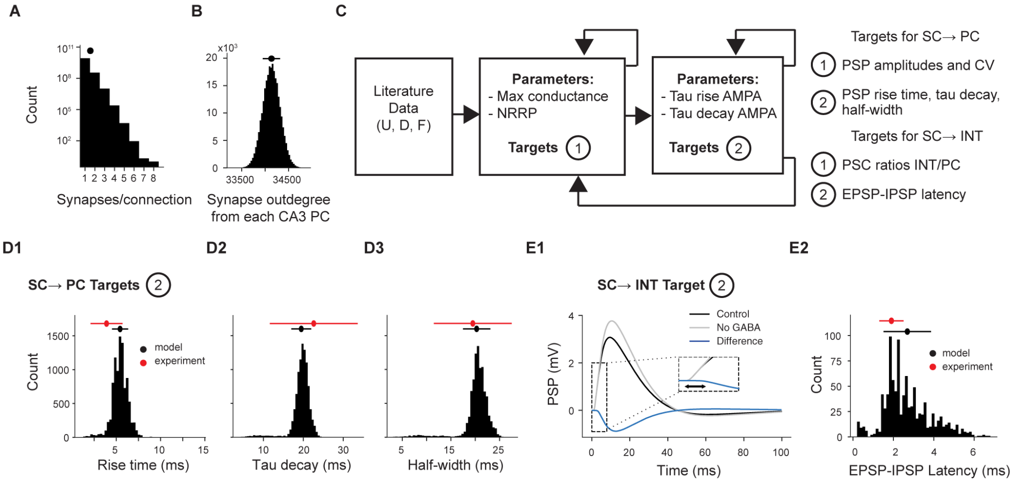


Figure S17: **Schaffer collaterals anatomy and physiology**. A. The distribution of the number of synapses per connection (1.0 ± 0.2 synapses/connection), the y-axis has a logarithmic scale. B. The distribution of efferent synapses made by a single SC (34,135 ± 185 synapses). C. Scheme of the workflow used for the fitting of SC → PC and SC → INT synapses. On the right, targets that are used for the two steps for the fitting of SC → PC and SC → INT. D. Fitting results of SC → PC synapses. Each plot reports the distribution of one PSP feature (rise time, tau decay, and half-width in D1, D2, and D3, respectively) computed over the 10,000 pairs of pre and postsynaptic neurons. On top, experimental (in red) and model (in black) mean and standard deviation values are reported with a dot and a bar, respectively. E. Fitting results of SC → INT synapses. E1 shows the average PC EPSPs in control conditions (black line) and when gabazine is applied (no GABA, gray line). The difference between the two is the IPSP induced by the feedforward inhibition (blue line). The inset shows the EPSP-IPSP latency, which is the difference between the onset of the IPSP and of the EPSP. E2. The EPSP-IPSP latency distribution of the 1000 randomly selected PCs.

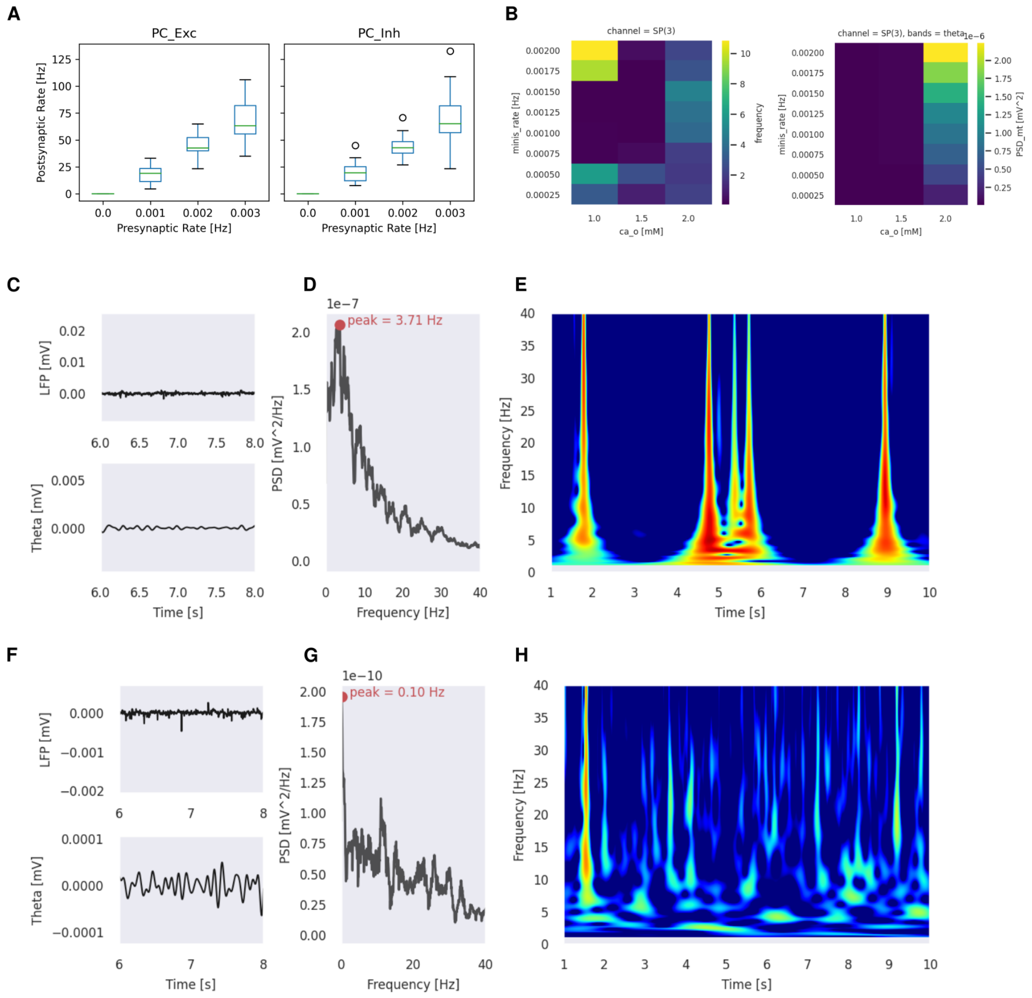


Figure S18: **Spontaneous synaptic release alone did not generate sustained theta oscillations in the CA1 model**. A. Relationship between spontanteous presynaptic release and postsynaptic events rates for pyramidal cells EPSPs (left) and IPSPs (right). B. Relationship between calcium level and spontaneous presynaptic release rate for stratum pyramidale LFP responses peak frequency (left) and theta band power (right) shows weak theta power across all simulation experiments. C-H. Example: 0.001 Hz presynaptic spontaneous release rate (cylinder circuit). C-E. 2 mM calcium. C. LFP and theta-band filtered LFP extracellular recordings from stratum pyramidale show irregular activity. D. Power spectral density (PSD) shows multiple noisy peaks 1-20 Hz with the highest peak just below theta range. E. Morlet complex wavelet spectrogram shows intermittent episodes of theta-band activity but these were associated with wide-range frequency response. F-H. 1 mM calcium. F. LFP and theta-band filtered LFP extracellular recordings show irregular activity but much smaller amplitude than for 2 mM calcium. G. PSD shows multiple noisy peaks across a wider frequency range several orders of magnitude less than for 2 mM although the highest peak is just within theta-band. H. Wavelet spectrogram shows a slightly more sustained period of theta and delta-band (1-3 Hz) activity but with more irregular, higher frequency events than 2 mM calcium.

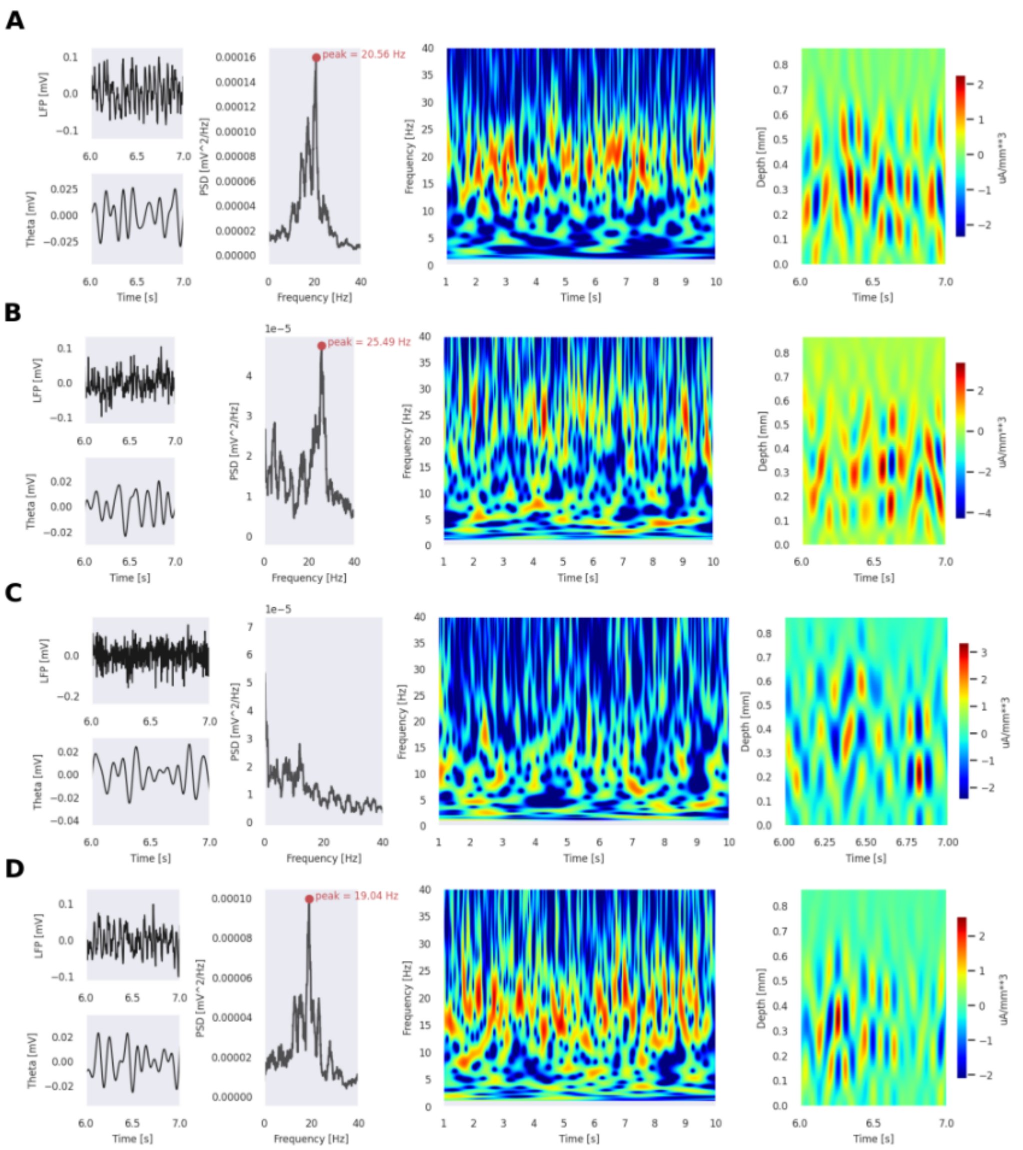


Figure S19: **Extrinsic random synaptic activity at low levels generated noisy beta but not theta oscillatory activity in the CA1 model**. For each panel: LFP and theta filtered LFP traces (far left), PSD (middle left), wavelet spectrogram (middle right), CSD (far right) A. Poisson rate 0.05 Hz. PSD shows broad and noisy peak power located in beta frequency band (13-25 Hz) B. Poisson rate 0.10 Hz. C. Poisson rate 0.60 Hz. PSD shows little power within theta and beta band (peak, not shown, 81.6 Hz). D. Poisson rate 0.05 Hz without pyramidal-pyramidal connectivity. Compared with panel A, absence of recurrent excitation in CA1 shows minor change in peak power and its frequency.

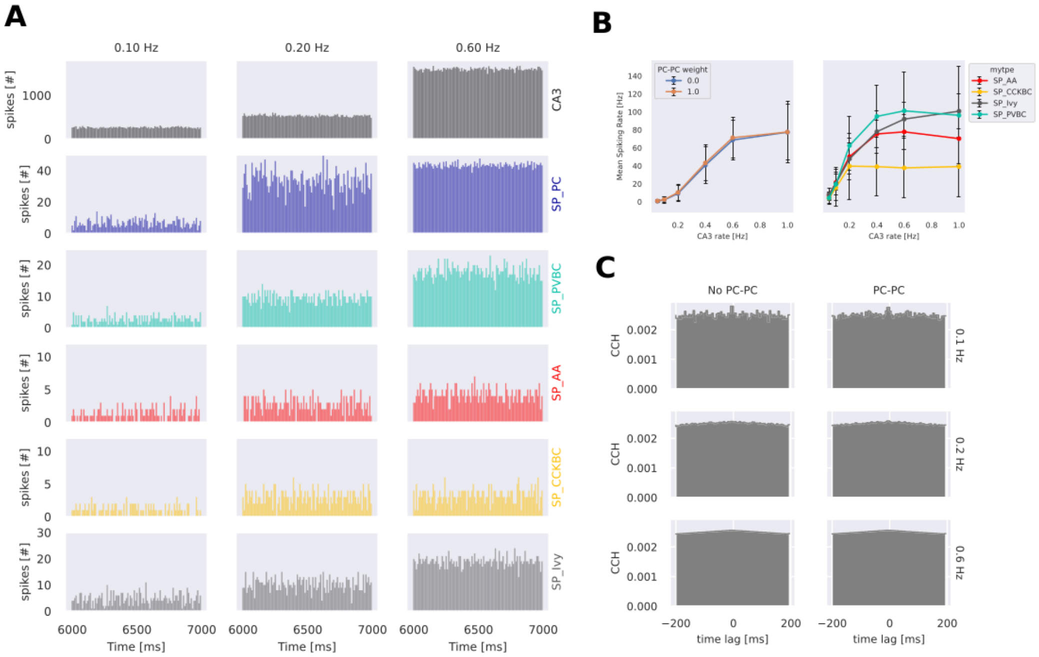


Figure S20: **Extrinsic random synaptic activity fails to induce oscillatory spiking response in the CA1 circuit**. A. Sample histograms of CA3 input spike time input (top) and output post-stimulus response of CA1 pyramidal and interneurons for a range of CA3 Poisson rates. B. Spiking input-output relationship between CA3 input spike rate and mean output spiking rates of pyramidal cells with and without PC-PC interconnections (left) and interneuron types where PC-PC interconnections were present (right). C. Pyramidal cell cross-correlation histograms (CCH) for different CA3 input spike rates both with and without PC-PC interconnections lack evidence of oscillatory response to randomly timed extrinsic afferent EPSPs.

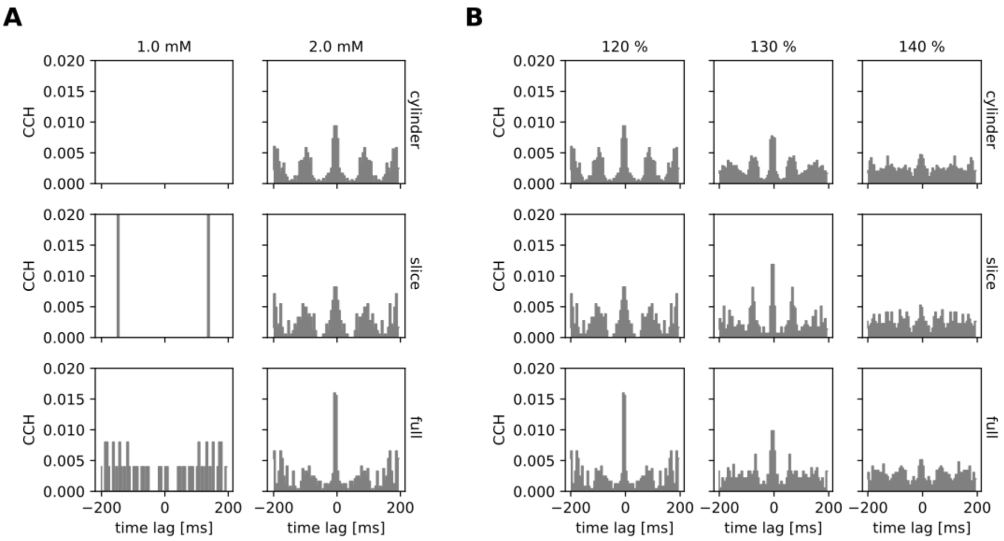


Figure S21: **Tonic depolarisation induces spiking oscillations at** $2\,\mathrm{mM}$ **but not** $1\,\mathrm{mM}$ **calcium for a narrow percentage of relative rheobase across circuit scales.** A. At 120% rheobase, for $1\,\mathrm{mM}$ calcium sparse and uncoordinated spiking but at oscillatory peaks occur at $2\,\mathrm{mM}$ in pyramidal cell cross-correlation histograms (CCH) in all sizes of circuit. B. For $2\,\mathrm{mM}$ calcium, increasing level of tonic depolarisation degrades the strength of oscillation for all sizes of circuit suggesting the resonance effect is limited to a narrow range of tonic depolarisation.

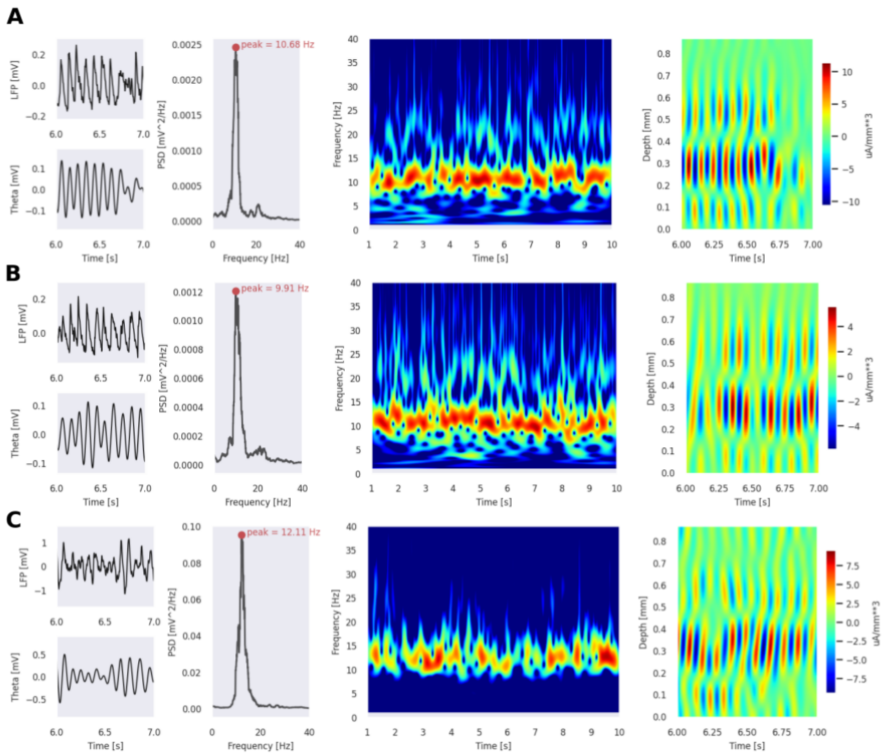


Figure S22: **Tonic depolarisation generates theta-band oscillations across circuit scales at 2 mM extracellular Calcium concentration**.Example: 2 mM Calcium, 120% rheobase depolarisation (recording electrode in SP). For each panel: LFP and theta filtered LFP traces (far left), PSD (middle left), Wavelet Spectrogram (middle right), CSD (far right). A. Cylinder circuit. B. Slice circuit. C. Full circuit.

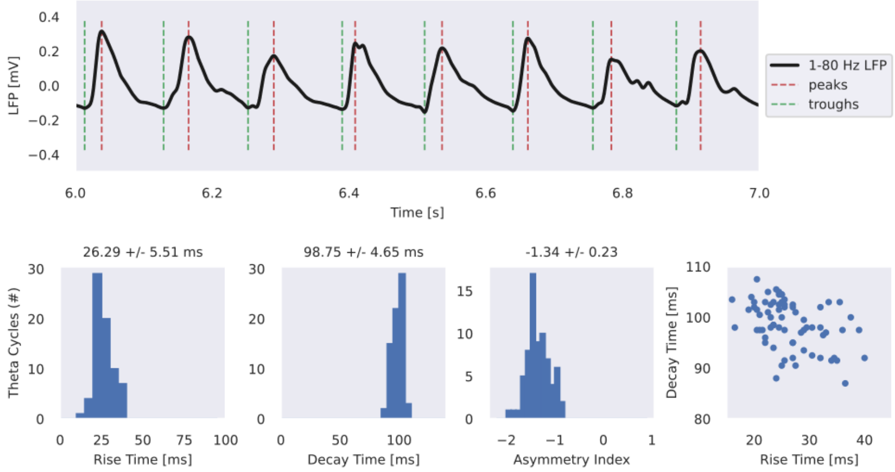


Figure S23: **LFP theta waves are highly asymmetric for an extrinsic excitatory oscillatory stimulus.** Top panel: 1 second example traces of 1-80 Hz filtered LFP from stratum pyramidal SP (3) electrode with estimated locations of theta peaks and troughs shown. Bottom panel: estimated rise times (trough to peak time) (far left), estimated decay times (peak to trough time) (middle left), calculated theta wave asymmetry index (middle right), and scatter plot of rise and decay times (n = 71 total waves). Example: 8 Hz signal frequency, 0.4 Hz cell frequency, 2 mM calcium, full circuit.

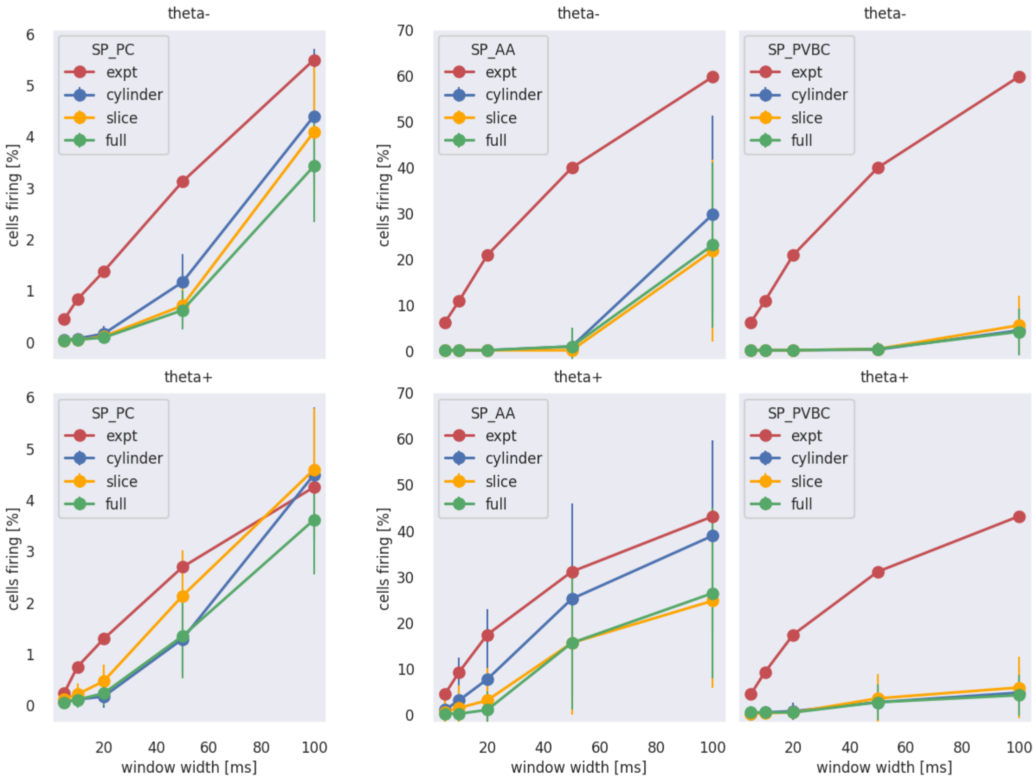


Figure S24: **Population synchrony of pyramidal cells does not match experimental theta trough ('theta-') and fast-spiking interneurons recruitment lower than experimental levels although better for SP_AA than SP_PVBC independent of circuit scale**. Example: 2 mM calcium, 0.4 Hz cell frequency and 8 Hz modulation frequency.

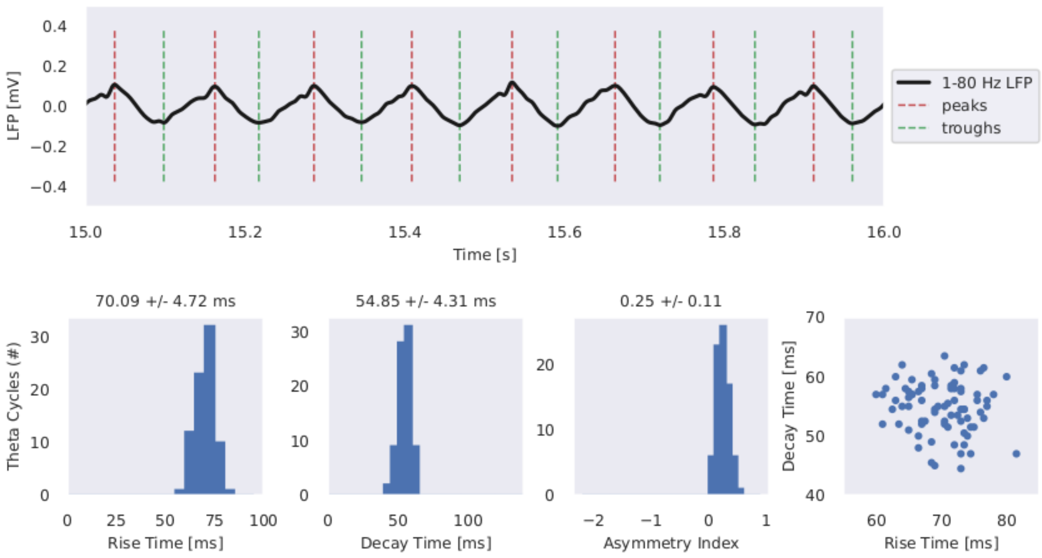


Figure S25: **LFP theta waves are nearly symmetric for an extrinsic inhibitory oscillatory stimulus**.Top panel: 1 second example traces of 1-80 Hz filtered LFP from str pyramidal SP(3) electrode with estimated locations of theta peaks and troughs shown. Bottom panel: estimated rise times (trough to peak time) (far left), estimated decay times (peak to trough time) (middle left), calculated theta wave asymmetry index (middle right), and scatter plot of rise and decay times (n = 79 total waves). Example: 8 Hz signal frequency, 1 μM ACh, depolarisation = 120%, 2 mM Calcium, cylinder circuit.

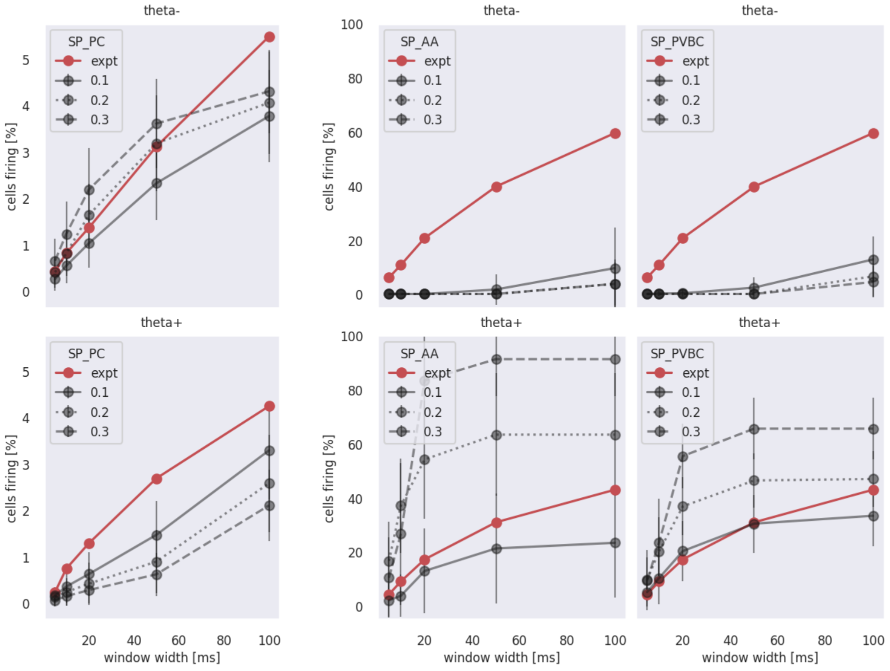


Figure S26: **Population synchrony of pyramidal cells matches experimental theta trough ("theta-") and fast-spiking interneurons SP_AA and SP_PVBC experimental matches theta peak ("theta+") for increased stimulus amplitudes**. Example: 120% depolarisation, 1 μM ACh.

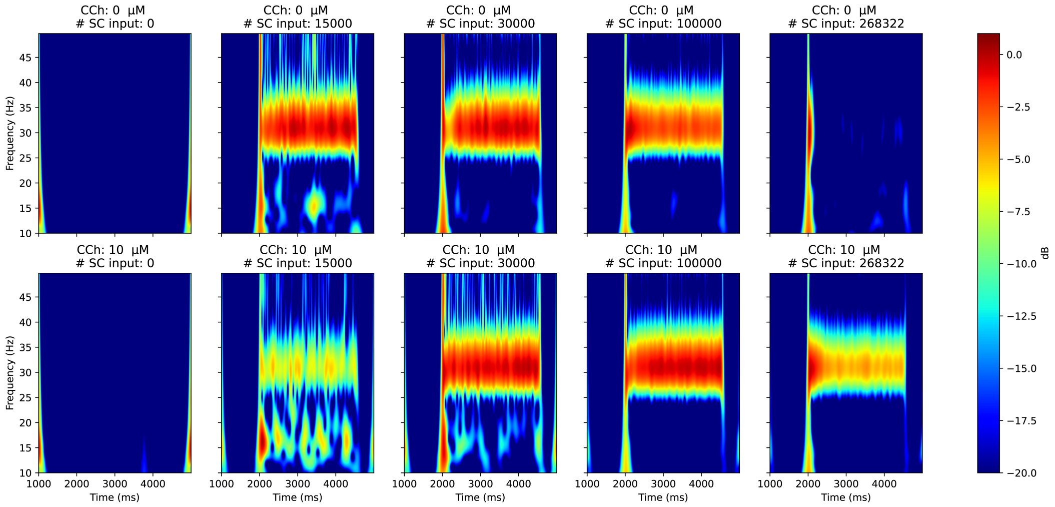


Figure S27: **Propagation of gamma oscillation from CA3 to CA1**. Gamma oscillation (31 Hz) robustly propagates to CA1 when there is a sufficient amount of SC input. (A) shows wavelet spectrograms of LFPs recorded in simulations where CA1 neurons were driven by oscillating input via different numbers of Schaffer Collaterals, but no CCh was applied. (B) shows the wavelet spectrograms of the same simulations, but with simulating the effect of $10\,\mu$M CCh. A wide range of the number of SC inputs is able to induce strong gamma oscillation in CA1 both in the case of CCh application and without CCh, but CCh seems to increase the number of inputs needed for stable gamma oscillation, probably due to its weakening effect on synapses.

#### Supplementary Tables

| Abbreviation | Full name |
|---|---|
| ACh | acetylcholine |
| AMPA | $\alpha$-amino-3-hydroxy-5-methyl-4-isoxazolepropionic acid |
| bAC | bursting accommodating |
| BP | back-projecting m-type neuron |
| BPAP | back-propagating action potential |
| CA | cornu ammonis |
| CA1 | field CA1 |
| CA3 | field CA3 |
| CCh | carbachol |
| cACpyr | classical accommodating for pyramidal cells |
| cAC | classical accommodating for interneurons |
| CB1R | cannabinoid receptor type 1 |
| CCK+ | cholecystokinin-positive |
| cNAC | classical non-accommodating |
| CSD | current source density |
| CV | coefficient of variation |
| CWT | continuous wavelet transform |
| DG | dentate gyrus |
| e-feature | electrophysiological feature |
| e-type | electrical type |
| EPSC | excitatory postsynaptic current |
| EPSP | excitatory postsynaptic potential |
| GABA | gamma-aminobutyric acid or $\gamma$-aminobutyric acid |
| GABA_A *R* | GABA_A receptor |
| HCN | hyperpolarization-activated cyclic nucleotide–gated channel |
| I_depol | depolarizing current |
| I_h | nonspecific hyperpolarization-activated cation current |
| I-O | input-output |
| INT | interneurons |
| IPSP | Inhibitory postsynaptic potential |
| KYNA | kynurenic acid |
| LFP | local field potential |
| m-type | morphological type |
| me-type | morpho-electrical type |
| minis/mPSP | miniature postsynaptic potentials |
| MOOC | massive online open course |
| MS | medial septum |
| MS OFF | medial septum inactivated stimulus condition |
| MS ON | medial septum activated stimulus condition |
| mM | millimolar = $10^{-3}$ |

| | |
|---|---|
| $\mu$M | micromolar = $10^{-6}$ |
| N.B. | nota bene |
| NMDA | N-methyl-d-aspartate |
| N_RRP | number (size) of the readily releasable pool (#vesicles) |
| nS | nanoSiemens |
| OLM | oriens-lacunosum moleculare m-type neuron |
| PC | pyramidal cell |
| PP | perforant path |
| PPA | perforant path-associated cell |
| PSC | postsynaptic current |
| PSD | power spectral density |
| PSP | postsynaptic potential |
| PV+ | parvalbumin-positive |
| PVBC | parvalbumin-positive basket m-type neuron |
| REM | rapid eye movement |
| SC | Schaffer collateral |
| SCA | Schaffer collateral associated neuron |
| SD rat | Sprague Dawley rat |
| SLM | stratum lacunosum moleculare |
| SLM_PPA | perforant path-associated m-type neuron with soma located in SLM |
| SO | stratum oriens |
| SO_BP | back-projecting m-type neuron with soma located in SO |
| SO_BS | bistratified m-type neuron with soma located in SO |
| SO_OLM | oriens-lacunosum moleculare m-type neuron with soma located in SO |
| SO_Tri | trilaminar m-type neuron with soma located in SO |
| SP | stratum pyramidale |
| SP_AA | axoaxonic m-type neuron with soma located in SP |
| SP_BS | bistratified m-type neuron with soma located in SP |
| SP_CCKBC | cholecystokinin-positive basket cell m-type neuron with soma in SP |
| SP_Ivy | ivy m-type neuron with soma located in SP |
| SP_PC | pyramidal m-type neuron with soma located in SP |
| SP_PVBC | parvalbumin-positive basket m-type neuron with soma located in SP |
| SR | stratum radiatum |
| SR_SCA | Schaffer collateral associated m-type neuron with soma located in SR |
| SST | somatostatin |
| STD | stardard deviation |
| STP | short-term plasticity |
| STTC | spike time tiling coefficient |
| Tri | trilaminar m-type neuron |
| U_SE | release probability of neurotransmitters |
| W rat | Wistar rat |

Table S1: **List of abbreviations and acronyms**

| | This paper | Yu et al (2020) | Bezaire et al (2016) | Cutsuridis et al (2010) | Traub et al (2000) | Traub et al (1992) |
|---|---|---|---|---|---|---|
| Regions | CA1 | EC, DG, CA3 | CA1 | CA1 | CA3 | CA3 |
| Circuit scales | full-scale | reduced full-scale | full-scale | microcircuit | microcircuit | microcircuit |
| Atlas | 3D layered atlas | 2D flat map | layered 3D slab | no | no | no |
| Neuron types | pyramidal<br>axoaxonic<br>basket (CCK+)<br>basket (PV+)<br>bistratified<br>ivy<br>OLM<br>trilaminar<br>PPA<br>SCA | granule cells<br>pyramidal<br>basket | pyramidal<br>axoaxonic<br>basket (CCK+)<br>basket (PV+)<br>bistratified<br>ivy<br>neurogliaform<br>OLM<br>SCA | pyramidal<br>axoaxonic<br>basket<br>bistratified<br>OLM | pyramidal<br>axoaxonic<br>basket<br>bistratified<br>OLM | pyramidal<br>interneurons |
| Short-term synaptic plasticity | yes | no | no | no | no | no |
| Neurotransmitters | AMPA<br>NMDA<br>GABA-A | AMPA<br>NMDA<br>GABA-A | AMPA<br>GABA-A<br>GABA-B | AMPA<br>NMDA<br>GABA-A<br>GABA-B | AMPA<br>GABA-A | AMPA<br>GABA-A<br>GABA-B |
| Spontaneous synaptic release (minis) | yes | no | no | no | no | no |
| Neuromodulation | yes | no | no | no | yes | yes |
| LFP | yes | no | yes | no | no | no |

Table S2: **Key feature comparison of realistic large-scale hippocampal network models with multicompartmental HH model neurons.** For reasons of space, this is a non-exhaustive list of features.

| MType | Percentage | N. cells | Density $\mu m^{-3}$ |
|---|---|---|---|
| SLM_PPA | 0.215 | 1008 | 313.652 |
| SR_SCA | 0.176 | 823 | 120.285 |
| SP_PC | 89 | 416842 | 264000 |
| SP_Ivy | 3.870 | 18127 | 11480.167 |
| SP_PVBC | 2.429 | 11378 | 7206.052 |
| SP_BS | 0.738 | 3457 | 2189.180 |
| SP_AA | 0.646 | 3025 | 1915.533 |
| SP_CCKBC | 1.581 | 7407 | 4691.101 |
| SO_OLM | 0.720 | 3374 | 700.382 |
| SO_BP | 0.083 | 391 | 81.142 |
| SO_Tri | 0.308 | 1440 | 298.944 |
| SO_BS | 0.233 | 1090 | 226.343 |
| Total | 100 | 468362 | |

Table S3: **Cell composition, counts and densities.**

[1]SD rat: Sprague Dawley rat, W rat: Wistar rat, LE rat: Long–Evans rat, G pig: Guinea pig.

| M-type | cNAC | cAC | bAC |
|---|---|---|---|
| SLM_PPA | 0.00% | 0.00% | 100.00% |
| SR_SCA | 0.00% | 100.00% | 0.00% |
| SP_AA | 0.00% | 0.00% | 100.00% |
| SP_BS | 75.00% | 0.00% | 25.00% |
| SP_CCKBC | 0.00% | 100.00% | 0.00% |
| SP_Ivy | 33.33% | 0.00% | 66.67% |
| SP_PC | 0.00% | 100.00% | 0.00% |
| SP_PVBC | 70.00% | 0.00% | 30.00% |
| SO_BP | 0.00% | 100.00% | 0.00% |
| SO_BS | 75.00% | 0.00% | 25.00% |
| SO_OLM | 0.00% | 100.00% | 0.00% |
| SO_Tri | 0.00% | 100.00% | 0.00% |

Table S4: **Morpho-electrical composition.**

| Cell type | Region | Species[1] | Age | Mean ($10^3/mm^3$) | N. animals | STD | SEM | Reference |
|---|---|---|---|---|---|---|---|---|
| All | CA1 | W rat | 9-10 w | 35.2 | 5 | 1.1 | 0.5 | Aika et al. (1994) |
| SO | SO | W rat | 9-10 w | 11.3 | 5 | 2.0 | 0.9 | Aika et al. (1994) |
| SP | SP | W rat | 9-10 w | 272.4 | 5 | 32.0 | 14.3 | Aika et al. (1994) |
| SR-SLM | SR+SLM | W rat | 9-10 w | 1.9 | 5 | 0.7 | 0.3 | Aika et al. (1994) |
| SP_PC | SP | W rat | 9-10 w | 264 | 5 | 32.6 | 14.6 | Aika et al. (1994) |

Table S5: Neuron density validation.

| Mtype | Dendrites | Tuft | Axon |
|---|---|---|---|
| SLM_PPA | SLM | | 2/3 SR-SLM |
| SR_SCA | SO-SLM | | SO-SLM |
| SP_PC | SO-SLM | SLM | |
| SP_Ivy | SO-SR | | 2/3 SO-1/3 SR |
| SP_BS | SO-SR | | SO-SR |
| SP_*BC | SO-SLM | SLM | 2/3 SO-1/4 SR |
| SP_AA | SO-SLM | SLM | 1/2 SO-SP |
| SO_Tri | SO-SR | | SO-SR |
| SO_OLM | | | SLM |
| SO_BS | SO | | SO-SR |
| SO_BP | SO-SLM | | SO-SLM |

Table S6: Placement "optional" rules.

| Mtype | Region | Species[1] | Weight | Mean ($100\mu m^{-1}$) | N. cells | STD | SEM | Reference |
|---|---|---|---|---|---|---|---|---|
| SO_BS | CA1 | SD rat | 250-350 g | 21.0 | 1[2] | 5.6[3] | 5.6 | Sik et al. (1995) |
| SO_BP | CA1 | SD rat | 250-350 g | 24.8 | 1 | 4.13 | 4.13 | Sik et al. (1993)[4] |
| SP_PC | CA1 | W rat | 180-200 g | 12.41 | 4 | 6.02 | 3.01 | Esclapez et al. (1999b) Bezaire and Soltesz (2013)[5] |
| SO_Tri | CA1 | SD rat | 250-350 g | 28.2 | 1 | 4.9[3] | 4.9 | Sik et al. (1995) |
| SP_PVBC | CA1 | SD rat | 250-350 g | 22.6 | 4 | 3.9[3] | 1.95 | Sik et al. (1995) |
| SO_OLM | CA1 | SD rat | 250-350 g | 26.6 | 2 | 4.0[3] | 2.83 | Sik et al. (1995) |

Table S7: Bouton density.

[2]The authors define the sample size (n) probably as the number of sampled segments rather than the number of animals

[3]the authors do not specify if this is std or SEM. Anyway, in a previous publication (Sik et al., 1993) they used std. We can assume they are std

[4]the authors do not mention species and age in the paper. Anyway, a later paper (Sik et al., 1995) mentions the result so we assume they use the same method

[5]Bouton density for PC was computed as a weighted mean of the bouton density per branch order (Esclapez et al., 1999b). Then we analyzed the axon, calculated the total length of each branch order, and used this to weight the bouton density.

| Pre | Post | Specie | Age | Weight | Slice thickness(um) | Distance | n | N | p | Reference |
|---|---|---|---|---|---|---|---|---|---|---|
| PC | PC | SD rat | - | 100-180 g | 400-500 | - | 11 | 989 | 0.011 | Deuchars and Thomson, 1996 |
| PC | OLM | SD rat | - | 90-150 g | 450-500 | - | 12 | 36 | 0.333 | Ali and Thomson, 1998 |
| PVBC | PC | SD rat | - | 120-200 g | 450 | - | 49 | 167 | 0.293 | Pawelzik et al., 2002 |
| PC | PVBC | SD rat | - | 120-200 g | 450 | - | 16 | 124 | 0.129 | Pawelzik et al., 2002 |
| BS | PC | SD rat | - | 120-200 g | 450 | - | 2 | 6 | 0.333 | Pawelzik et al., 2002 |
| PC | BS | SD rat | - | 120-200 g | 450 | - | 2 | 4 | 0.500 | Pawelzik et al., 2002 |
| CCKBC | PC | SD rat | - | 120-200 g | 450 | - | 21 | 88 | 0.239 | Pawelzik et al., 2002 |
| PC | CCKBC | SD rat | - | 120-200 g | 450 | - | 8 | 81 | 0.099 | Pawelzik et al., 2002 |
| CCKBS | PC | SD rat | - | 120-200 g | 450 | - | 5 | 36 | 0.139 | Pawelzik et al., 2002 |
| PC | CCKBS | SD rat | - | 120-200 g | 450 | - | 6 | 35 | 0.171 | Pawelzik et al., 2002 |
| PC | SCA | SD rat | - | 120-200 g | 450 | - | 0 | 32 | 0 | Pawelzik et al., 2002 |
| Ivy | PC | W rat | - | 140-200 g | 450 | - | 3 | 5 | 0.600 | Fuentealba et al., 2008 |
| Ivy | Ivy | W rat | - | 140-200 g | 450 | - | 1 | 4 | 0.250 | Fuentealba et al., 2008 |
| INT | PC | W rat | - | 140-200 g | 450 | - | 6 | 21 | 0.286 | Fuentealba et al., 2008 |
| BC | PC | SD rat | - | 120-200 g | 450-500 | - | 57 | 263 | 0.217 | Ali et al., 1999 |
| BC | PC | SD rat | - | 120-200 g | - | 50-100 | 46 | 89 | 0.517 | Ali et al., 1999 |
| BC | PC | SD rat | - | 120-200 g | - | >150-200 | 6 | 120 | 0.050 | Ali et al., 1999 |
| PC | BS | SD rat | - | 90-180 g | 450-500 | - | 8 | 53 | 0.151 | Ali et al., 1998 |
| PC | BC | SD rat | - | 90-180 g | 450-500 | - | 9 | 195 | 0.046 | Ali et al., 1998 |
| PC | SP_INT | SD rat | - | 90-180 g | 450-500 | - | 23 | 371 | 0.062 | Ali et al., 1998 |
| SCA | SCA | W rat | 18-23 d | - | 300-330 | - | 20 | 240 | 0.083 | Ali, 2007 |
| CCKBC | PC | SD rat | 16-20 d | - | 350 | - | | | <0.100 | Neu et al., 2007 |
| SO | SLM | SD rat | 18-22 d | - | 300-350 | - | 1 | 20 | 0.050 | Elfant et al., 2008 |

Table S8: Connection probabilities per m-type pair and other related parameters.

| From | To | Region | Species[1] | Age | Weight | mean | n. conns | STD | SEM | Reference |
|---|---|---|---|---|---|---|---|---|---|---|
| SP_BS | SP_PC | CA1 | - | - | - | 6 | 1 | - | - | Buhl, Halasy, and Somogyi (1994) |
| SP_PVBC | PV | CA1 | SD rat | - | 250-350 g | 1.55 | 64 | 1.08 | 0.14 | Sik et al. (1995) |
| SP_PC | SO_OLM | CA1 | W rat | 14-21 d | - | 2.83 | 6 | 1.94 | 0.79 | Biro (2005) |
| SO_OLM | SP_PC | CA1 | W rat | 10-17 d | - | 10 | 2 | 9.90 | 7 | Maccaferri et al. (2000) |
| AA | GC | CA3 | W rat | Young | - | 8 | 1 | - | - | Buhl, Han, et al. (1994) |
| AA | SP_PC | CA1 | W rat | Young | - | 5.89 | 9 | - | - | Buhl, Han, et al. (1994) |
| SP_PC | SP_PC | CA1 | SD rat | - | 100 -180 g | 1.17 | 6 | 0.41 | 0.17 | Deuchars and Thomson (1996) |
| SP_CCKBC | SP_PC | CA1 | SD rat<br>Mouse | 14-21 d<br>> 21 d | - | 8.3 | 14 | 0.8 | 0.21 | Földy et al. (2010) |
| SR_SCA | SP_PC | CA1 | W rat | - | > 120 g | 5.33 | 3 | 1.15 | 0.67 | Vida et al. (1998) |
| SP_PVBC | SP_PC | CA1 | SD rat<br>Mouse | 14-21 d<br>> 21 d | - | 11 | 15 | 0.6 | 0.15 | Földy et al. (2010) |
| SR_SCA | SR_SCA | CA1 | W rat | 18-21 d | - | 3.5 | 9 | 1.5 | 0.5 | Ali (2011) |

Table S9: Number of synapses per connection.

| M-type | Region | Specie | Age | Weight | mean | n. cells | std | SEM | Reference |
|---|---|---|---|---|---|---|---|---|---|
| SP_PVBC | CA1 | SD rat | - | 250-350 g | 10436 | 4 | 1393.26 | 696.63 | Sik et al., 1995 |
| SP_BC | CA1 | W rat | 7-8 w | - | 10828 | 1 | 0.00 | 0.00 | Halasy et al., 1996 |
| SR_SCA | CA1 | W rat | - | >120 g | 5998 | 1 | 0.00 | 0.00 | Vida et al., 1998 |
| SR_CCKBC | CA1 | W rat | - | >120 g | 7964 | 1 | 0.00 | 0.00 | Vida et al., 1998 |
| SP_BS | CA1 | SD / W rat | 7-8 w | 250-350 g | 12676 | 2 | 5549.37 | 3924.00 | Sik et al., 1995; Halasy et al., 1996 |
| SO_OLM | CA1 | SD rat | - | 250-350 g | 16847 | 1 | 0.00 | 0.00 | Sik et al., 1995 |
| SLM_PPA | CA1 | W rat | - | >120 g | 8015 | 1 | 0.00 | 0.00 | Vida et al., 1998 |
| SO_Tri | CA1 | SD rat | - | 250-350 g | 15767 | 1 | 0.00 | 0.00 | Sik et al., 1995 |

Table S10: Experimentally available data for divergence of synapses per m-type.

| M-type | Specie | Age | Weight | PC | INT | n | Reference |
|---|---|---|---|---|---|---|---|
| PC | SD rat | - | 250-350 g | 42.151 | 57.849 | 130 | Takacs et al., 2012 |
| OLM | W rat | 2 m | - | 89.157 | 10.843 | 34 | Katona et al., 1999 |
| Tri | W rat | - | 300-400 g | 40 | 60 | 52 | Ferraguti et al., 2005 |
| AA | - | - | - | 100 | 0 | - | AA.VV. |

Table S11: Available data on the divergence of synapses for different m-types to excitatory and inhibitory groups within CA1.

| | model SO | model SP | model SR | model SLM | model out | exp SO | exp SP | exp SR | exp SLM |
|---|---|---|---|---|---|---|---|---|---|
| SLM_PPA | 0.000 | 0.000 | 42.438 | 57.306 | 0.256 | 0.000 | 0.000 | 37.700 | 62.300 |
| SO_BP | 27.048 | 28.990 | 40.491 | 3.087 | 0.385 | | | | |
| SO_BS | 29.801 | 17.232 | 50.586 | 1.981 | 0.400 | 48.100 | 12.400 | 39.500 | 0.000 |
| SO_OLM | 0.358 | 0.495 | 40.895 | 55.855 | 2.396 | | | | |
| SO_Tri | 56.670 | 26.564 | 15.547 | 0.204 | 1.015 | 58.120 | 18.110 | 23.770 | 0.000 |
| SP_AA | 5.949 | 89.659 | 4.392 | 0.000 | 0.000 | | | | |
| SP_BS | 33.775 | 19.989 | 45.875 | 0.152 | 0.210 | 47.600 | 9.850 | 42.550 | 0.000 |
| SP_CCKBC | 32.540 | 45.842 | 21.602 | 0.009 | 0.008 | 29.400 | 62.850 | 7.750 | 0.000 |
| SP_Ivy | 32.293 | 33.412 | 34.265 | 0.019 | 0.011 | | | | |
| SP_PC | 74.662 | 12.693 | 3.580 | 0.006 | 9.059 | | | | |
| SP_PVBC | 30.928 | 44.182 | 24.830 | 0.046 | 0.015 | | | | |
| SR_SCA | 29.081 | 13.885 | 54.179 | 2.410 | 0.444 | 0.000 | 0.200 | 97.400 | 2.400 |
| SR_CCKBC | | | | | | 9.100 | 57.700 | 31.750 | 1.450 |
| SR_Tri | | | | | | 27.000 | 23.000 | 50.000 | 0.000 |

Table S12: Laminar distribution

| Rule | From | To | Rule type[6] | U (mean ± STD) | D (ms) (mean ± STD) | F (ms) (mean ± STD) | N_RRP | Hill scaling | CV Validation pathway used |
|---|---|---|---|---|---|---|---|---|---|
| 1 | SP_PC | SP_PC | E2 | 0.65±0.1 | 671±17 | 17±5 | 2 | 2.79 | - |
| 2 | SP_PC | SO_OLM | E1 | 0.09±0.12 | 138±211 | 670±830 | 1 | 2.79 | - |
| 3 | SP_PC | SO_Tri<br>SO_BS<br>SO_BP | E1 | 0.09±0.12 | 138±211 | 670±830 | 1 | 1.94 | - |
| 4 | SP_PC | SP_AA | E2 | 0.23±0.09 | 410±190 | 10±11 | 1 | 1.09 | - |
| 5 | SP_PC | SP_BS | E2 | 0.23±0.09 | 410±190 | 10±11 | 1 | 1.94 | - |
| 6 | SP_PC | SP_CCKBC | E2 | 0.23±0.09 | 410±190 | 10±11 | 1 | 1.09 | - |
| 7 | SP_PC | SP_Ivy | E2 | 0.5±0.02 | 671±17 | 17±5 | 1 | 1.94 | - |
| 8 | SP_PC | SP_PVBC | E2 | 0.23±0.09 | 410±190 | 10±11 | 1 | 1.09 | - |
| 9 | SP_PC | SR_SCA<br>SLM_PPA | E2 | 0.23±0.09 | 410±190 | 10±11 | 1 | 1.94 | - |
| 10 | INH | INH | I2 | 0.26±0.05 | 930±360 | 1.6±0.6 | 1 | 1.94 | - |
| 11 | SP_AA | SP_PC | I2 | 0.1±0.01 | 1278±760 | 10±6.7 | 1 | 1.94 | SP_AA→SP_PC |
| 12 | SP_BS | SP_PC | I2 | 0.13±0.03 | 1122±156 | 9.3±0.7 | 1 | 1.94 | - |
| 13 | SP_PVBC | SP_PC | I2 | 0.16±0.02 | 965±185 | 8.6±4.3 | 9 | 1.94 | SP_PVBC→SP_PC |
| 14 | SO_OLM<br>SO_BS<br>SO_BP | SP_PC | I2 | 0.3±0.08 | 1250±520 | 2±14 | 1 | 1.94 | - |
| 15 | SO_Tri | SP_PC | I2 | 0.3±0.08 | 1250±520 | 2±14 | 1 | 1.94 | - |
| 16 | SLM_PPA | SP_PC | I3 | 0.16±0.01 | 168±15 | 13±0.5 | 1 | 1.94 | - |
| 17 | SP_CCKBC | SP_PC | I3 | 0.16±0.04 | 153±120 | 12±3.5 | 1 | 1.94 | SP_CCKBC→SP_PC |
| 18 | SR_SCA | SP_PC | I3 | 0.15±0.03 | 185±32 | 14±5.8 | 1 | 1.94 | SR_SCA→SP_PC |
| 19 | SP_Ivy | SP_PC | I3 | 0.32±0.14 | 144±80 | 62±31 | 1 | 1.94 | - |

| | | | | | | | | | |
|---|---|---|---|---|---|---|---|---|---|
| 20 | SP_PVBC | SP_AA | I2 | 0.24±0.15 | 1730±530 | 3.5±1.5 | 1 | 1.94 | SP_PVBC→SP_AA |
| | SP_CCKBC | SP_CCKBC | | | | | | | |
| 21 | SR_SCA | SR_SCA | I1 | 0.11±0.03 | 115±100 | 1542±700 | 3 | 1.94 | SP_CCKBC→SP_CCKBC |
| | SLM_PPA | SLM_PPA | | | | | | | |
| 22 | SP_PVBC | SP_PVBC | I2 | 0.26±0.05 | 930±360 | 1.6±0.6 | 9 | 1.94 | SP_PVBC→SP_PVBC |

Table S13: Presynaptic dynamics parameters.

[6]Rule types: E1: excitatory facilitating, E2: excitatory depressing, I1: inhibitory facilitating, I2: inhibitory depressing, I3: inhibitory pseudo linear.

| Rule | From | To | Rule type[6] | gsyn (nS) (mean ± STD) | $\tau_{decay}$ fast (ms) (mean ± STD) | NMDA/AMPA ratio (mean ± STD) | $\tau_{decay}$ NMDA (ms) (mean ± STD) | PSP validation pathway used |
|---|---|---|---|---|---|---|---|---|
| 1 | SP_PC | SP_PC | E2 | 0.65±0.1 | 3±0.2 | 1.22 | 148.5 | SP_PC→SP_PC |
| 2 | SP_PC | SO_OLM | E1 | 1.0±0.05 | 1.7±0.14 | 0.28 | 148.5 | SP_PC→SO_OLM |
| 3 | SP_PC | SO_Tri<br>SO BS<br>SO BP | E1 | 1.0±0.05 | 1.7±0.14 | 0.28 | 148.5 | SP_PC→SO_OLM |
| 4 | SP_PC | SP_AA | E2 | 2.93±0.7 | 4.12±0.5 | 0.28 | 148.5 | MEAN RULES 5-8 |
| 5 | SP_PC | SP_BS | E2 | 1.88±0.1 | 4.12±0.5 | 0.28 | 148.5 | SP_PC→SP_BS |
| 6 | SP_PC | SP_CCKBC | E2 | 3.63±0.4 | 4.12±0.5 | 0.86 | 298.75 | SP_PC→SP_CCKBC |
| 7 | SP_PC | SP_Ivy | E2 | 3.64±0.4 | 4.12±0.5 | 0.28 | 148.5 | SP_PC→SP_Ivy |
| 8 | SP_PC | SP_PVBC | E2 | 2.56±0.05 | 4.12±0.4 | 0.28 | 148.5 | SP_PC→SP_PVBC |
| 9 | SP_PC | SR_SCA<br>SLM_PPA | E2 | 2.93±0.7 | 4.12±0.5 | 0.86 | 298.75 | MEAN RULES 5-8 |
| 10 | INH | INH | I2 | 3.74±0.3 | 4±0.8 | - | - | SP_PVBC→SP_PVBC |
| 11 | SP_AA | SP_PC | I2 | 2.01±0.1 | 11.2±0.9 | - | - | SP_AA→SP_PC |
| 12 | SP_BS | SP_PC | I2 | 1.92±0.1 | 16.1±1.1 | - | - | SP_BS→SP_PC |
| 13 | SP_PVBC | SP_PC | I2 | 1.87±0.2 | 5.94±0.47 | - | - | SP_PVBC→SP_PC |
| 14 | SO_OLM<br>SO BS<br>SO BP | SP_PC | I2 | 1.61±0.3 | 8.3±2.2 | - | - | SO_Tri→SP_PC |
| 15 | SO_Tri | SP_PC | I2 | 1.61±0.3 | 7.75±0.9 | - | - | SO_Tri→SP_PC |
| 16 | SLM_PPA | SP_PC | I3 | 2.2±0.15 | 8.8±0.25 | - | - | MEAN RULES 17-18 |
| 17 | SP_CCKBC | SP_PC | I3 | 1.7±0.3 | 9.35±1.0 | - | - | SP_CCKBC→SP_PC |
| 18 | SR_SCA | SP_PC | I3 | 2.69±0.3 | 8.3±0.44 | - | - | SR_SCA→SP_PC |

| | | | | | | | | |
|---|---|---|---|---|---|---|---|---|
| 19 | SP_Ivy | SP_PC | I3 | 0.65±0.05 | 16±2.5 | - | - | SP_Ivy→SP_PC |
| 20 | SP_PVBC<br>SP_CCKBC | SP_AA<br>SP_CCKBC | I2 | 3.74±0.3 | 2.67±0.13 | - | - | SP_PVBC→SP_PVBC |
| 21 | SR_SCA<br>SLM_PPA | SR_SCA<br>SLM_PPA | I1 | 3.74±0.3 | 4.5±0.55 | - | - | SP_PVBC→SP_PVBC |
| 22 | SP_PVBC | SP_PVBC | I2 | 3.74±0.3 | 2.67±0.13 | - | - | SP_PVBC→SP_PVBC |

Table S14: Postsynaptic dynamics parameters.

| From | To | Experimental Feature | Min | Max | Mean | SD | Species[1] | Weight | Region | N. | n. | Reference |
|---|---|---|---|---|---|---|---|---|---|---|---|---|
| SC | EXC | Number of afferent synapses | 13059 | 28697 | 20878 | - | W rat | 300 g | CA1 | 7 | - | Bezaire and Soltesz (2013) |
| SC | INH | Number of afferent synapses | 7952 | 17476 | 12714 | - | W rat | >110 g | CA1 | 4 | 70 | Bezaire and Soltesz (2013) |
| SC | PV+ INTs | Number of synapses per connection | 1 | 3 | 1.19 | 0.48 | SD rat | 200-300 g | CA3 | - | 274 | Sik et al. (1993) |
| SC | All | Number of efferent synapses | 15295 | 27440 | 21368 | - | SD rat | 200-300 g | CA1 | 6 | - | Bezaire and Soltesz (2013) |

Table S15: Schaffer collaterals anatomy experimental data. N.: number of animals, n.: number of synapses.

| From | To | Average synapse number | Percentage of afferent synapses | Species[1] | Weight | Region | N. | Reference |
|---|---|---|---|---|---|---|---|---|
| SC | SLM | 55 | 0.3% | SD rat | 200-300 g | CA1 | 6 | Bezaire and Soltesz (2013) |
| SC | SR | 14515 | 67.9% | SD rat | 200-300 g | CA1 | 6 | Bezaire and Soltesz (2013) |
| SC | SP | 1507 | 7.1% | SD rat | 200-300 g | CA1 | 6 | Bezaire and Soltesz (2013) |
| SC | SO | 5291 | 24.7% | SD rat | 200-300 g | CA1 | 6 | Bezaire and Soltesz (2013) |

Table S16: Schaffer collaterals layer profile. N.: number of animals. Averages are taken from Table 20 of Bezaire and Soltesz (2013)

| SC→Exc | Experimental Feature | Value (Mean) | SD | SEM | UoM | Species[1] | Age | Weight | R. | n. | Reference |
|---|---|---|---|---|---|---|---|---|---|---|---|
| PSP Magnitude | PSPs amplitudes | 0.14 | 0.106 | 0.01 | mV | G Pig | - | 600-900 g | CA1 | 72 | Sayer et al. (1990) |
| | PSPs CV | 0.76 | - | - | - | G Pig | - | 600-900 g | CA1 | 72 | Sayer et al. (1990) |
| PSP Kinetics | PSPs rise time | 3.9 | 1.8 | 0.21 | ms | G Pig | - | 600-900 g | CA1 | 72 | Sayer et al. (1990) |
| | PSPs half-width | 19.5 | 8 | 0.94 | ms | G Pig | - | 600-900 g | CA1 | 72 | Sayer et al. (1990) |
| | PSP tau decay | 22.6 | 11 | 1.3 | ms | G Pig | - | 600-900 g | CA1 | 72 | Sayer et al. (1990) |
| NMDA Kinetics | NMDA/AMPA ratio | 1.23 | 0.03 | 0.01 | - | Mouse | 17-23 d | - | CA1 | 12 | Le Roux et al. (2013) |
| | NMDA tau rise | 2.93 | - | - | ms | SD rat | 6-12 w | - | CA1 | 52 | Andrásfalvy and Magee (2001) |
| | NMDA tau decay | 148.5 | - | - | ms | SD rat | 6-12 w | - | CA1 | 52 | Andrásfalvy and Magee (2001) |
| Short term plasticity | U | 0.14 | 0.08 | 0.03 | - | W rat | 14-28 d | - | CA1 | 8 | Wierenga and Wadman (2003) |
| | D | 186 | 71 | 25.1 | ms | W rat | 14-28 d | - | CA1 | 8 | Wierenga and Wadman (2003) |
| | F | 129 | 68 | 24.04 | ms | W rat | 14-28 d | - | CA1 | 8 | Wierenga and Wadman (2003) |

Table S17: Schaffer collaterals physiologyexperimental data for SC→Exc synapses. UoM: Units of Measurement, R.: region, n.: number of cells.

| SC→Inh | Experimental Feature | Value (Mean) | SD | SEM | UoM | Species[1] | Age | R. | n. | Reference |
|---|---|---|---|---|---|---|---|---|---|---|
| PSC Magnitude | PSCs ratios SC-INH/SC-PC | 1.09 | 1.14 | 0.29 | - | W rat | 4-6 w | CA1 | 16 | Glickfeld and Scanziani (2006) |
| | PSCs ratios SC-INH/SC-PC | 8.15 | 6 | 1.5 | - | W rat | 4-6 w | CA1 | 16 | Glickfeld and Scanziani (2006) |
| PSP Kinetics | EPSP-IPSP latency | 1.9 | 0.6 | 0.2 | ms | W rat | 10 d | CA1 | 9 | Pouille and Scanziani (2001) |
| NMDA Kinetics | NMDA/AMPA ratio | 0.16 | 0.07 | 0.02 | - | Mouse | 17-23 d | CA1 | 12 | Le Roux et al. (2013) |
| | NMDA tau rise | 2.93 | - | - | ms | SD rat | 6-12 w | CA1 | 52 | Andrásfalvy and Magee (2001) |
| | NMDA tau decay | 154.6 | 81.9 | 25.9 | ms | Mouse | 14-24 d | CA1 | 10 | Cornford et al. (2019) |
| Short term plasticity | U | 0.11 | 0.04 | 0.01 | - | W rat | 14-28 d | CA1 | 20 | Wierenga and Wadman (2003) |
| | D | 307 | 233 | 52.1 | ms | W rat | 14-28 d | CA1 | 20 | Wierenga and Wadman (2003) |
| | F | 195 | 134 | 29.96 | ms | W rat | 14-28 d | CA1 | 20 | Wierenga and Wadman (2003) |

Table S18: Schaffer collaterals physiology experimental data for SC→Inh synapses. UoM: Units of Measurement, R.: region, n.: number of cells.

| Resting Membrane Potential (RMP) | | | | | | | | | | | | | |
|---|---|---|---|---|---|---|---|---|---|---|---|---|---|
| Neuron Type | Dose (μM) | Drug[7] | Application | Region | Layer | Species[1] | Age Weight | Vm ctr (mV) | Vm ACh (mV) | ΔVm (mV) | Current (nA) | N. cells | Reference |
| PC | 10 | CCh | focal | CA1 | SP | Mouse | Adult | -79 | -75.4 | 3.6 | 0.12 | 13 | Dasari and Gulledge (2011) |
| FSBC (PVBC) | 5 | CCh | bath | CA3 | SP | Mouse | 15-23 d | -65 | -58.9 | 6.1 | 0.17 | 8 | Szabó et al. (2010) |
| AAC | 5 | CCh | bath | CA3 | SP | Mouse | 15-23 d | -65 | -61.4 | 3.6 | 0.07 | 11 | Szabó et al. (2010) |
| RSBC | 5 | CCh | bath | CA3 | SP | Mouse | 15-23 d | -65 | -58.7 | 6.3 | 0.13 | 7 | Szabó et al. (2010) |
| INTs | 10 | CCh ACh | focal | CA1 | all | Mouse | 18-25 d | -70 | -55.5 | 14.5 | 0.37 | 102 | McQuiston and Madison (1999) |
| PC | 3 | CCh | bath | CA1 | SP | SD rat | Adult | -66 | -61 | 5 | 0.16 | 4 | Sheridan and Sutor (1990) |

| | | | | | | | | | | | | | |
|---|---|---|---|---|---|---|---|---|---|---|---|---|---|
| PC | 10 | CCh | bath | CA1 | SP | Mouse | 10-40 d | -61.3 | -56 | 5.3 | 0.17 | 12 | Palacios-Filardo et al. (2021) |
| PC | 1 | CCh | bath | CA1 | SP | W rat | 13-15 d | -75 | -72.4 | 2.6 | 0.09 | 8 | Buchanan et al. (2010) |
| PC | 50 | CCh | TBC | CA1 | SP | SD rat | 11-28 d | -72 | -65 | 7 | 0.23 | 9 | J. H. Williams and Kauer (1997) |
| PC | 100 | CCh | bath | SUB | all | SD rat | 200 300 g | -62.2 | -53.2 | 9 | 0.3 | 9 | Kawasaki and Avoli (1996) |

Firing Rate (FR)

| Neuron Type | Dose (μM) | Drug[7] | Application | Region | Layer | Species[1] | Age Weight | FR ctr (Hz) | FR ACh (Hz) | ΔFR (Hz) | Current (nA) | N. cells | Reference |
|---|---|---|---|---|---|---|---|---|---|---|---|---|---|
| OLM | 10 | Musc ACh | bath | CA1 | SO | Mouse | 14-21 d | 20 | 25 | 5 | 0.02 | 43 | Lawrence et al. (2006) |
| Other ADP | 10 | Musc ACh | bath | CA1 | SO | Mouse | 14-21 d | 20 | 37 | 17 | 0.12 | 43 | Lawrence et al. (2006) |

| | | | | | | | | | | | | | |
|---|---|---|---|---|---|---|---|---|---|---|---|---|---|
| PC | 3 | CCh | bath | CA1 | SP | SD rat | Adult | 6 | 9 | 3 | 0.06 | 4 | Sheridan and Sutor (1990) |
| CCKSCA | 10 | Musc | bath | CA1 | SR | Mouse | 15-20 d | 12.7 | 32 | 19.3 | 0.07 | 21 | Cea-del Rio et al. (2011) |

Table S19: Curated dataset on neuronal excitability changes caused by cholinergic modulation.

[7] ACh: Acetylcholine, CCh: Carbachol, Musc: Muscarine

| From | To | Dose (μM) | Drug[7] | Application | Region | Layer | Species[1] | Age<br>Weight | PSP<br>PSC | Ratio<br>ACh/ctr | N.<br>conn | Reference |
|---|---|---|---|---|---|---|---|---|---|---|---|---|
| SC | PC | 10 | CCh | focal | CA1 | SP | Mouse | Adult | EPSP | 38% | 9 | Dasari and Gulledge (2011) |
| PC/SC | PC | 0.01 | CCh | bath | CA1 | SLM<br>SR | SD rat | 5-10 w | EPSP | 100% | 5 | M. Hasselmo and Schnell (1994) |
| PC/SC | PC | 0.1 | CCh | bath | CA1 | SLM<br>SR | SD rat | 5-10 w | EPSP | 96% | 5 | M. Hasselmo and Schnell (1994) |
| PC/SC | PC | 1 | CCh | bath | CA1 | SLM<br>SR | SD rat | 5-10 w | EPSP | 81% | 5 | M. Hasselmo and Schnell (1994) |
| PC/SC | PC | 10 | CCh | bath | CA1 | SLM<br>SR | SD rat | 5-10 w | EPSP | 74% | 5 | M. Hasselmo and Schnell (1994) |
| PC/SC | PC | 100 | CCh | bath | CA1 | SLM<br>SR | SD rat | 5-10 w | EPSP | 47% | 13 | M. Hasselmo and Schnell (1994) |
| PC/SC | PC | 500 | CCh | bath | CA1 | SLM<br>SR | SD rat | 5-10 w | EPSP | 0% | 5 | M. Hasselmo and Schnell (1994) |
| PC/SC | PC | 0.01 | CCh | bath | CA1 | SLM<br>SR | SD rat | 5-10 w | EPSP | 100% | 5 | M. Hasselmo and Schnell (1994) |
| PC/SC | PC | 0.1 | CCh | bath | CA1 | SLM<br>SR | SD rat | 5-10 w | EPSP | 98% | 5 | M. Hasselmo and Schnell (1994) |
| PC/SC | PC | 1 | CCh | bath | CA1 | SLM<br>SR | SD rat | 5-10 w | EPSP | 67% | 5 | M. Hasselmo and Schnell (1994) |
| PC/SC | PC | 10 | CCh | bath | CA1 | SLM<br>SR | SD rat | 5-10 w | EPSP | 43% | 5 | M. Hasselmo and Schnell (1994) |
| PC/SC | PC | 100 | CCh | bath | CA1 | SLM<br>SR | SD rat | 5-10 w | EPSP | 13% | 13 | M. Hasselmo and Schnell (1994) |

| | | | | | | | | | | | | |
|---|---|---|---|---|---|---|---|---|---|---|---|---|
| PC/SC | PC | 500 | CCh | bath | CA1 | SLM<br>SR | SD rat | 5-10 w | EPSP | 0% | 5 | M. Hasselmo and Schnell (1994) |
| MF | PC | 1 | Musc | bath | CA3 | - | Rat | 100<br>200 g | EPSP | 100% | 19 | S. Williams and Johnston (1990) |
| MF | PC | 1 | Musc | bath | CA3 | - | Rat | 100<br>200 g | EPSC | 89% | 14 | S. Williams and Johnston (1990) |
| MF | PC | 10 | Musc | bath | CA3 | - | Rat | 100<br>200 g | EPSP | 77% | 7 | S. Williams and Johnston (1990) |
| MF | PC | 10 | Musc | bath | CA3 | - | Rat | 100<br>200 g | EPSC | 66% | 7 | S. Williams and Johnston (1990) |
| FSBC | PC | 5 | CCh | bath | CA3 | SP | Mouse | 15-23 d | IPSC | 30% | 16 | Szabó et al. (2010) |
| AAC | PC | 5 | CCh | bath | CA3 | SP | Mouse | 15-23 d | IPSC | 27% | 16 | Szabó et al. (2010) |
| RSBC | PC | 5 | CCh | bath | CA3 | SP | Mouse | 15-23 d | IPSC | 6% | 13 | Szabó et al. (2010) |
| SC | PC | 5 | CCh | bath | CA1 | SP | W rat | 14-18 d | EPSC | 32% | 11 | Sevilla et al. (2002) |
| SCC | PC | 0.1 | CCh | bath | CA1 | SP | SD rat | Adult | EPSP | 96% | 12 | Sheridan and Sutor (1990) |
| SCC | PC | 0.3 | CCh | bath | CA1 | SP | SD rat | Adult | EPSP | 83% | 12 | Sheridan and Sutor (1990) |
| SCC | PC | 1 | CCh | bath | CA1 | SP | SD rat | Adult | EPSP | 60% | 12 | Sheridan and Sutor (1990) |
| SCC | PC | 3 | CCh | bath | CA1 | SP | SD rat | Adult | EPSP | 30% | 12 | Sheridan and Sutor (1990) |
| SCC | PC | 10 | CCh | bath | CA1 | SP | SD rat | Adult | EPSP | 6% | 12 | Sheridan and Sutor (1990) |

Table S20: Curated dataset on synaptic transmission changes caused by cholinergic modulation.

| Species[1] | Age Weight | Dose (µM) | Drug[7] [8] | Region | Layer | Slice Thickness (µm) | ACSF (mM) | | | Measurement | Effects | N. | Reference |
|---|---|---|---|---|---|---|---|---|---|---|---|---|---|
| | | | | | | | Ca | Mg | K | | | | |
| G pig | 2-3 m | 50 | CCh | CA1 and CA3 | SP | 500 | 2 | 1.6 | 5 | Intracellular recording of PC | CCh-induced rhythmic bursts were recorded in both CA3 and CA1 PC of intact hippocampal slices. Where the CA3 and CA1 regions had been separated by a razor blade cut, rhythmic bursts were still observed in CA3 but not in CA1 neurons. | 5 | Bianchi and Wong (1994) |
| SD rat | 11-28 d | 50 | CCh | CA1 and CA3 | SR | 400 | 2.5 | 1.3 | 2.5 | Extracellular recording in SR | After a cut between CA3 and CA1, CCh induced robust oscillations in CA3, but not in CA1. | 4 | J. H. Williams and Kauer (1997) |
| W rat | 15-25 d | 20 | CCh | CA1 and CA3 | SR | 450 | 2 | 2 | 3 | Extracellular recording in SR | CCh-induced 40-Hz oscillations are generated within the CA3 area and propagate to CA1 in intact slice. After the cut between CA3 and CA1, oscillations in CA3 persisted, whereas activity was observed in neither CA1 nor dentate gyrus. | 4 | Fisahn et al. (1998) |
| SD rat | 10-30 d | 4-13 | CCh | CA1 and CA3 | SP | 400 | 2 | 2 | 5 | Extracellular recording in SP | Low concentrations of CCh (4-13 µM ) produced a regular pattern of synchronous discharges in the delta range (0.5-2 Hz). This rhythmic pattern originated in CA3, as assessed by isolating CA3 from CA1. | 16 | Fellous and Sejnowski (2000) |
| SD rat | 10-30 d | 13-60 | CCh | CA1 and CA3 | SP | 400 | 2 | 2 | 5 | Extracellular recording in SP | Higher concentrations of CCh (13-60 µM ) produced short episodes of synchronous population discharges at a regular frequency (5-10 Hz). Separating CA1 from CA3, oscillations were found in isolated CA3, but not in isolated CA1, and dual field recordings in CA3 and CA1 revealed a positive latency ( 10-15 ms) in CA1. | 3 | Fellous and Sejnowski (2000) |
| SD rat | 10-30 d | 8-25 | CCh | CA1 and CA3 | SP | 400 | 2 | 2 | 5 | Extracellular recording in SP | Concentrations of CCh in the 8-25 µM range produced faster population discharges (40-50 Hz) in the gamma band in both CA1 and CA3. This oscillation was present in CA3 minislices, but was never observed in CA1 minislices. | 6 | Fellous and Sejnowski (2000) |
| SD rat | 200-300 g | 1-100 | CCh | CA1 | all | 400 | 2 | 1 | 3 | Extracellular recordings | The power spectra show a single distinguishable power peak in the gamma frequency band (45 Hz) in CA1 mini-slice. Maximal $\gamma$ power was recorded at the SO-SP border. | 78 | Pietersen et al. (2014) |
| SD rat | 200-300 g | 10 | CCh | CA1 and CA3 | SP | 400 | 2 | 1 | 3 | Extracellular recording in SP | In intact hippocampal slices CCh induced $\gamma$ in both area CA3 and CA1.The $\gamma$ in CA1a was phase-locked to the slow $\gamma$ in CA3b. In these slices the power spectrum in CA1a had a single peak in the gamma frequency band with a dominant frequency of 33 Hz. | 53 | Pietersen et al. (2014) |
| LE rat | 3-8 m 350-500 g | - | Tail pinches (correlated with ACh release) | CA1 | all | - | - | - | - | Extracellular recordings | ACh release was highly correlated with the appearance of both spontaneous and induced theta oscillations (such release lagged behind theta initiation by 25-60 s) | - | Zhang et al. (2010) |
| ChAT-Cre mouse | 2-6 m | - | Optogenetic stimulation | CA1 and CA3 | all | - | - | - | - | Extracellular recordings | Cholinergic stimulation completely blocked sharp wave ripples and strongly suppressed the power of both slow oscillations (0.5–2 Hz in anesthetized, 0.5–4 Hz in behaving animals) and supratheta (6–10 Hz in anesthetized, 10–25 Hz in behaving animals) bands. The same stimulation robustly increased both the power and coherence of theta oscillations (2–6 Hz) in urethane-anesthetized mice | - | Vandecasteele et al. (2014) |

Table S21: Network effects of ACh.

[8] All bath application except for Bianchi and Wong (1994) that used bath and focal application.

| Pre Mtype | Post Mtype | Rat Age | [Ca2+] | Type | n | Frequency | | Reference | Notes |
|---|---|---|---|---|---|---|---|---|---|
| | | | | | | Mean | SEM | | |
| Excitatory | Pyramidal | 21-32 D | 2,5 | mEPSC | 8 | 3.120 | 0.320 | Ito and Schuman (2009) | |
| Inhibitory | Pyramidal | 20-28 D | 2 | sIPSC | 3 | 5.567 | 2.541 | Hájos and Mody (1997) | KYNA blocks AMPA/NMDA |
| Inhibitory | Inhibitory | 20-28 D | | sIPSC | 24 | 2.433 | 0.309 | Hájos and Mody (1997) | |
| Excitatory | Pyramidal | | 2 | sEPSC | 16 | 0.660 | 0.260 | Esclapez et al. (1999a) | Soma |
| Excitatory | Pyramidal | | 2 | mEPSC | 5 | 0.130 | 0.040 | Esclapez et al. (1999a) | Soma + TTX |
| Excitatory | Pyramidal | | 2 | sEPSC | 13 | 1.790 | 0.370 | Esclapez et al. (1999a) | Dendrites |
| Excitatory | Pyramidal | | 2 | mEPSC | 11 | 0.300 | 0.030 | Esclapez et al. (1999a) | Dendrites + TTX |
| Excitatory | Pyramidal | 1.5-6 M | 2 | sEPSC | 8 | 0.530 | 0.160 | Cossart et al. (2000) | Soma in SP |
| Excitatory | Pyramidal | 1.5-6 M | 2 | mEPSC | 8 | 0.020 | 0.010 | Cossart et al. (2000) | Soma in SP + TTX |
| Excitatory | Pyramidal | 1.5-6 M | 2 | sEPSC | 9 | 2.010 | 0.420 | Cossart et al. (2000) | Dendrites in SR |
| Excitatory | Pyramidal | 1.5-6 M | 2 | mEPSC | 9 | 0.480 | 0.100 | Cossart et al. (2000) | Dendrites in SR + TTX |
| Excitatory | Pyramidal | 1.5-6 M | 2 | sEPSC | 6 | 0.430 | 0.100 | Cossart et al. (2000) | Dendrites in SO |
| Excitatory | Pyramidal | 1.5-6 M | 2 | mEPSC | 6 | 0.040 | 0.020 | Cossart et al. (2000) | Dendrites in SO + TTX |
| Excitatory | Pyramidal | 1.5-6 M | 2 | sEPSC | 5 | 1.960 | 0.560 | Cossart et al. (2000) | Soma in SO |
| Excitatory | Pyramidal | 1.5-6 M | 2 | mEPSC | 5 | 0.150 | 0.070 | Cossart et al. (2000) | Soma in SO + TTX |
| Inhibitory | Pyramidal | 1.5-6 M | 2 | sIPSC | 8 | 16.820 | 2.210 | Cossart et al. (2000) | Soma in SP |
| Inhibitory | Pyramidal | 2.5-6 M | 2 | mIPSC | 8 | 13.040 | 4.700 | Cossart et al. (2000) | Soma in SP + TTX |
| Inhibitory | Pyramidal | 3.5-6 M | 2 | sIPSC | 9 | 17.290 | 2.300 | Cossart et al. (2000) | Dendrites in SR |
| Inhibitory | Pyramidal | 4.5-6 M | 2 | mIPSC | 9 | 1.980 | 0.580 | Cossart et al. (2000) | Dendrites in SR + TTX |
| Inhibitory | Pyramidal | 5.5-6 M | 2 | sIPSC | 6 | 21.150 | 3.990 | Cossart et al. (2000) | Dendrites in SO |
| Inhibitory | Pyramidal | 6.5-6 M | 2 | mIPSC | 6 | 3.650 | 0.630 | Cossart et al. (2000) | Dendrites in SO + TTX |
| Inhibitory | Pyramidal | 7.5-6 M | 2 | sIPSC | 5 | 32.090 | 7.780 | Cossart et al. (2000) | Soma in SO |
| Inhibitory | Pyramidal | 8.5-6 M | 2 | mIPSC | 5 | 11.410 | 2.060 | Cossart et al. (2000) | Soma in SO + TTX |

Table S22: Summary of estimated rates of spontaneous synaptic release in rat CA1. TTX used to inactivate fast sodium channels in all experiments except Hájos and Mody (1997). D:Days, M:Months.

| Mtype | Mean angle (deg) | Angular deviation (deg) | Mean firing rate (Hz) | Min firing rate (Hz) | Max firing rate (Hz) | N. | Species[1] | Reference |
|---|---|---|---|---|---|---|---|---|
| SP_PVBC | 271 | 68 | 7.3 | 3.4 | 17.6 | 5 | rat | Klausberger et al. (2003) |
| SP_AA | 185 | 55 | 17.1 | 9.1 | 25.1 | 2 | rat | Klausberger et al. (2003) |
| SP_PC | 20 | 65 | - | - | - | 6 | rat | Klausberger et al. (2003) |
| SO_OLM | 19 | 57 | 4.9 | 2.5 | 6.1 | 3 | rat | Klausberger et al. (2003) |
| SP_BS | 1 | 60 | 5.9 | 0.5 | 21.7 | 5 | rat | Klausberger et al. (2004) |
| SP_CCKBC | 173 | - | 9.4 | 4.6 | 12.7 | 4 | rat | Klausberger (2005) |
| SP_Ivy | 30.7 | 63.1 | 4.2 | - | - | 4 | rat | Fuentealba et al. (2008) |

Table S23: Phase tuning of rat CA1 morphological type.

| Neuron Type | Mean rate (Hz) | SE rate (Hz) | N. | SD rate (Hz) | Recording condition | Source |
|---|---|---|---|---|---|---|
| Pyramidal cells | 1.4 | 0.1 | 246 | 1.6 | in vivo, behaving | Csicsvari et al. (1999) |
| SP interneurons | 16.3 | 1.52 | 55 | 11.3 | in vivo, behaving | Csicsvari et al. (1999) |
| Alveus/Oriens (a/o) interneurons | 11.9 | 1.5 | 68 | 12.4 | in vivo, behaving | Csicsvari et al. (1999) |

Table S24: Long-term discharge rates of rat CA1 neurons in vivo during theta periods.

| Campaign | Circuit Size | SC | Report On | Duration (s) | Time(s) | Max Mem per Node | N. of Nodes | Total Mem (GB) | Simulator |
|---|---|---|---|---|---|---|---|---|---|
| cav | cylinder | No | Soma+LFP | 10.0 | 20927.09 | 636.35 | 6 | 3.729 | N |
| cav | full | No | Soma+LFP | 10.0 | 15891.12 | 3201.16 | 300 | 937.84 | CN |
| cav | slice | No | Soma+LFP | 10.0 | 16195.65 | 603.95 | 16 | 9.437 | N |
| minis | cylinder | Yes | Soma+LFP | 10.0 | 34241.17 | 2574.45 | 16 | 40.226 | N |
| ms-input | cylinder | No | Soma+LFP | 20.0 | 6220.22 | 700.16 | 60 | 41.025 | N |
| sasaki | slice | Yes | Soma | 1.5 | 5318.58 | 2083.16 | 20 | 40.687 | N |
| sc-oscillatory | cylinder | Yes | Soma+LFP | 10.0 | 15403.8 | 1236.87 | 36 | 43.484 | N |
| sc-oscillatory | cylinder | Yes | Soma+LFP | 10.0 | 3772.47 | 1202.79 | 36 | 42.286 | CN |
| sc-oscillatory | full | Yes | Soma+LFP | 10.0 | 35914.36 | 5315.56 | 300 | 1557.293 | CN |
| sc-oscillatory | slice | Yes | Soma+LFP | 10.0 | 7598.11 | 1928.43 | 36 | 67.796 | CN |
| sc-unstructured | cylinder | Yes | Soma+LFP | 10.0 | 17444.86 | 1448.93 | 36 | 50.939 | N |
| zemankovich | slice | Yes | Soma+LFP | 5.0 | 15888.6 | 2208.81 | 36 | 77.653 | N |

Table S25: Resources and simulators used in validating the current CA1 model. cav: calcium-voltage scan. osc control: oscillatory; ms: medial septum, SC: Schaffer collateral; N: Neuron; CN: CoreNeuron.

### S1 Author contribution

#### Author contributions

H.M. conceived and led the study. S.K., M.M., A.M.T., F.S., E.M., A.M., and T.F. co-led the study. A.M.T. and A.M. planned and supervised experiments. A.R., J.B., A.A., D.B., K.K. planned and supervised data integration, strategies and algorithms, model building, simulation experiments, and analysis. K.K. and A.R. built the CA1 circuit and implemented version control. A.A., J.B., D.B., A.R. reconstructed Schaffer collaterals. A.A., C.C., J.B., A.R. modeled acetylcholine. J.B. and A.R. worked on theta. A.A., J.B. and A.R. worked on oscillation propagation. F.S. and J-D.C. planned and supervised the development of algorithms, software and workflows, computing infrastructure, and technical integration. A.R., J.B., A.A., D.B., K.K., C.C. wrote the manuscript. A more detailed listing of author contributions is shown below.

##### Manuscript

A.R., J.B., A.A., D.B., K.K, and C.C. wrote the manuscript. S.S., E.G., L.K., N.R.G. contributed to the writing of the manuscript. E.B., J.B., A.A., A.R., and K.K. generated the figures. D.B. contributed to the generation of figures. J.B. compiled and integrated all the references. All authors provided input to the manuscript.

##### Data curation

M.F.S. and A.R. co-supervised and data integration. A-K.K., A.R., A.A., and J.B. performed data integration.

##### Modeling

E.M., S.K., M.M., A.R., and H.M. co-supervised model building and analysis. M.M., R.M., C.A.L., L.V. and W.V.G. performed modeling of channels and cells. T.D., H.L., and A.R.: created Atlas and coordinate system. A.R. performed neuronal composition modeling. J.L.R. and M.G. implemented the voxelized cell placement algorithms. A.R., A.E., and M.R. performed modeling of anatomical connections. A.A. and A.E. performed modeling of synapses. A.R., K.K. and V.S. built the CA1 circuit. A.Ar. computed single cell rheobase J.H. and M.G. implemented the projection algorithm. A.A., J.B., D.B., A.R. reconstructed Schaffer collaterals. A.A., C.C., J.B., A.R. modeled acetylcholine.

##### Validation

L.K. and K.K. performed clone validation. N.R.G., J.B.H., F.P., and C.F. morphology collage. S.S. performed validation of the single cell models. A.R. and K.K. performed validation of the cell compositions. A.R. and K.K. performed validation of anatomical connections. A.A. and A.E. performed validation of synapses. A.A., J.B. and D.B. performed validation of the Schaffer collaterals. S.S. and

J.B. performed validation of gamma propagation (Zemancovic). A.R., K.K. and A.A. performed the SC input-output(I/O) validation (Sasaki). P.R., J.B., and A.A. performed statistical analyses.

#### Simulations

H.M. supervised simulations. J.B. and A.R. worked on theta oscillations. A.A., J.B. and A.R. worked on oscillation propagation. A.R., K.K. and A.A. worked on SC input-output relationship

#### Software development

J.H. and M.G. developed the algorithm to reconstruct Schaffer collaterals. L.V.H and W.V.G. developed algorithmic changes and scripts for BluePyOpt and BluePyMM. G.I. automation of simulation campaigns.

#### Visualizations

J.P., A.R., and H.M. supervised scientific visualizations and movies E.B., N.R.G., M.A., and F.P. generated scientific visualizations. C.F. generated movies.

#### Computing infrastructure

J-D.C and J.G.K. supervised the assurance, efficiency, and maintenance of adequate high performance computing capacity. P.K. supported infrastructure for simulation and M.G. provided engineering support.

#### Experiments

A.M.T. and A.M. performed paired recordings.

Y.S., A.M, S.L and J.F reconstructed neurons.

INITIALS: A.A. Alberto Antonietti, A.Ar. Alexis Arnaudon, A.E. András Ecker, A.M. Audrey Mercer, A.R. Armando Romani, A.M.T. Alex M. Thomson, A-K.K. Anna-Kristin Kaufmann, C.A.L. Carmen Alina Lupascu, C.C. Cristina Colangelo, C.F. Cyrille Favreau, D.B. Davide Bella, E.B. Elvis Boci, E.G. Elisabetta Giacalone, E.M. Eilif Muller, F.S. Felix Schürmann, G.I. Genrich Ivaska, H.L. Huanxiang Lu, H.M. Henry Markram, J.B. Julian Budd, J.B.H. Juan B. Hernando, J.F. Joanne Falck, J.H. Joni Herttuainen, J.G.K. James Gonzalo King, J.L.R. Juan Luis Riquelme, F.P. Fabien Petitjean, J.P. Judit Planas, J-D.C. Jean-Denis Courcol, K.K. Kerem Kurban, L.K. Lida Kanari, L.V. Liesbeth Vanherpe, M.A. Marwan Abdellah, M.F.S. Mohameth François Sy, M.G. Michael Gevaert, M.M. Michele Migliore, M.W.R. Michael W. Reimann, N.R.G. Nadir Román Guerrero, P.K. Pramod Kumbhar, P.R. Pranav Rai, R.M. Rosanna Migliore, S.K. Szabolcs Káli, S.L. Sigrun Lange, S.R. Srikanth Ramaswamy, S.S. Sára Sáray, T.D. Thomas Delemontex, T.F. Tamás F. Freund, Y.S. Ying Shi, V.S. Vishal Sood, W.V.G. Werner Van Geit

#### Acknowledgments

The authors would like to thank all the people involved in the rat CA1 hippocampus project over the last years, in particular Attila Guylás, Luc Guyot, and Arseny Povolotsky. We would also like to thank those who offered valuable advice or data: Giorgio Ascoli, Norbert Ha jos, Jesse Jackson, Corette Wierenga, Sylvain Williams. Experimentalists who generated data for this project or who recorded, dye-filled and/or reconstructed neurones: A.B. Ali, J. Deuchars, A..P Bannister, R. Begum, N. Botcher, J. Deuchars, K. Eastlake, D. I. Hughes, M. Ilia, J. Kerkhoff, S. Kirchhecker, H. Pawelzik, and H. Trigg. D.C. West designed data collection and analysis software. We thank those who worked on the Explore feature of the Hippocampus hub: Anil Tuncel, Pavlo Getta, Caitlin Monney, Stefano Antonel, Alexander Dietz, and Liviu Soltuzu. Finally, we are indebted to Karin Holm for copy editing and publication advice and support.
